# Targeting of retrovirus-derived *Rtl8a*/*8b* reduces social response and increases apathy-like behavior associated with GABRB2 reduction

**DOI:** 10.1101/2024.08.02.606341

**Authors:** Yoshifumi Fujioka, Hirosuke Shiura, Masayuki Ishii, Ryuichi Ono, Tsutomu Endo, Hiroshi Kiyonari, Yoshikazu Hirate, Hikaru Ito, Masami Kanai-Azuma, Takashi Kohda, Tomoko Kaneko-Ishino, Fumitoshi Ishino

**Author notes:** Correspondence to: Fumitoshi Ishino,. Laboratory for Functional Non-coding Genomics, RIKEN Center for Integrative Medical Sciences (IMS), Kanagawa 230-0045, Japan. Department of Animal Resource Sciences, Graduate School of Agricultural and Life Sciences, the University of Tokyo, Tokyo 113-8657, Japan (Tsutomu Endo). Institute of Research, TMDU, Tokyo 113-8510, Japan.

## Abstract

Retrotransposon Gag-like (RTL) 8A, 8B and 8C are eutherian-specific genes derived from a certain retrovirus. They clustered as a triplet of genes on the X chromosome, but their function remains unknown. Here, we demonstrate that Rtl8a and Rtl8b play important roles in the brain: their double knockout (DKO) mice not only exhibit reduced social responses and increased apathy-like behavior, but also become obese from young adulthood. Mouse RTL8A/8B proteins are expressed in the prefrontal cortex and hypothalamus and localize to both the nucleus and cytoplasm of neurons, presumably due to the N-terminal nuclear localization signal-like sequence at the N-terminus. An RNAseq study in the cerebral cortex revealed reduced expression of several GABA type A receptor subunits, in particular the β2 subunit of Gabrb2, in DKO. We confirmed the reduction of GABRB2 protein in the DKO cerebral cortex by Western blotting. As GABRB2 has been implicated in the etiology of several neurodevelopmental disorders, it is likely that the reduction of GABRB2 is one of the major causes of the neuropsychiatric defects in the DKO mice.

## Introductions

In the human genome, there are 11 <u>r</u>etrotransposon Gag-like (RTL, also known as <u>s</u>ushi-ichi-<u>r</u>elated retrotransposon <u>h</u>omolog (SIRH)) genes that encode proteins homologous to the sushi-ichi GAG (and sometimes also to POL) protein(s) (Ono et al, 2001; Charlier et al, 2001; Brandt et al, 2005; Youngson et al, 2005; Ono et al, 2006). The Ty3/gypsy retrotransposon which includes the sushi-ichi retrotransposon, is an infectious retrovirus in *Drosophila melanogaster* (Kim et al, 1994; Song et al, 1994). Recently, it has been reported that RTL/SIRH genes were domesticated from three independent lineages of Ty3/gypsy (formally known as *Metaviridae*) (Henriques et al, 2024). Importantly, certain RTL/SIRH genes play essential and/or important roles in eutherian development (Kaneko-Ishino & Ishino, 2012, 2015, 2023; Imakawa et al, 2022; Shiura et al, 2023), such as paternally expressed 10 (*Peg10*), *Rtl1* (aka *Peg11*) and *Ldoc1* (aka *Rtl7* or *Sirh7*) in the placenta (Ono et al, 2006; Sekita et al, 2008; Naruse et al, 2014), *Rtl4* (aka *Sirh11* or *Zcchc16*) (Lim et al, 2013; Irie et al, 2015) and *Rtl1*(Kitazawa et al, 2021; Chou et al, 2022) in the brain, and *Rtl6* (aka *Sirh3*), *Rtl5* (aka *Sirh8*) and *Rtl9* (aka *Sirh10*) in the innate immune system of the brain against bacteria, viruses and fungi as microglial genes, respectively (Irie et al, 2022; Ishino et al, 2023). *Rtl1* is also important for muscle development (Kitazawa et al, 2020). The biological roles of *RTL8A, 8B* and *8C* (aka *SIRH5, 6* and *4*) remain unknown, although their involvement has recently been implicated in two human diseases, Angelman syndrome (AS) and amyotrophic lateral sclerosis (ALS) (Pandya et al, 2021; Whiteley et al, 2021). AS is a neurodevelopmental genomic imprinting disorder caused by paternal duplication of chromosome 15q11-q13 harboring maternally expressed *ubiquitin protein ligase E3A* (*UBE3A*) or *UBE3A* mutations (Nicholls & Knepper, 2001; Chamberlain & Lalande, 2010; Germain et al, 2020). Both RTL8A-C and PEG10 proteins, another RTL gene product, were increased in differentiated neurons from AS patient iPSCs, suggesting that both RTL8A-C and PEG10 proteins are direct targets of UBE3A (Pandya et al, 2021). Therefore, it is likely that RTL8A-C is decreased in Prader-Willi syndrome (PWS) which is caused by maternal duplication of the same chromosomal region associated with a double dose of UBE3A. PWS is another neurodevelopmental genomic imprinting disorder characterized by late-onset obesity and autism spectrum disorder (ASD)-like symptoms from the juvenile period (Nicholls & Knepper, 2001; Tauber & Hoybye, 2021). Whiteley *et al*. demonstrated that RTL8A-C decreased but PEG10 increased in the differentiated neurons of ALS patient iPSCs, suggesting that PEG10 is a direct target of ubiquitin-like protein (ubiquilin) 2, UBQLN2 (Whiteley et al, 2021), a major gene for responsible for ALS (Deng et al, 2011), and also that RTL8A-C is protected from proteasome-dependent degradation by complex formation with UBQLN2. It is also reported that RTL8 is required for the nuclear translocation of UBQLN2 and thus for quality control in the neuronal nuclei (Mohan et al, 2022).

Inhibitory neurotransmission is primarily mediated by γ-aminobutyric acid (GABA), which activates synaptic GABA type A (GABA_A_) receptors. In humans, GABA_A_ receptors are composed of pentameric subunits selected from eight different classes, such as α1-6, β1-3, γ1-3, δ, ε, π, θ, ρ1-3 (Sigel *et al*, 2012; Sente *et al*, 2022). The major isoform of GABA_A_ receptors in the brain is composed of two α1, two β2 and one γ2 subunits (McKernan & Whiting, 1996; Baumann et al, 2002). Importantly, several GABA_A_ receptor subunits have been shown to be associated with psychiatric disorders. For example, alterations in the GABA type A receptor subunit beta2 (*GABRB2*) have been implicated in several neuropsychiatric disorders, including schizophrenia (SZ), bipolar disorder (BP), frontotemporal dementia (FTD) and ASD (Lo et al, 2004; Petryshen et al, 2005; Ma et al, 2005; Lo et al, 2007; Chen et al, 2009; Yuan et al, 2015; Jiang et al, 2018; Satterstrom et al, 2020). *GABRB2* has also been reported to be associated with epilepsy, Alzheimer’s disease, substance dependence, depression, internet gaming disorder, and premenstrual dysphoric disorder (Barki & Xue, 2022).

In this work, we addressed the biological function of *Rtl8a* and *Rtl8b* using their double knockout (DKO) mice and demonstrated that they play an important role in neuropsychiatric behavior, possibly via *Gabrb2* downregulation. We also discuss the possible association of *RTL8* downregulation with PWS, since the phenotypes of the *Rtl8a* and *Rtl8b* DKO mice are quite similar to those of patients with late PWS, ASD-like behaviors as well as late-onset obesity.

## Results

### Conservation of *RTL8A-C* in eutherians

*RTL8A-C* (aka *SIRH5, 6* and *4*) forms a multi-gene cluster on the X chromosome. In eutherians, its number (2-4, mostly 3, excluding pseudogenes) and amino acid (aa) sequence are well conserved (Table S1, red: dN/dS <0.1, pink: 0.1 ≤ dN/dS < 0.2), suggesting its important role in eutherians. In most cases, *RTL8s* within the same species exhibit the highest homology to each other compared to those in other species, suggesting independent gene conversion events in each species, and therefore are not in a precise orthologous relationship among eutherians (Fig. S1).

In mice, *Rtl8a* and *Rtl8b* encode identical proteins with 113 aa, while *Rtl8c* encodes a protein with 112 aa exhibiting 70% aa homology to the former (Figs. 1A, B and Fig. S2). They share homology with the suchi-ichi retrotransposon GAG (48.7 and 47.4% similarity and 34.2 and 28.9% identity, respectively). Mouse RTL8A-C (mRTL8A-C) proteins share a high overall aa sequence homology with those of other eutherian species (Table S1, Fig. S2), but mRTL8A (also mRTL8B) has a rodent-specific nuclear localization signal (NLS)-like sequence (K-K/R-X-K/R) in the N-terminus (Kalderon et al, 1984), whereas mRTL8C has a unique leucine zipper motif (76-97 aa) (Figs. 1A, B and Fig. S2).

**Figure 1.**
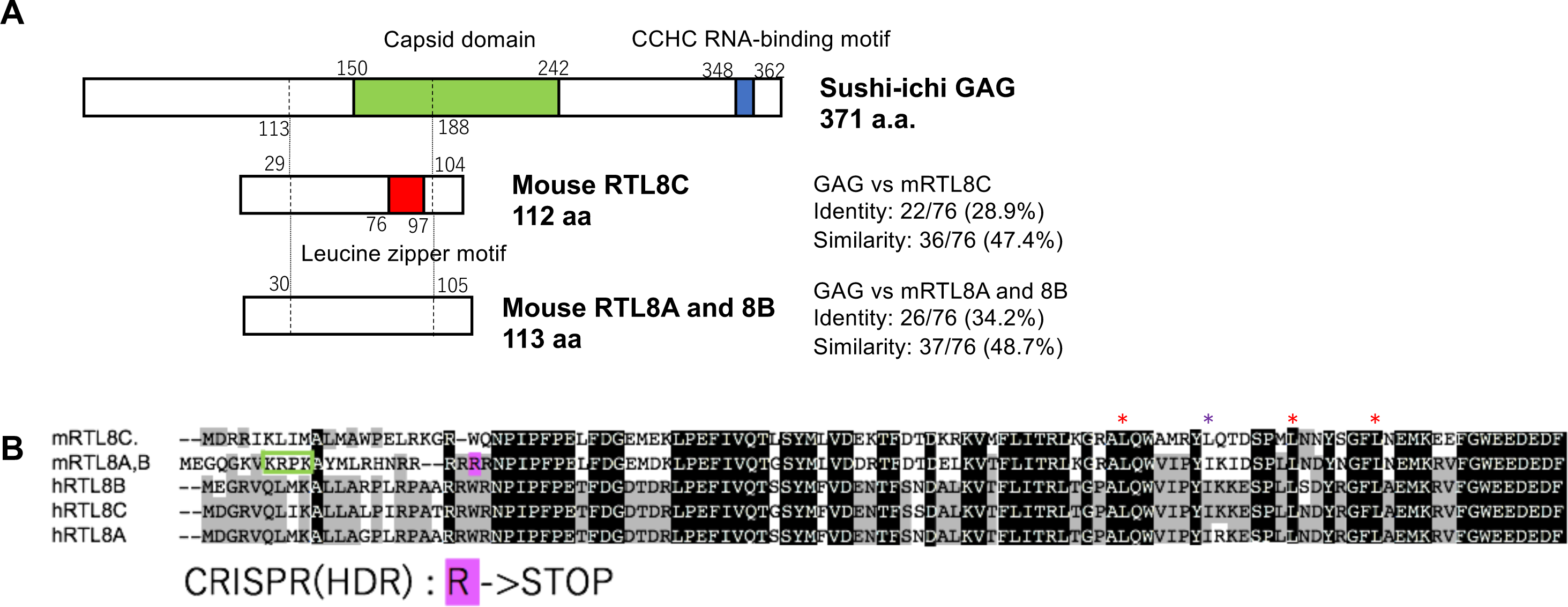
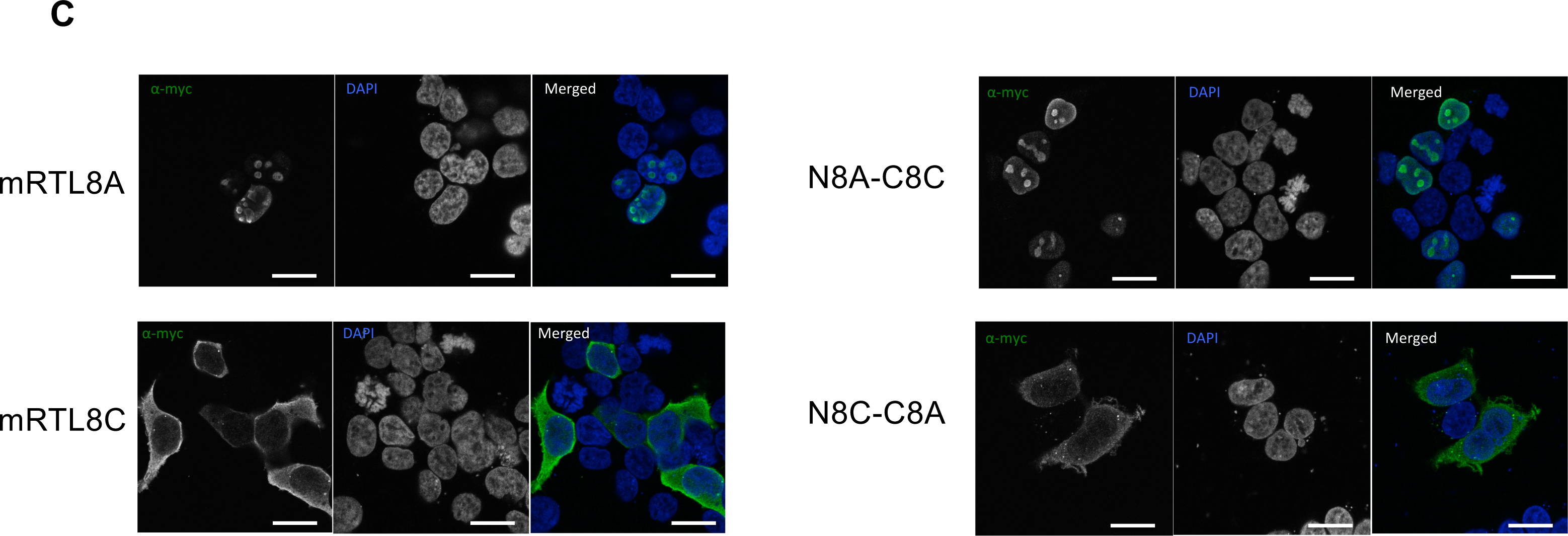
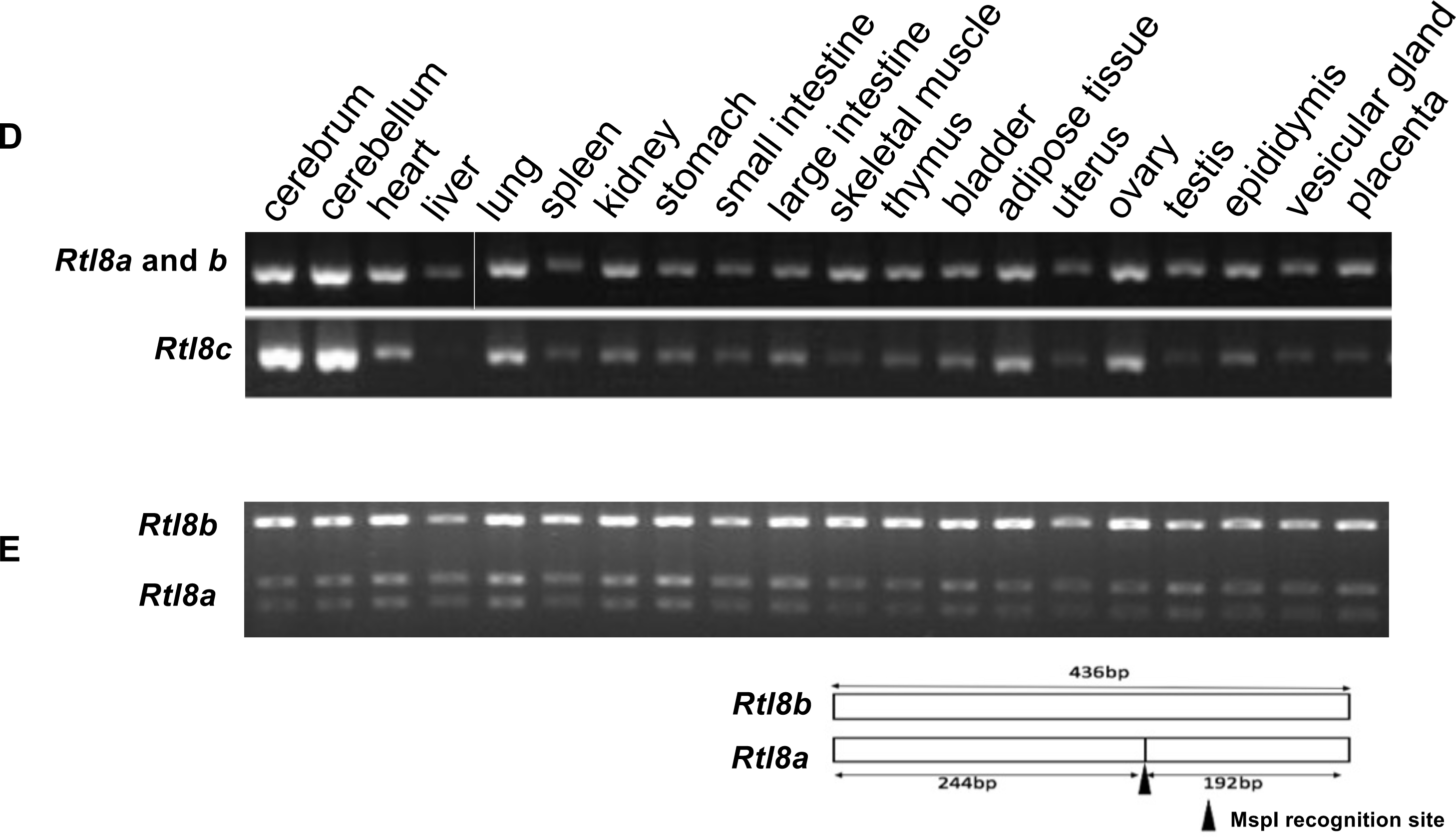
Characteristics of mouse RTL8A-C proteins. **(A)** Both mRTL8A and 8B comprise 113 aa with an identical aa sequence while mRTL8C comprises 112 aa including a unique leucine zipper motif (the red box) (see also Fig. S2). **(B**) Alignment of mouse and human RTL8A-C. The leucine zipper motif of mRTL8C is indicated by 4 asterisks (top). It should be noted that the second leucine (the purple asterisk) is unique to mRTL8C (see also Fig. S2). mRTL8A/8B has a rodent-specific nuclear localization signal (NLS)-like peptide in the N-terminus (the green box). A stop codon shown in pink was replaced with arginine (R) at 23 aa from the N-terminus in *Rtl8a* and *8b* DKO. **(C)** Subcellular localization of mRTL8A and mRTL8C proteins (left), and chimera proteins that have the N-half of mRTL8A and C-half of mRTL8C (N8A-C8C) and the N-half of RTL8C and C-half of RTL8A (N8C-C8A). Bar: 20 μm. **(D)** Expression profiles of *Rtl8a/8b* and *Rtl8c.* RT-PCR results in adult tissues and organs are shown. **(E)** RFLP analysis of *Rtl8a* and *8b*. The upper bands represent *Rtl8a* expression while the bottom two bands represent *Rtl8b* expression. The black arrowhead indicates the location of the recognition site of MspI.

We examined the subcellular localization of the mRTL8A and 8C proteins in NIH3T3 cells by plasmid overexpression: the former in the nuclei (nucleoli) and the latter in the cytoplasm (Fig. 1C, left). In addition, the subcellular localization of two chimeric proteins of the N- and C-terminal halves of mRTL8A and 8C, such as N8A-C8C and N8C-C8A, was determined by their N-terminus (Fig. 1C, right), suggesting that the rodent-specific NLS-like sequence of mRTL8A (and 8B) can function *in vivo*.

*Rtl8a*-*c* mRNAs exhibit higher expression in the cerebrum and cerebellum, but are ubiquitously expressed in all other tissues and organs, with the exception of no *Rtl8c* expression in the liver (Fig. 1D). *Rtl8a* and *Rtl8b* exhibit similar expression patterns as estimated by a DNA polymorphism analysis using an MspI recognition site (Fig. 1E). To address their biological functions, we first generated *Rtl8c* flox mice, and then both *Rtl8a* and *Rtl8b* were knocked out by introducing a stop codon at a site 23 aa from the N-terminus of each gene using CRISPR/Cas9 mediated genome editing (Figs. S3-S6).

### Reproductive and growth abnormalities in *Rtl8a*/*8b* DKO mice

*Rtl8a*/*Rtl8b* heterozygous (hetero) DKO female mice exhibited abnormal maternal care, and both hetero DKO females and hemizygous DKO males (hereafter referred to as DKO males) (Fig. S7) exhibited overgrowth from the young adult stage (Fig. 2). Unexpectedly, half of the hetero DKO females had delivery problems: most of their pups died shortly after birth, regardless of genotype (Fig. S8, left). In the case of homozygous (homo) DKO mothers, no pups survived (Fig. S8, right). However, no fetal and postnatal lethality was observed in the DKO pups when their fertilized eggs were transferred into pseudopregnant ICR females and cared for by the ICR mothers. These results suggest abnormal maternal behavior of the hetero and homo DKO mothers. It is likely that the hetero DKO females may exhibit either normal or abnormal maternal behavior due to random X chromosome inactivation (XCI) in female mice (Lyon, 1961; Takagi & Sasaki, 1975).

**Figure 2.**
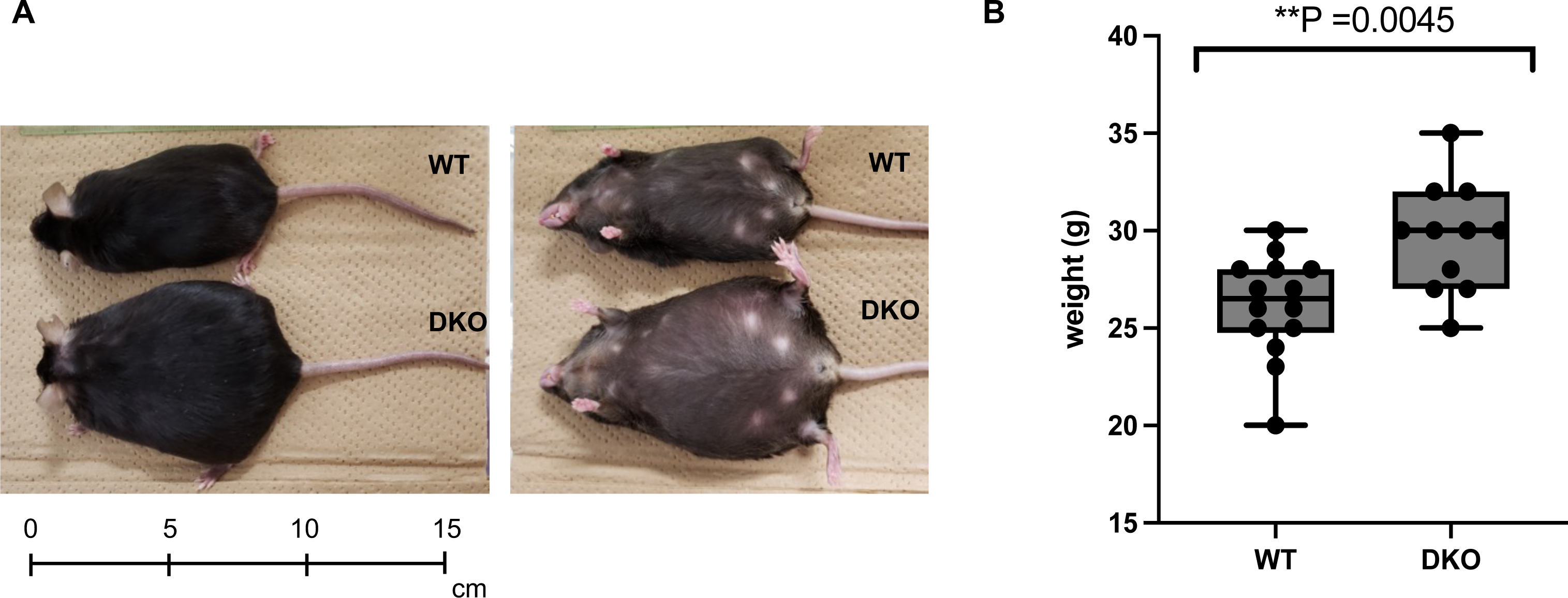
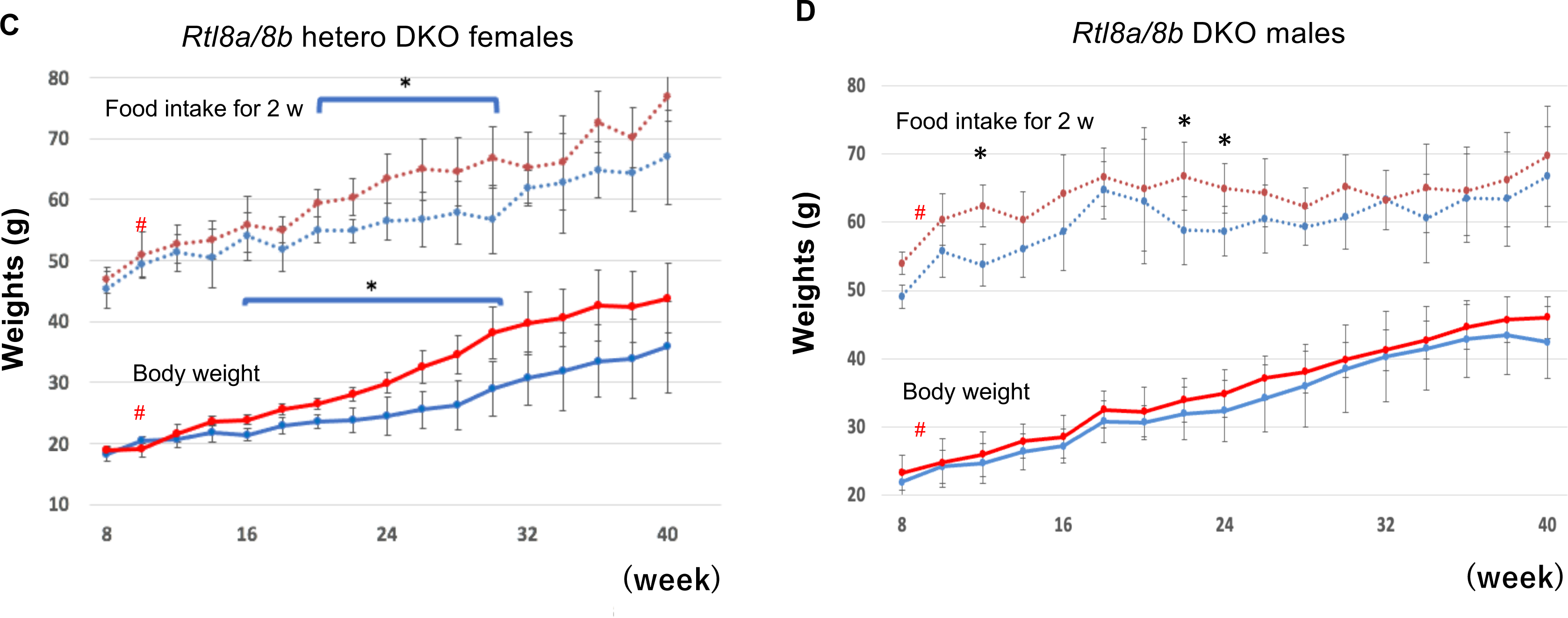
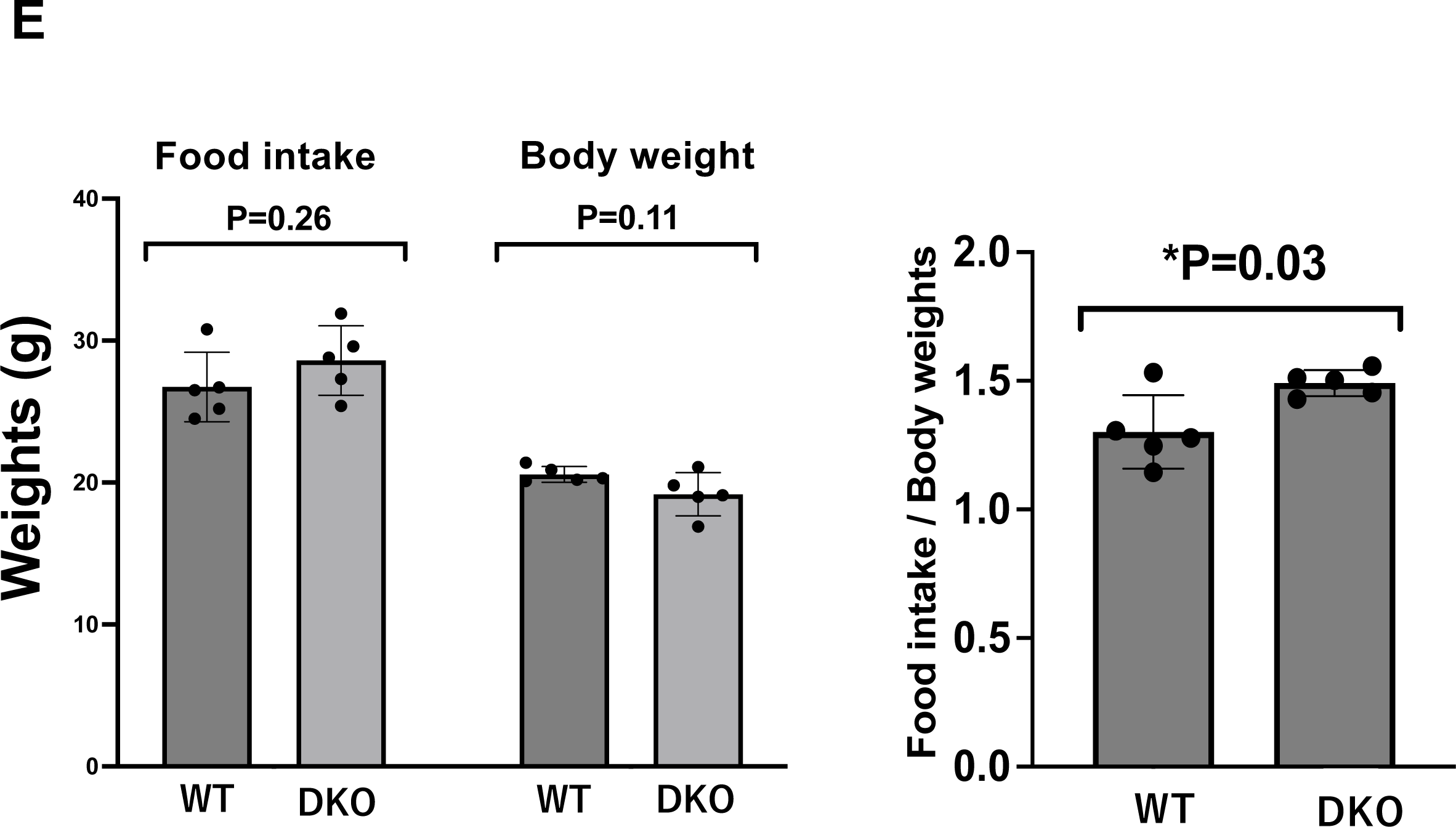
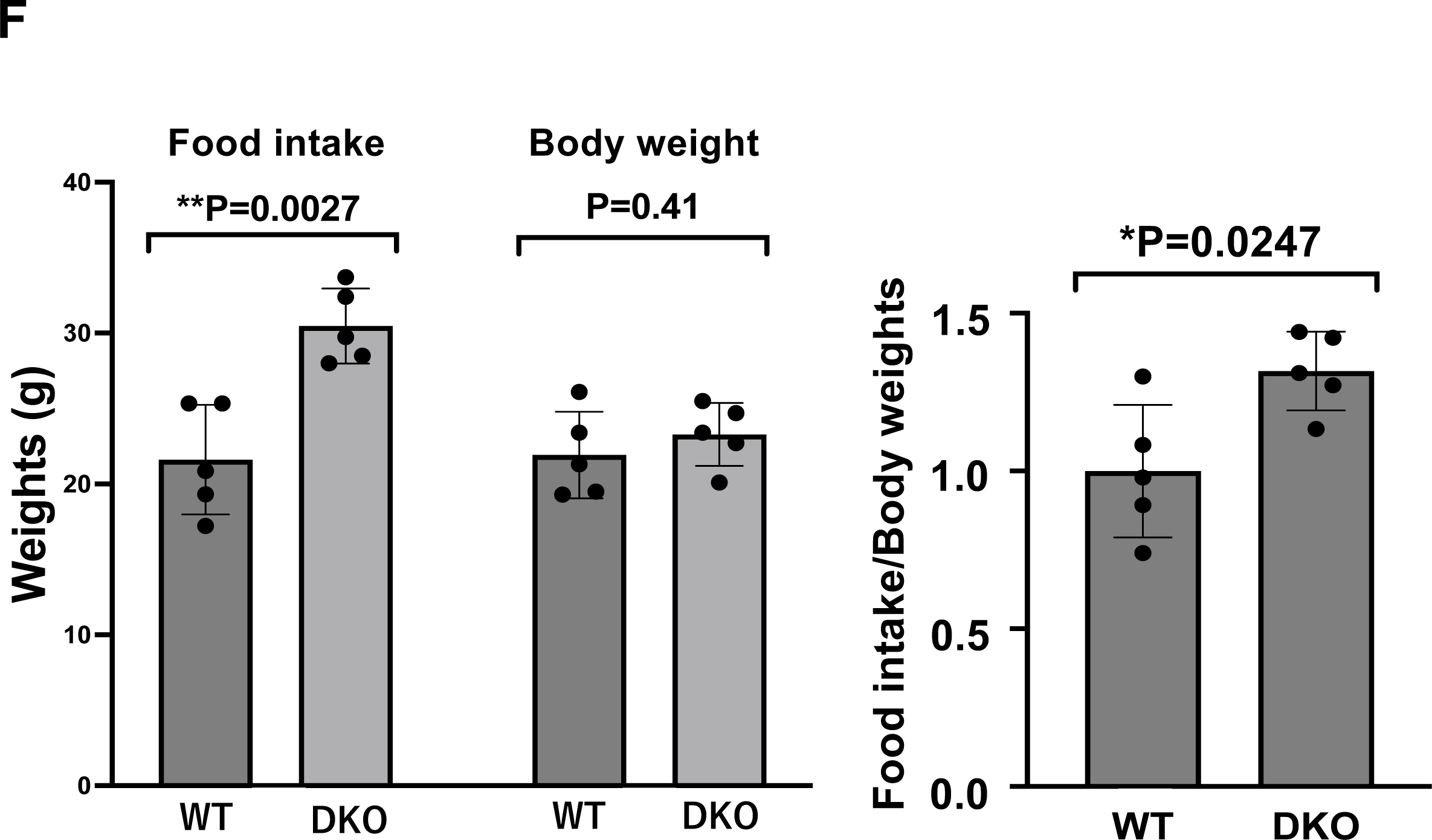
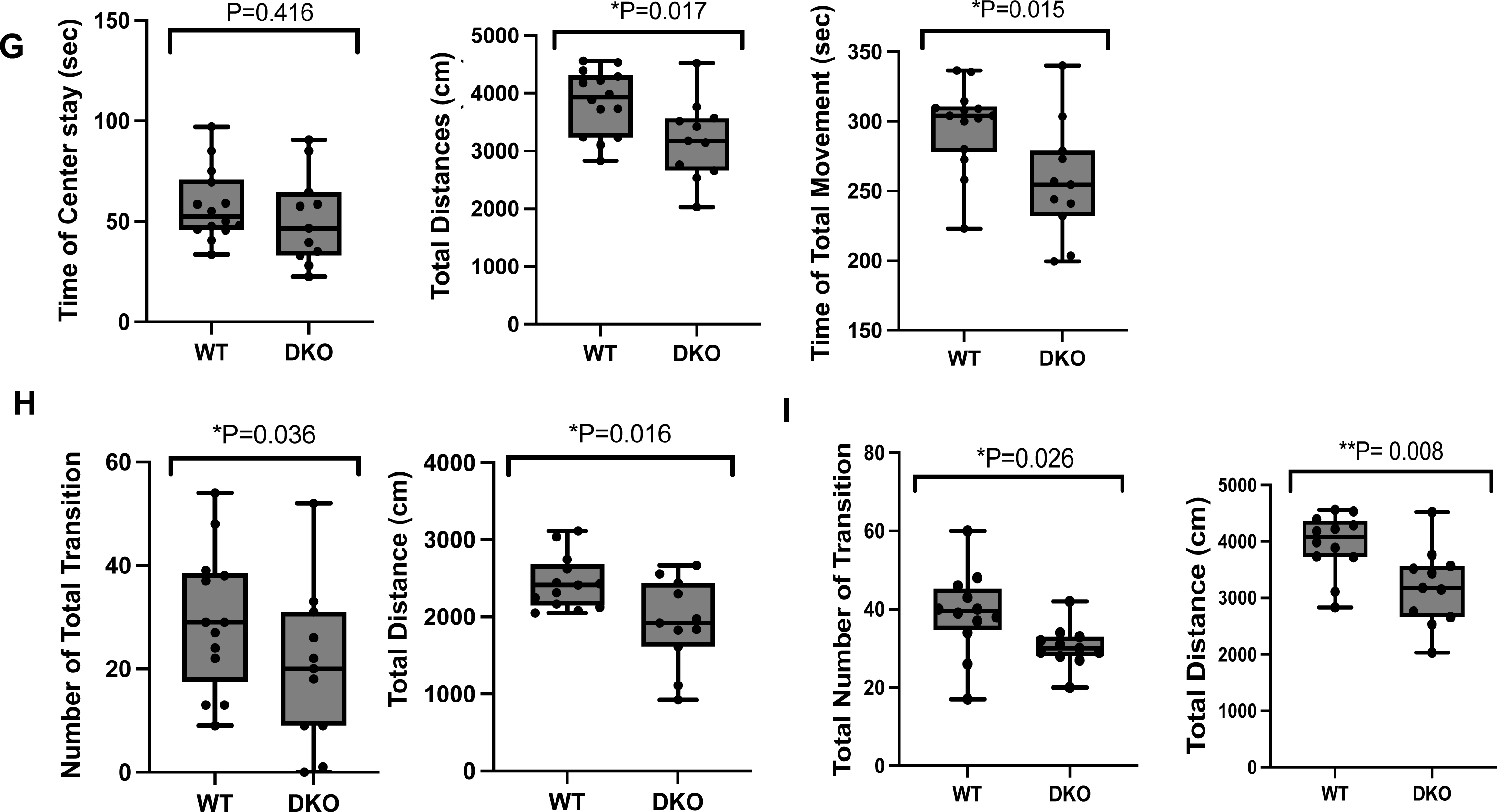
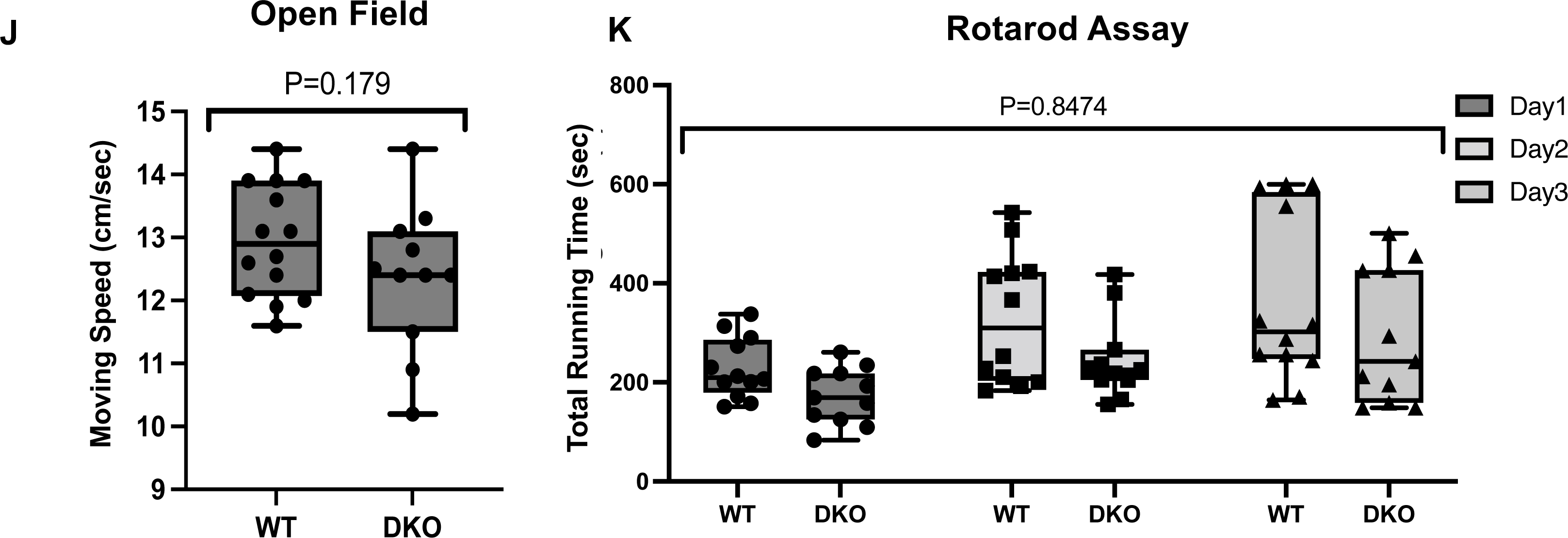
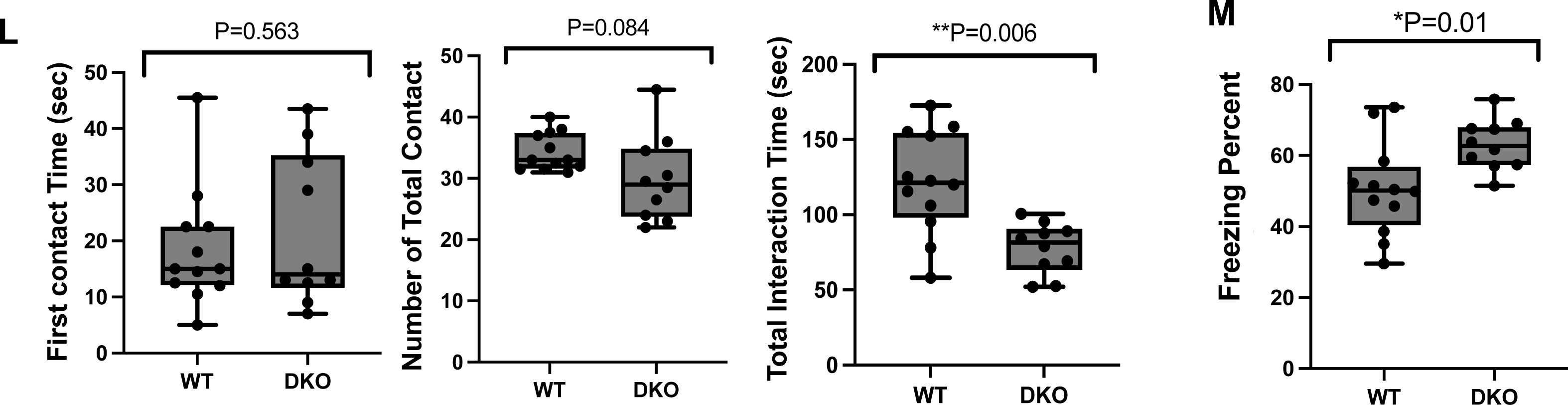
Obesity and abnormal behaviors of *Rtl8a* and *8b* DKO. **(A)** One example of the maximal weight of *Rtl8a* and *8b* hetero DKO females (bottom, 75 g) compared with a WT littermate female (top, 34 g) at 80 w. **(B)** Weight of hemizygous DKO males (n=11) and littermate males (n=14) at 12 w. (**C** and **D**) Body weight (bottom) and food intake every two weeks (top) of the WT (n = 5) are shown. Hetero DKO female (n =5) mice and littermate WT females (n = 5) (**C**) and hemizygous DKO males (n =5) and littermate WT males (n = 5) (**D**). Blue: WT, Red: DKO. * P < 0.05. (**E** and **F**) Overeating of hetero DKO females at 10 w (**E**) and hemizygous DKO males at 9 w (**F**). Left: Food intake for one week (females between 9-10 w and males between 8-9 w) and their body weight (females at 10 w and males at 9 w). Right: Their ratio indicates that both DKO females and males exhibited overeating just before becoming obese. **(G-M)** Behavior tests. Reduced locomotive activity was observed in the OF test **(G) (**WT: n = 12, DKO: n = 10, t-test), L/D transmission test **(H) (**WT: n = 14, DKO: n = 11, t-test) and EPM test **(I)** (WT: n = 12, DKO: n = 11, t-test). *P < 0.05, **P < 0.01. Normal physical capability was observed in the OF test **(J)** (WT: n=14, DKO: n=11, t-test) and rotarod assay **(K)** (WT: n = 12, DKO: n = 11, two-way ANOVA 0.8474). Reduced social activity was observed in the social behavior test **(L)** (WT: n = 12, DKO: n = 10, t-test). Increased apathy-like behavior was observed in the tail suspension test **(M)** (WT: n = 12, DKO: n = 10, t-test). *P < 0.05, **P < 0.01.

Both the hetero DKO females and DKO males gradually became obese after 8 weeks of age (w) in association with the increased food intake and this trend continued thereafter (Figs. 2A-F). For example, one of the maximum weights of the hetero DKO females reached 75 g at 80 w, compared to the 34 g of a WT littermate female (Fig. 2A) and average weights of hemizygous males and WT littermate males are 29.4 g and 28.6g at 12 w (Fig. 2B). Interestingly, they exhibited significant overeating (increased food intake/weight ratio) at 9-10 w (hetero DKO females) and at 8-9 w (DKO males) (indicated by red # marks in Figs. 2C and D), just before the onset of obesity (Figs. 2E and F). These findings suggest some dysfunction of the hypothalamus, a key region in appetite control.

### Abnormal behaviors in *Rtl8a/8b* DKO mice

The DKO mice exhibited reduced locomotive activity, decreased social responses and increased apathy. In the open field (OF: Fig. 2G), light/dark transition (L/D: Fig. 2H) and elevated plus-maze tests (EPM: Fig. 2I), the DKO males exhibited reduced locomotive activity in terms of the total distance (3953 vs 3191 cm in OP; t-test 0.017, 2373 vs 1925 cm; t-test 0.016 in L/D, and 1961 vs 1653 cm; t-test 0.023 in EPM), total movement (296 vs 257 sec; t-test 0.015 in OF) and number of transitions (29.7 vs 17.9 times; t-test 0.036 in L/D and 37.5 vs 30.4 times; t-test 0.046 in EPM). The other elements of these tests were normal (Figs. S9 and S10) including normal moving speed (OF: Fig. 2J) and a normal score on the rotarod test (Fig. 2K). Although the average scores of the DKO were lower than those in the WT, but they are not statistically significant, suggesting their normal physical functioning.

In the social activity test, the DKO males had similar first contact time and the number of total contacts compared with their WT littermates, but the total interaction time was reduced (121.5 vs 81.6 sec; t-test 0.006), suggesting a reduction in social activity (Fig. 2L). In the tail suspension test (Fig. 2M), their freezing time was increased (50.3 vs 63 sec; t-test 0.01), suggesting increased apathy. We did not observe any abnormality in the fear conditioning test except for lower locomotive activity (Fig. S11). We also found that some DKO males were severely injured by attacks from other DKO males, while certain other DKO males suffered severe self-inflicted injuries by biting and scratching themselves during normal breeding. These findings suggest that the loss of RTL8A/8B somehow affects neuronal functions in the brain, possibly in the cerebral cortex (see later sections).

### mRTL8A/8B protein expression in the brain

mRTL8A/8B proteins were detected in certain specific regions, such as the hypothalamus, frontal/temporal cortex, striatum and olfactory bulb (Figs. 3A-P and Fig. S12), which appear to be associated with the abnormal DKO phenotypes mentioned above. There were no significant differences in brain or cortex weight between DKO and WT at 20 w (Fig. S13). For the detection of the mRTL8A/8B proteins, we used an in-house made anti-mRTL8A/8B antibody by immunization with their unique N-terminal peptides (Fig. 1B and Fig. S2). We confirmed the antibody specificity by checking whether the WT signals disappeared in the DKO brain (Figs. 3F-J and L, N, P).

**Figure 3.**
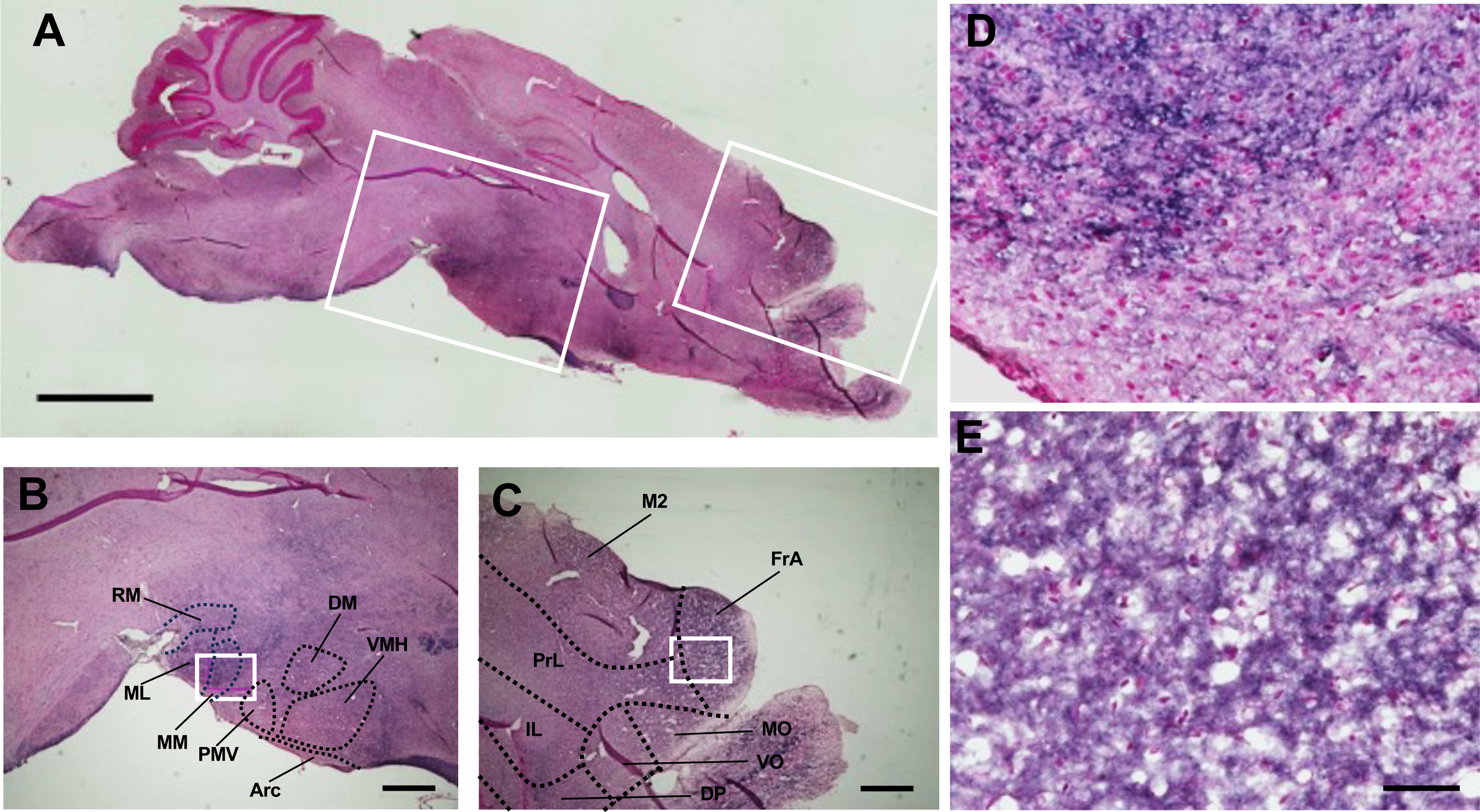
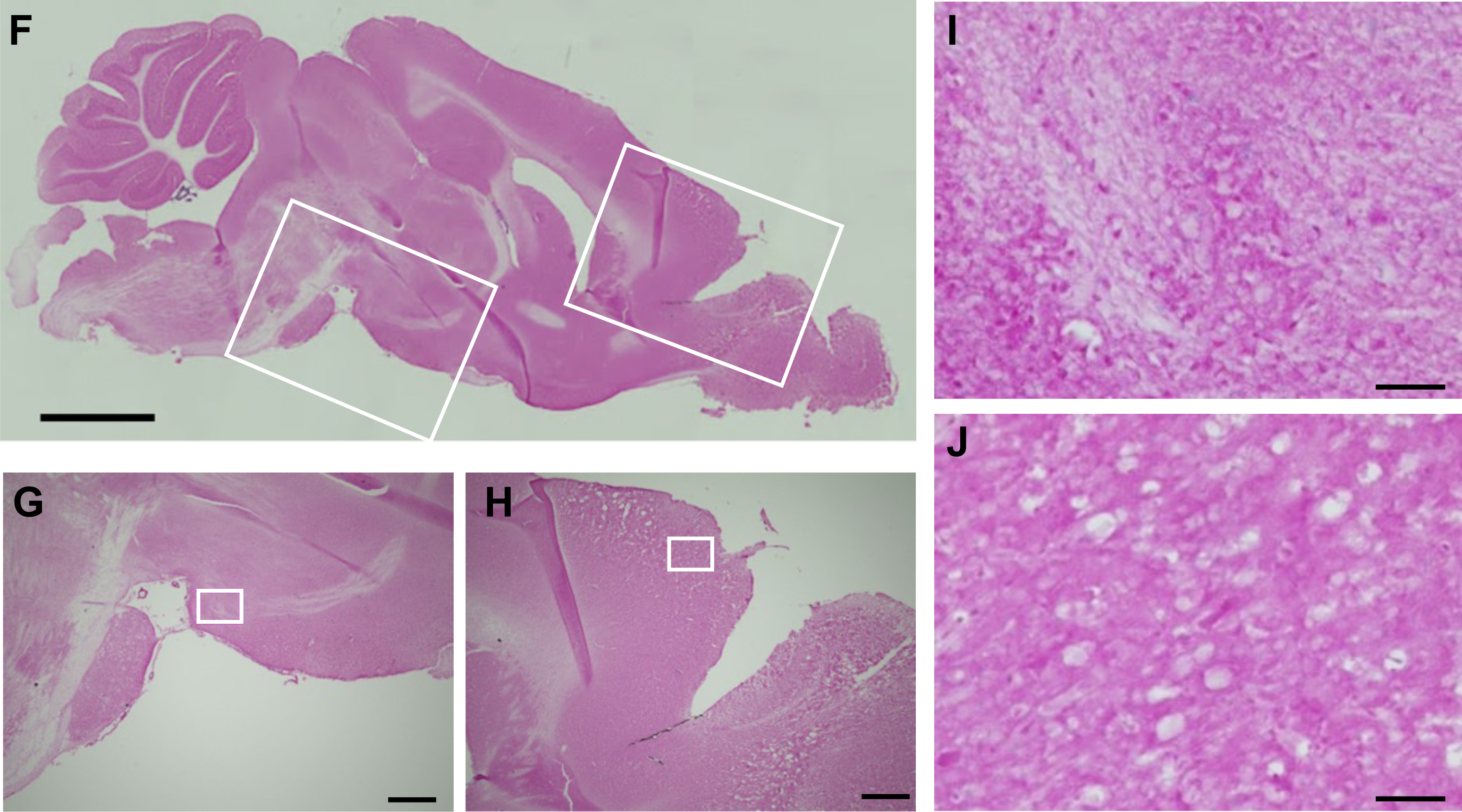
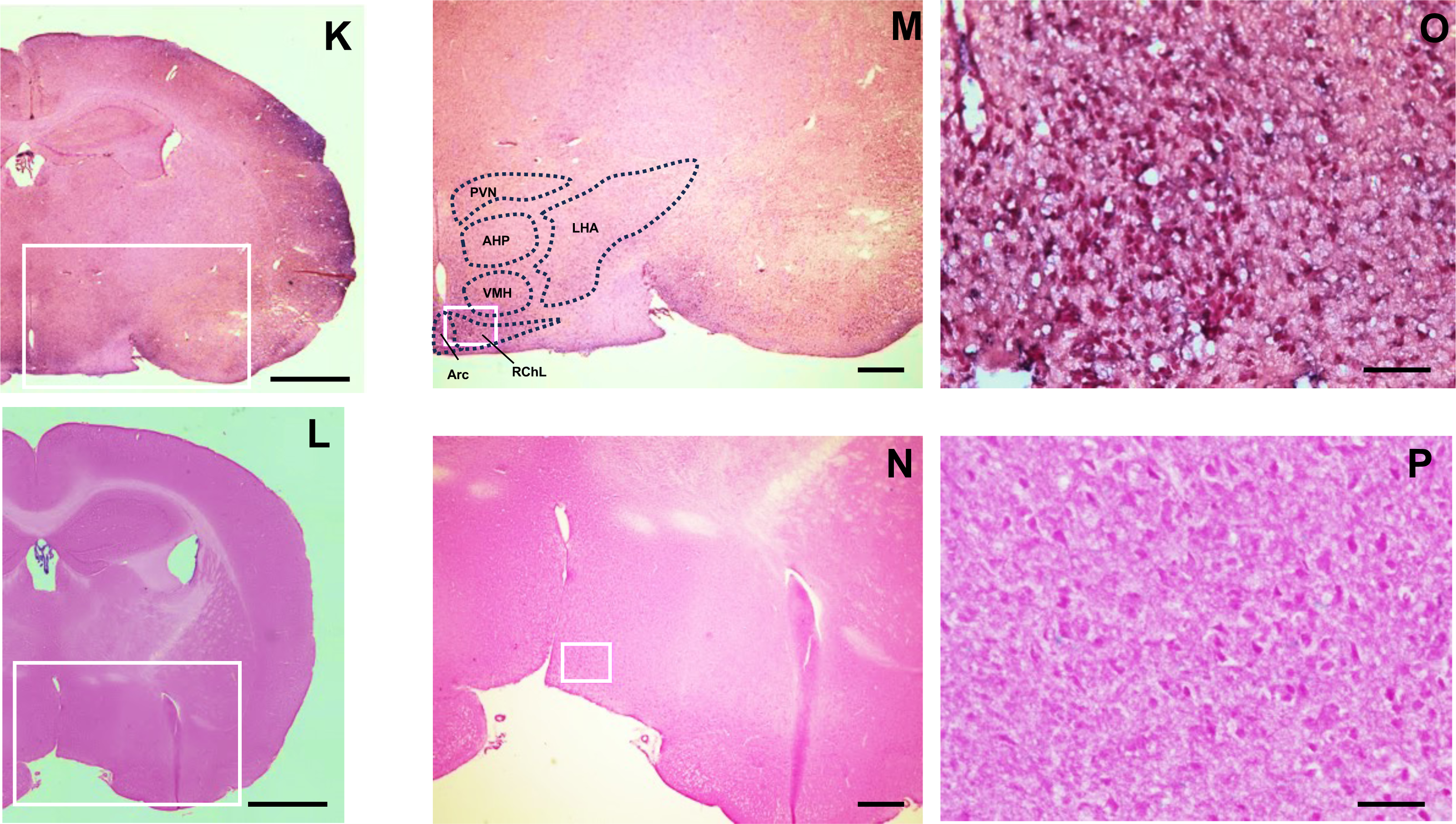
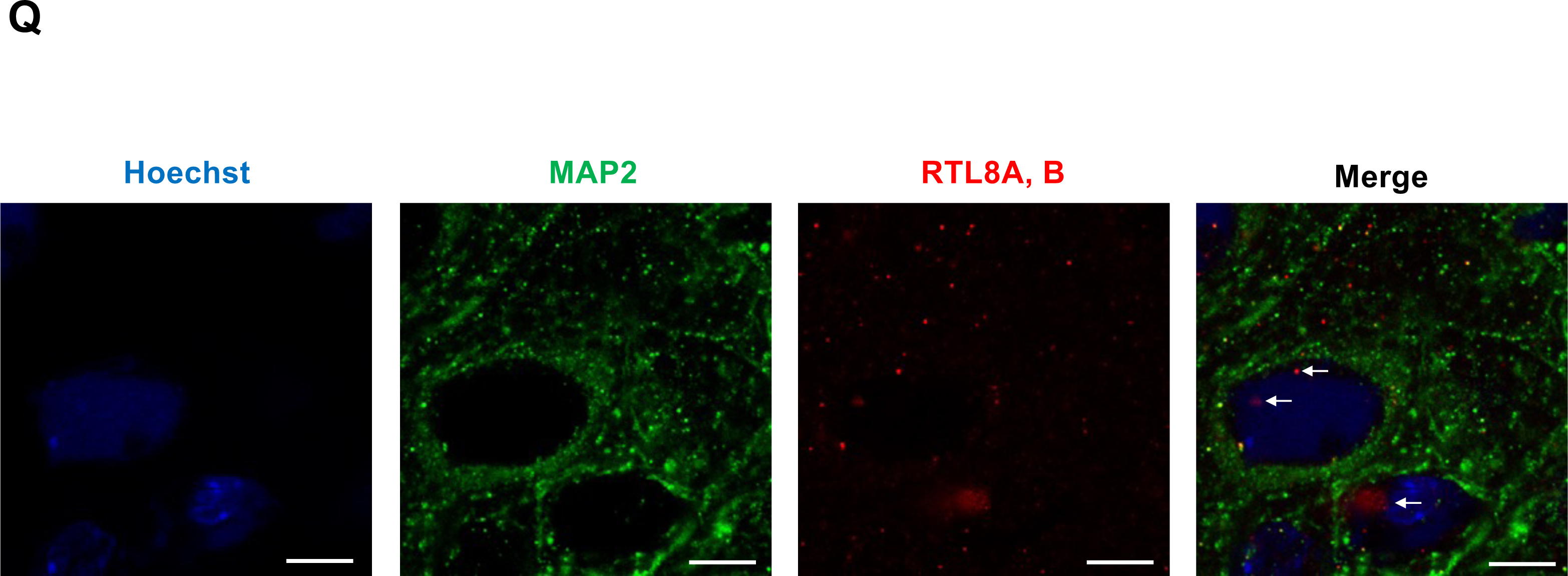
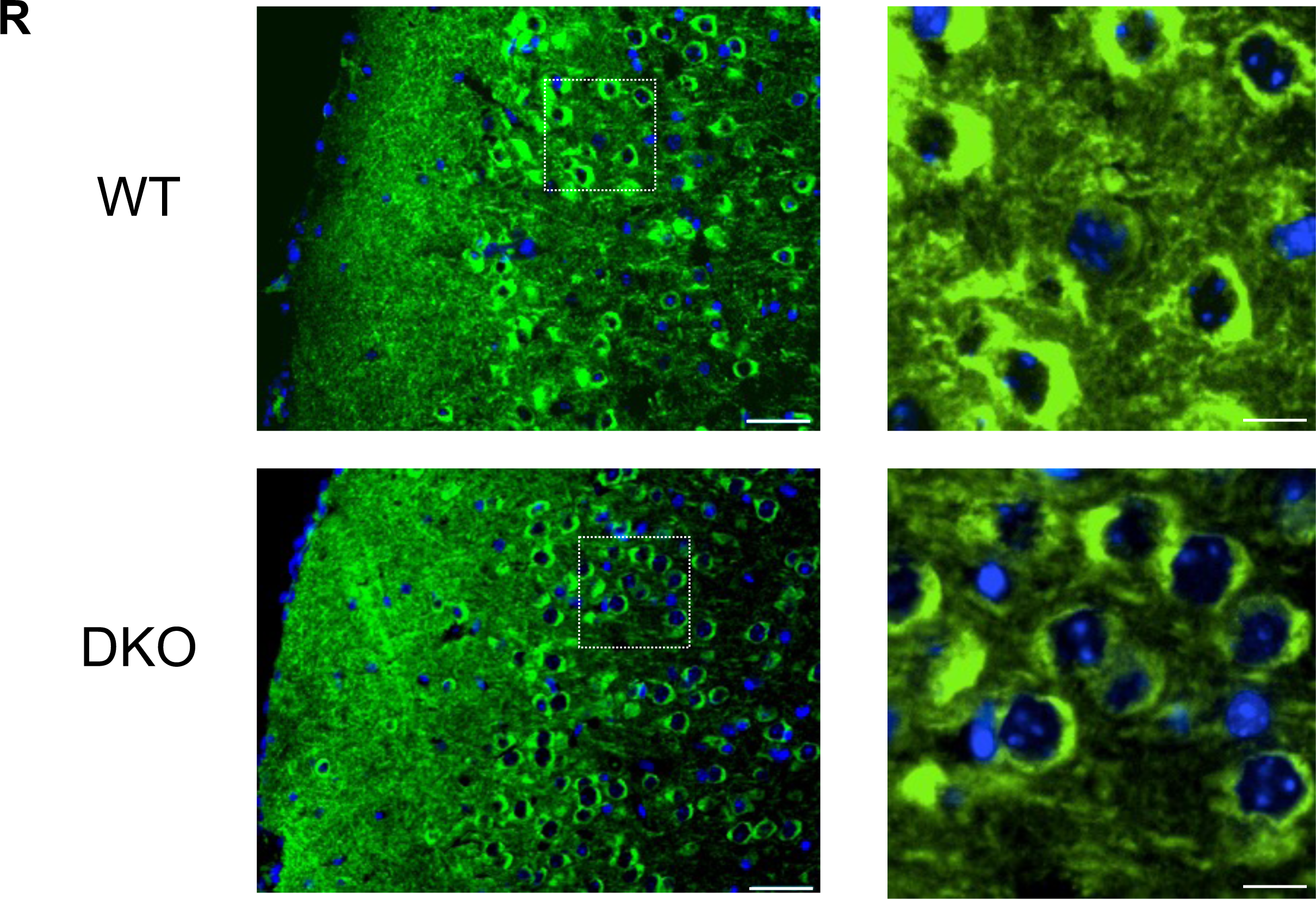
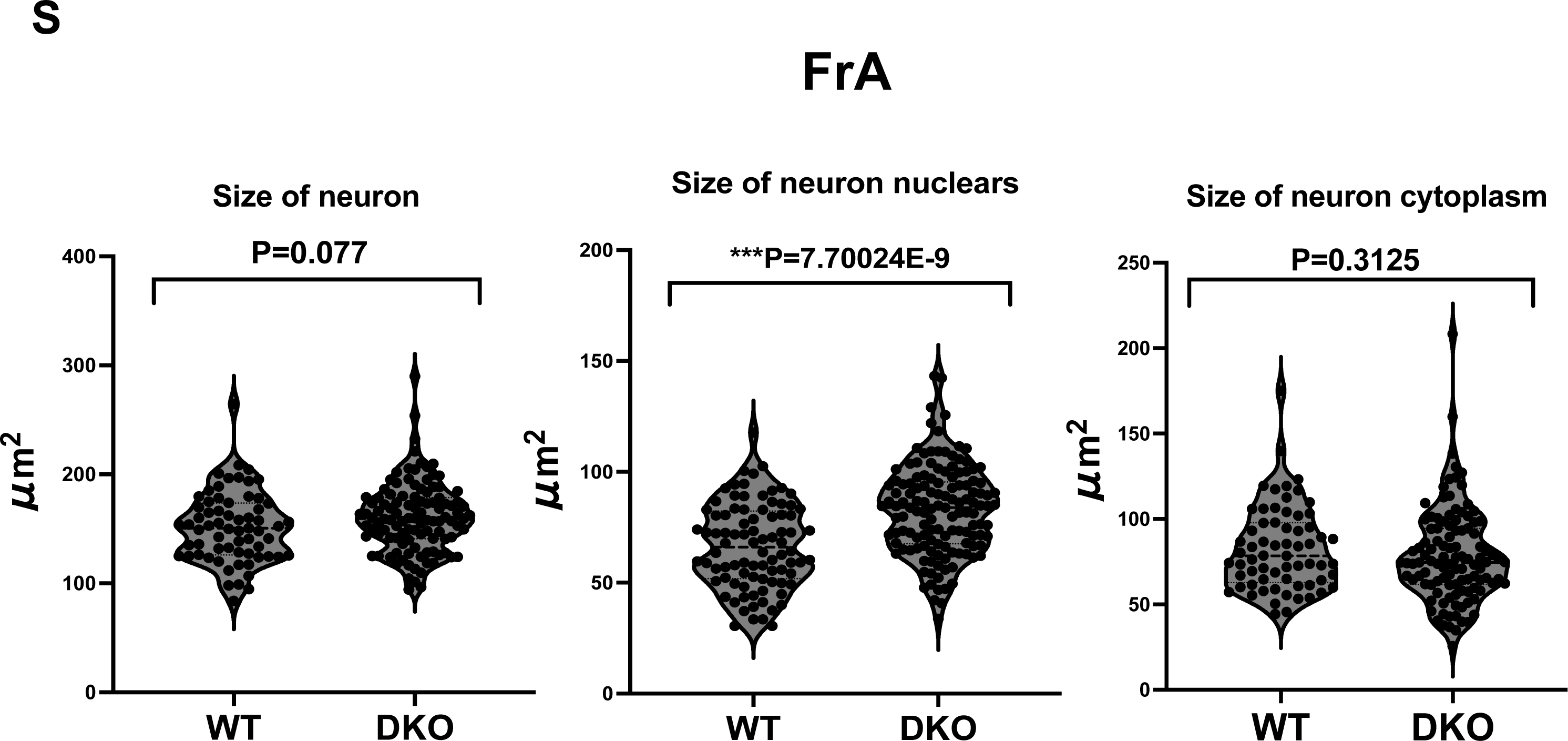
Immunostaining of mRTL8A and 8B proteins in the brain. **(A)** A sagittal section image of the WT mice at 20 w. mRTL8A/8B protein was detected in the frontal cortex and hypothalamus. Bar: 1mm. **(B)** An enlarged hypothalamus area surrounded by a central white square in **(A)**. Arc: Arcuate nucleus, DMH: Dorsomedial hypothalamic nuclei, ML: medial mammillary nucleus, MM: Medial mammillary nucleus, PMV: premammillary nucleus, ventral part, RM: retromammillary nucleus, VMH: Ventromedial hypothalamic nuclei. Bar: 200 μm. **(C)** An enlarged frontal cortex area surrounded by a right white square in **(A)**. DP: dorsal peduncular cortex, FrA: frontal association, IL: infralimbic cortex, M2: secondary motor cortex, MO: medial orbital cortex, PrL: Prelimbic cortex, VO: ventral orbital cortex. Bar: 200 μm. (**D**) A magnified view of a white square in (**B**). Bar: 50 μm. (**E**) A magnified view of a white square in (**C**). Bar: 50 μm. (**F-J**) Corresponding sagittal section images of the DKO mice at 20 w to (**A-E**). **(K** and **L)** Coronal section images of the WT and DKO at 20 w. Bar: 1mm. **(M** and **N)** An enlarged area surrounded by a white square in **(K)** and **(J)**, respectively. AHP: anterior hypothalamus nucleus, LHA: lateral hypothalamic area. PVN: paraventricular nucleus, RChL: Retrochiasmatic area, lateral part. Bar: 200 μm. **(O** and **P)** A magnified view of a white square in (**L**) and (**M**), respectively. Bar: 50 μm. **(Q)** Co-immunofluorescence staining images of MAP2 and RTL8A/8B in the FrA region in WT (sagittal sections, each at 20 w, n=2). Arrows indicate RTL8A/8B in the nuclei. Scale bars: 5 μm. Blue: Hoechst, Green: MAP2, Red: RTL8A, B.**(R)** Immunofluorescence staining of FrA in WT and DKO mice. Left: Immunofluorescence staining images of MAP2 in the FrA region in WT and DKO mice (sagittal sections, each at 20 w, n=2). Right: Enlarged images of the dashed box areas in the left pictures. Green: MAP2, blue: DAPI. Scale bars: 40 μm (left) and 10 μm (right), respectively. **(S)** The sizes of the neuronal cells and their nuclei in (**R**) were measured by a blind observer using ImageJ. Three individuals from each of the WT and KO mice were examined for the measurement of a total of 65 and 109 neurons, 88 and 132 neuronal nuclei, and 65 and 109 neuronal cytoplasm samples, respectively (see also Fig. S15). All measurements were statistically analyzed with Student’s t-test and the results are displayed as Violin Plots. *P < 0.05, **P < 0.01, ***P< 0.001.

mRTL8A/8B were detected in a wide area of the prefrontal cortex (PFC), including the orbital cortex, prelimbic cortex (PrL) and frontal association (FrA) (Figs. 3C and D), which may be related to abnormal behaviors in DKO mice. The orbitofrontal cortex (OFC) is a ventral subregion of the PFC (Szczepanski & Knight, 2014), and involved in the integration of sensory information, emotion processing, decision-making and behavioral flexibility (Hare et al, 2009; Hare & Duman, 2020). The PrL is involved in depression-like despair behavior (Pizzagalli & Roberts, 2022) and maintaining attention to category-relevant information, and flexibly updates category representations (Broschard et al, 2021), while the FrA is engaged in stimulus integration during associative learning (Nakayama et al, 2015). The FrA receives projection from the insular (IC) and perirhinal (PRh) cortices in the temporal lobe where high mRTL8A/8B expression was also detected (Fig. S12). Functional impairment of all these regions may explain the reduced locomotive activity and social interaction as well as the increased apathy-like behavior of the DKO mice.

mRTL8A/8B were also detected in the most hypothalamic regions (Figs. 3A, B, D, K, M and O), including several important hypothalamic nuclei related to feeding behavior, such as the arcuate nucleus (Arc) (Lanfray & Richard, 2017), dorsomedial and ventromedial hypothalamic nuclei (DMH and VMH) (Merkestein et al, 2014), paraventricular nucleus (PVN) (Yousefvand & Hamidi, 2020), anterior hypothalamic area, posterior part (AHP) and lateral hypothalamus(LHA) (Figs. 3E and F). In addition, they were detected in the nucleus accumbens (Acb) in the ventral striatum (Fig. S12) and medial forebrain bundle integrated in the LHA that connects the Acb to the ventral tegmental area (VTA) in the midbrain, two important control centers for feeding behavior (Atasoy et al, 2012).

### Neuronal expression of RTL8A/8B

Co-immunostaining experiment of mRTL8A/8B with a neuronal marker, microtubule-associated protein (MAP) 2 in the prefrontal cortex, demonstrated that these proteins are localized in both the nucleus (indicated by arrows) and cytoplasm of neurons (Fig. 3Q). It is consistent with previous reports that human RTL8A-C protein is expressed in the iNeurons differentiated from AS and ALS patients’ iPSCs (Pandya et al, 2021; Whiteley et al, 2021) and also with the fact that human RTL8A/8B are involved in the nuclear transition of UBQLN2 that is necessary for nuclear protein quality control (Mohan et al, 2022). It is also consistent with our results that mRTL8A/8B with the N-terminal NLS-like signal is accumulated in the nuclei of NIH3T3 cells while mRTL8C without it is located in the cytoplasm (Fig. 1C).

Immunofluorescence staining with anti-MAP2 antibody also demonstrated that the nuclei of the layer 2/3 pyramidal cells in the FrA were significantly increased in size (Figs. 3R and S, Figs. S14 and S15) in the 20 w adult brain. This was not observed in the other cortical regions without mRTL8A/8B expression, such as the primary somatosensory cortex, shoulder region (S1Sh) (Figs. S16-S19). It has been reported that the pyramidal cells in the mPFC moderate stress-induced depressive behaviors (Shrestha et al, 2015), suggesting that this neuronal alteration is related to the apathy-like behavior observed in the DKO mice possibly due to the abnormal nuclear protein quality described above. We did not observe any signs of neuroinflammation by immunofluorescence staining for Ubiquitin and glial fibrillary acidic protein (GFAP) (Fig. S20).

### Reduced expression of GABRB2 in the DKO cerebral cortex

RNAseq analysis suggested that *Gabrb2*, the β2 subunit of the GABA_A_ receptor known to be an important psychiatric factor, was downregulated in the DKO cerebral cortex, and this was confirmed at the protein level by Western blotting. To elucidate the cause of the abnormal behavior in DKO, we performed RNAseq analysis on the 20 w cerebral cortex of DKO and WT. Importantly, the gene set enrichment analysis (GSEA) using the GO cellular component terms (false discovery rate (FDR) < 0.05) revealed that the top two most downregulated gene sets were “integral component of presynaptic membrane” (6e-09) and “GABAergic synapse” (1e-08) (Figs. 4A-D), and only three GO terms, “respirasome”, “respiratory chain complex” and “mitochondrial respirasome” were significantly upregulated (Figs. 4A and B, Figs. S21-23). However, among them, only *Gabrb2* and *Adra1a* remained as downregulated genes under the condition of 2.0-fold increase and decrease in a total of 20 up-regulated and 28 down-regulated genes (Figs. 4E and F).

**Figure 4.**
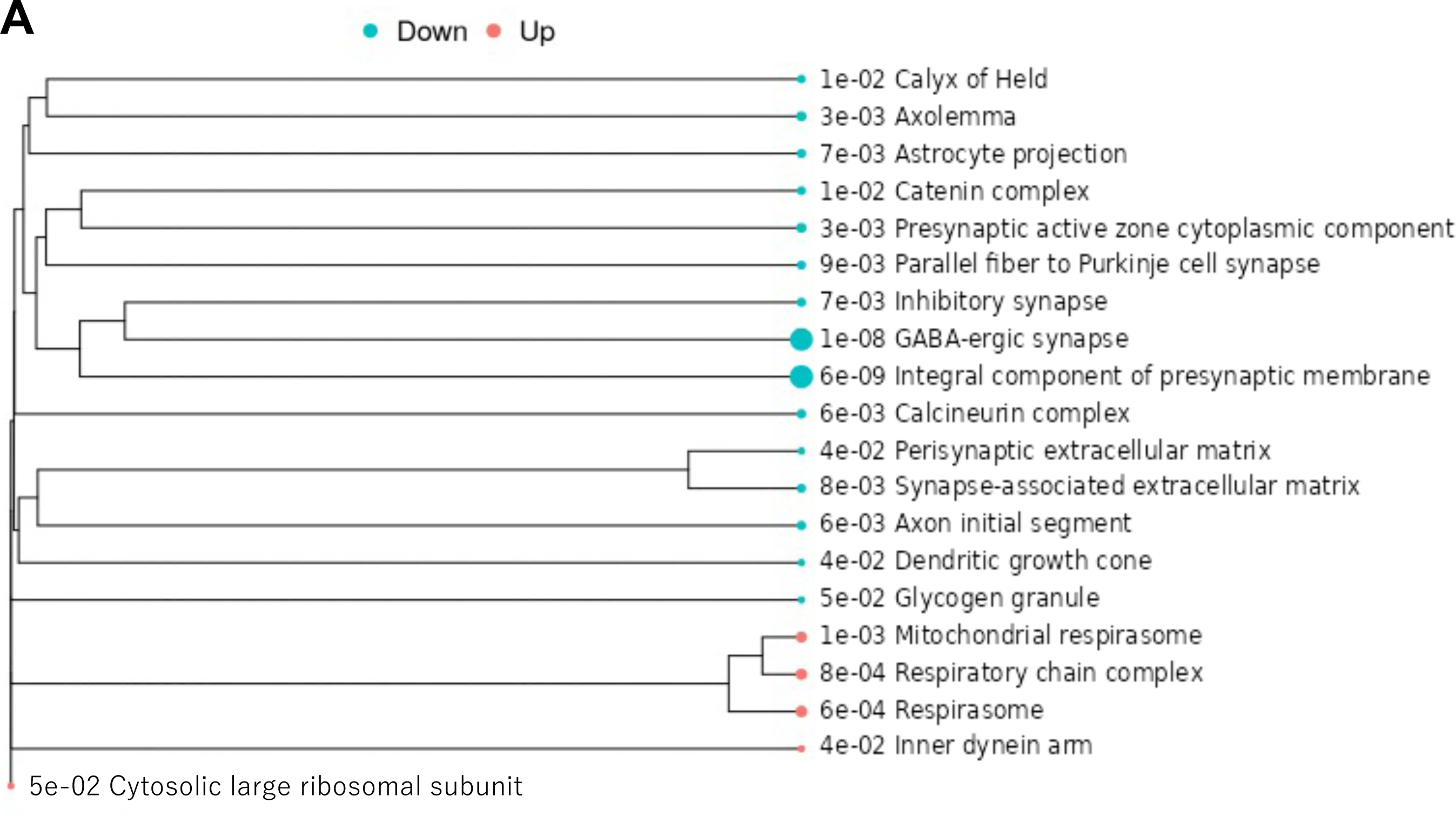
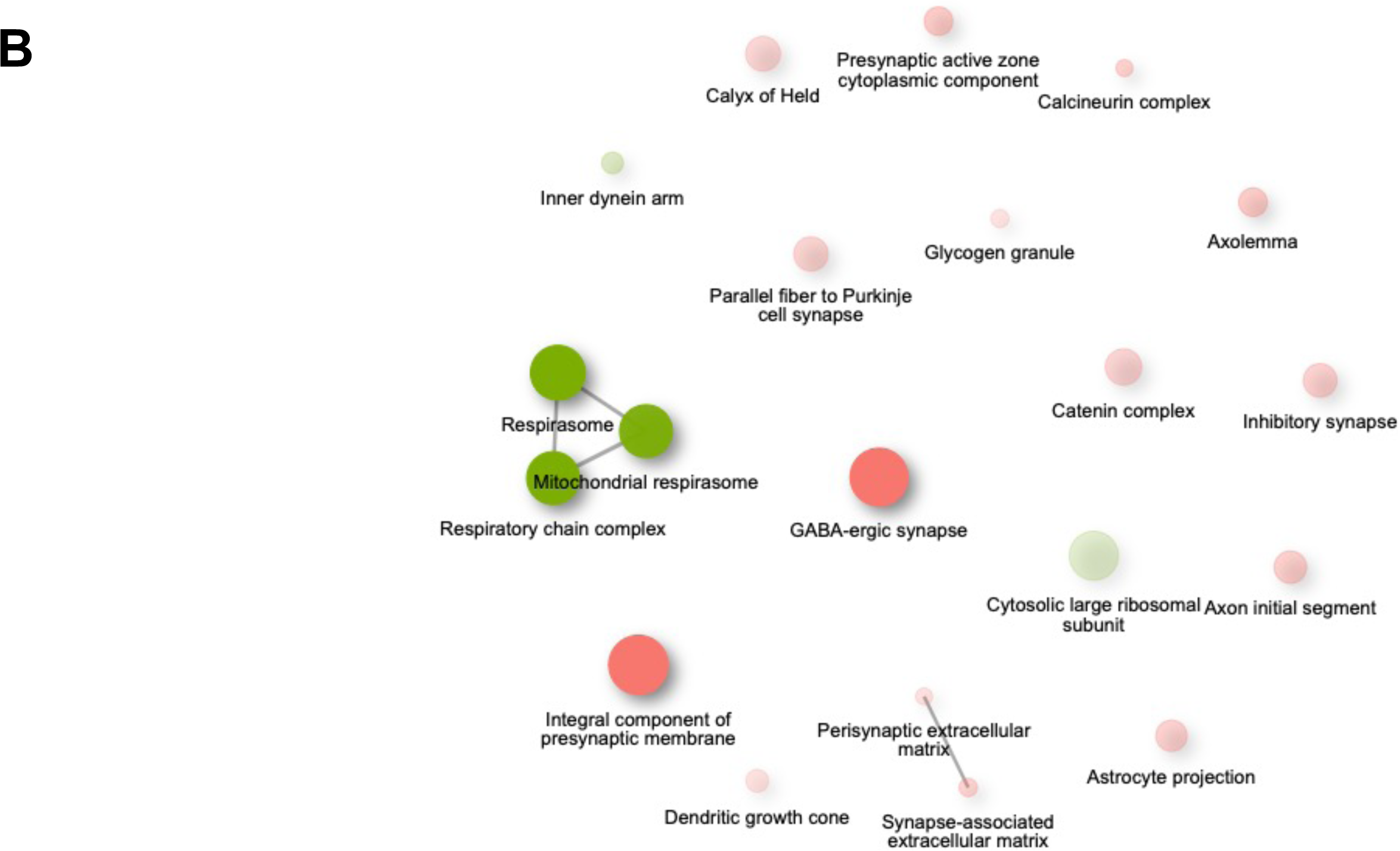
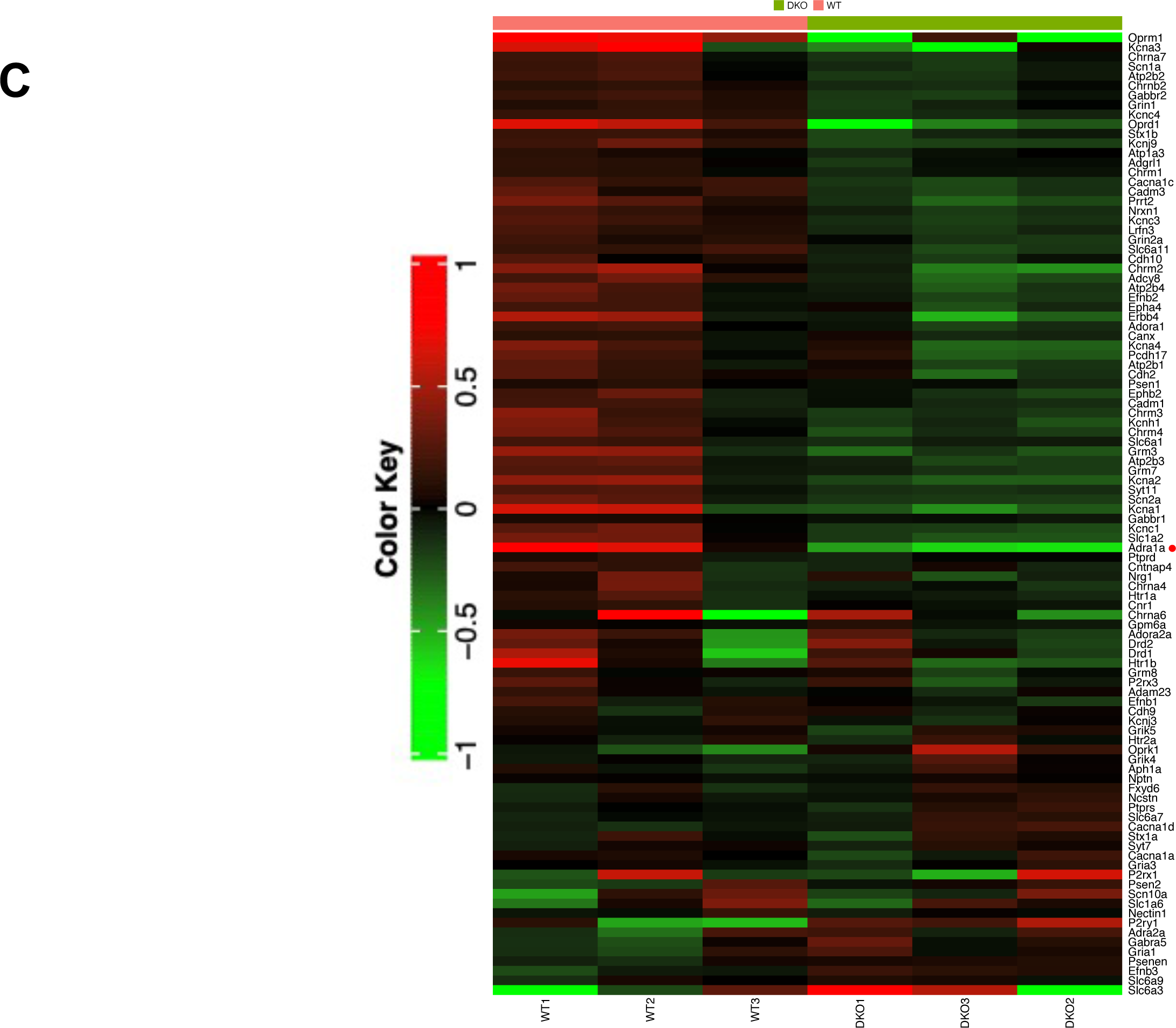
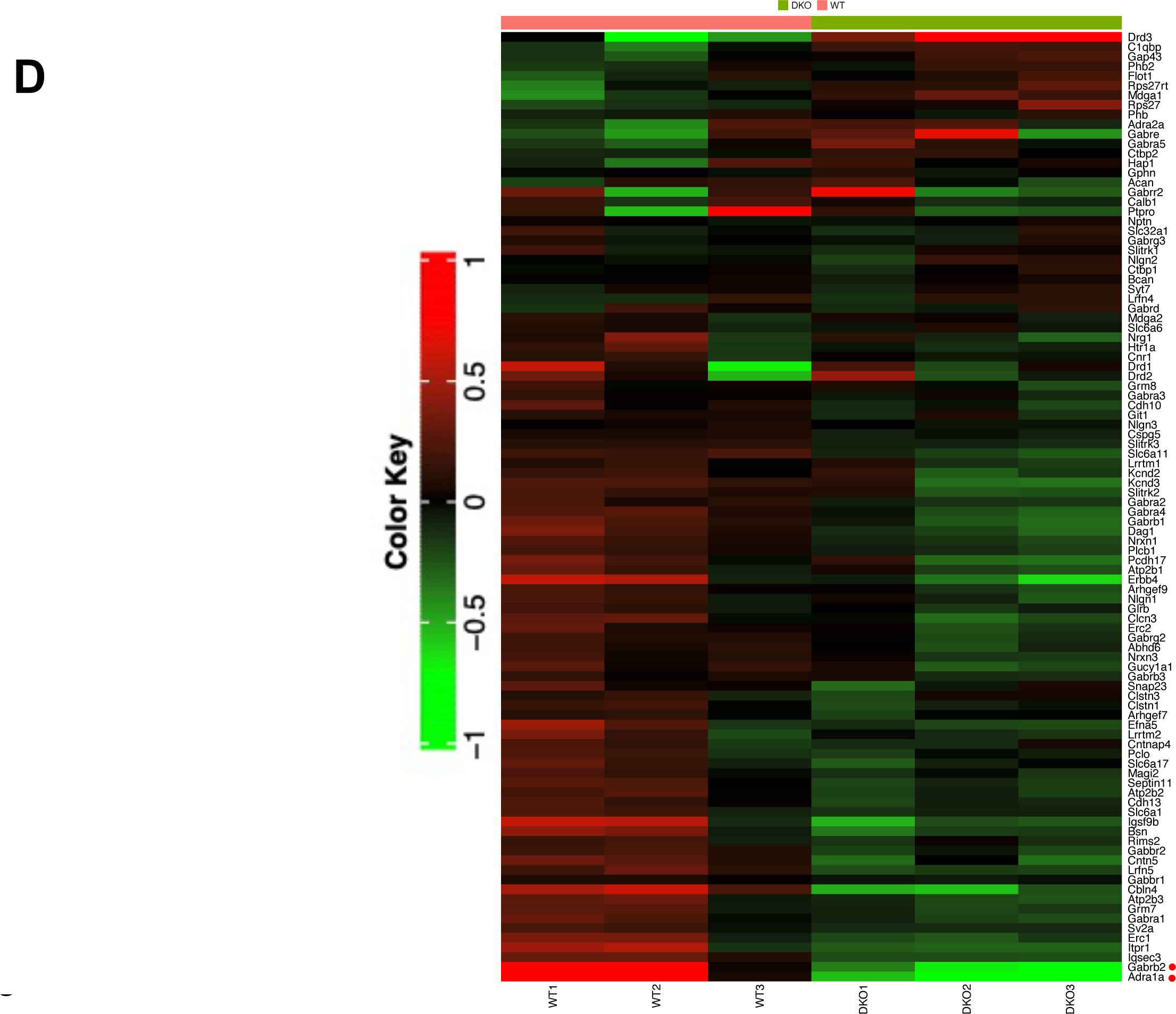
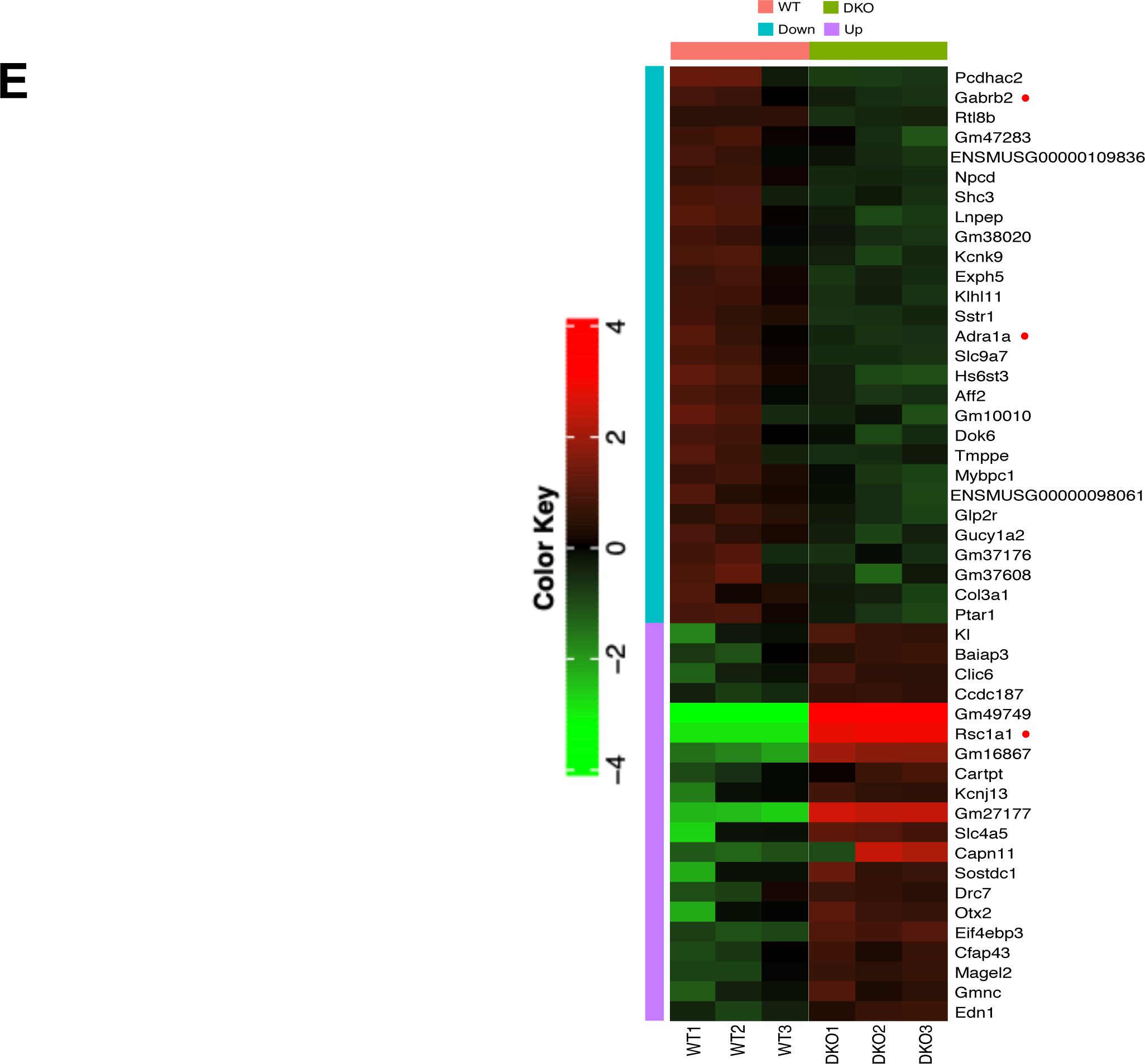
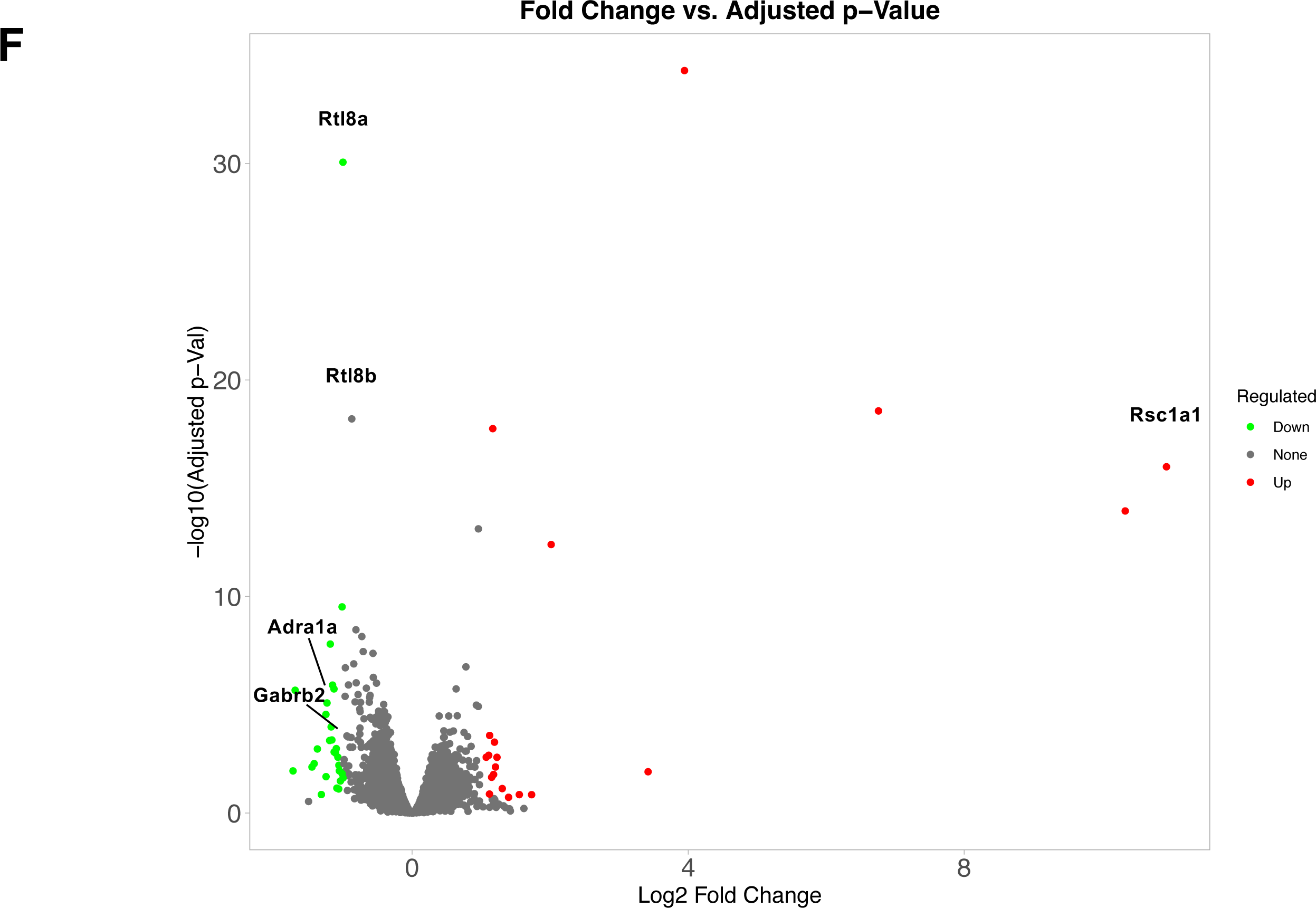
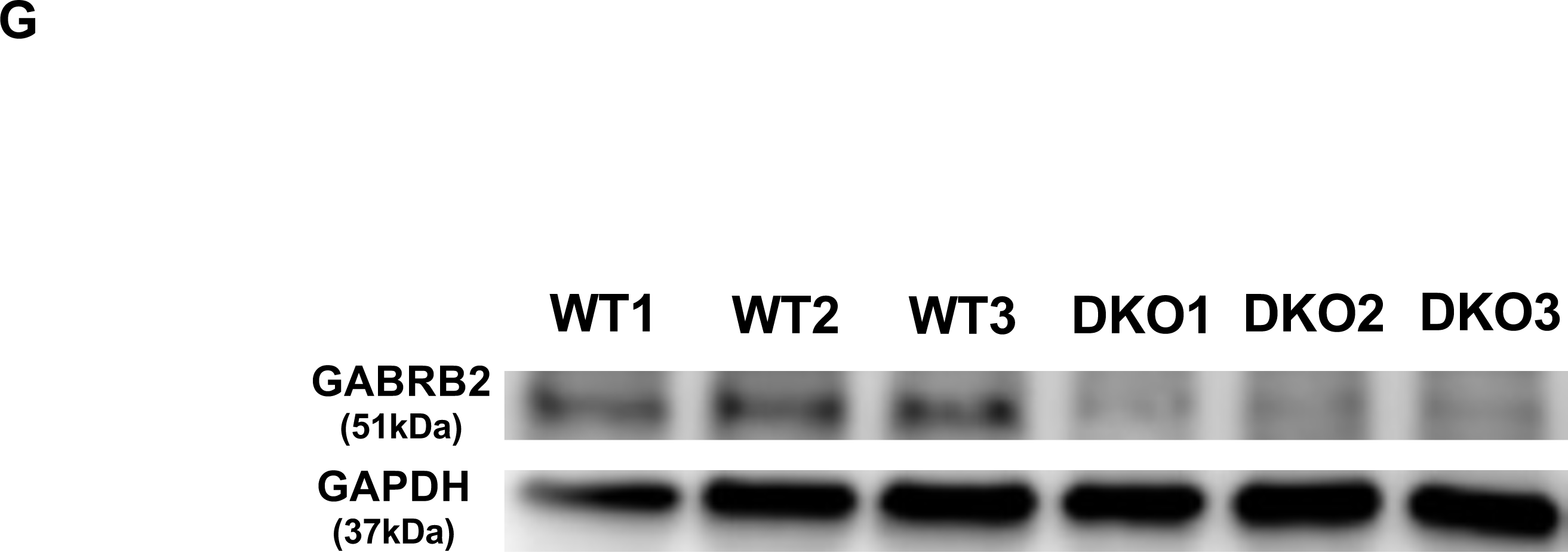
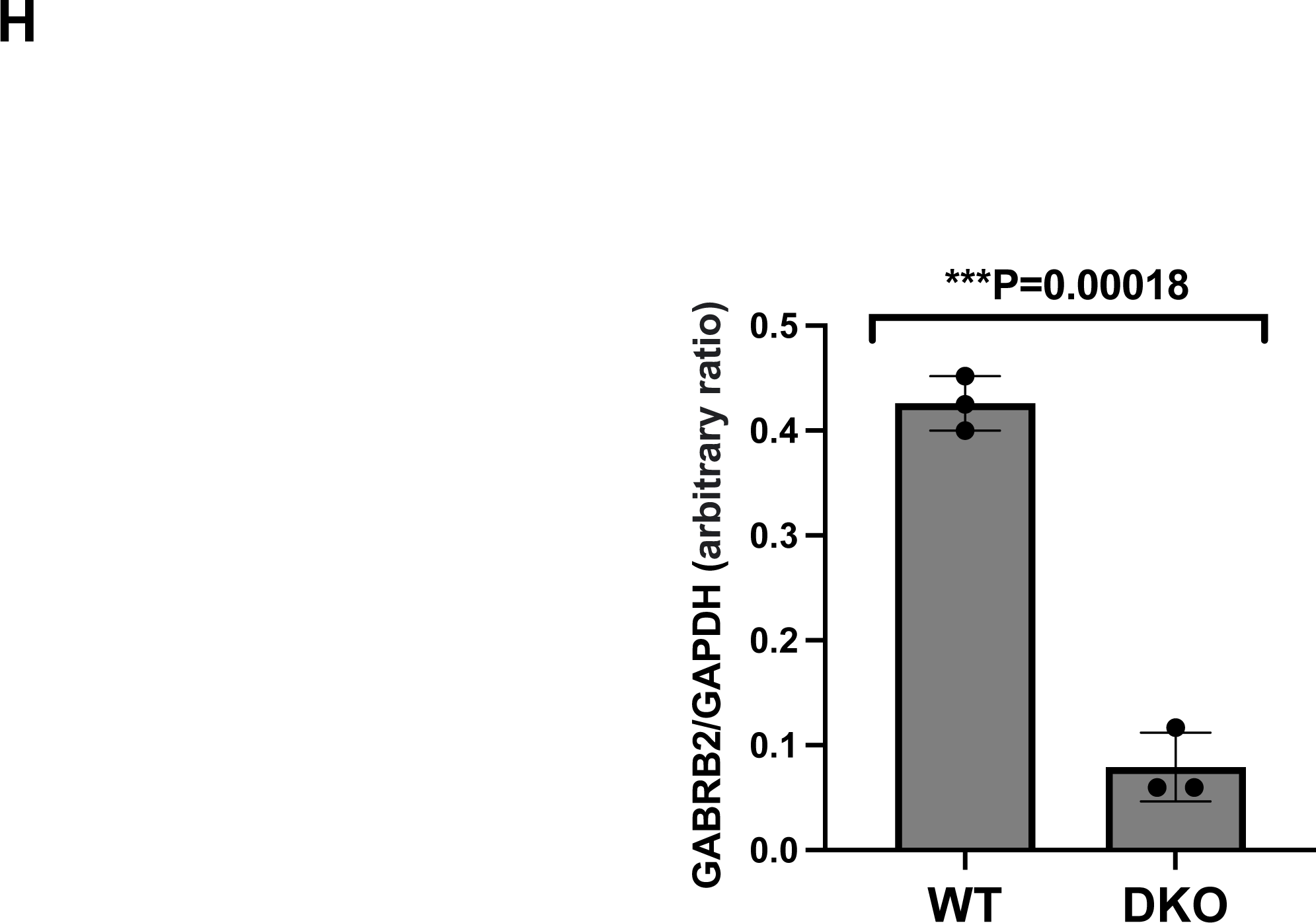
Differentially expressed genes between DKO and WT cerebral cortex. (**A** and **B**) The tree plot shows the result of gene set enrichment analysis using the GO cellular component terms (FDR < 0.05). Light blue and pink represent downregulated and upregulated genes, respectively. Circle size correlates with the number of genes in each gene set. (**B**) The network diagram shows the relationship between the results. Two pathways (nodes) are connected if they share 30% (default, adjustable) or more genes. Green and red represent up- and down-regulated pathways, respectively. Darker nodes represent more significantly enriched gene sets. Larger nodes represent larger gene sets. Thicker edges represent more overlapping genes. (**C** and **D**) The heat maps show the expression level (normalized Fold Change (FC) of each individual for the gene expression level categorized by “integral component of presynaptic membrane” (**C**) and “GABAergic synapse” (**D**). Red dots indicate genes suspected to be associated with the DKO phenotype. Left panel color indicates Log2FC. (**E)** The heatmap of a total of 20 up-regulated and 28 down-regulated genes that exhibited 2.0-fold increase and decrease, respectively. Red dots indicate genes suspected to be associated with the DKO phenotype. **(F)** The volcano plot shows the FC and the false discovery rate (FDR) of the genes analyzed in (**A**). The names of the genes that appear to be particularly associated with the DKO phenotype are indicated. X= FC, Y=-log10(FDR). (**G**) The results of Western blotting using antibodies against GABRB2 and GAPDH. (**H**) The results of (**G**) quantifying the expression levels using imageJ’s Gel tool.

Both *GABRB2* and *ADRA1A* have been reported to be associated with several psychiatric disorders: The former with SZ, BP, FTD and ASD (Lo et al, 2004; Petryshen et al, 2005; Ma et al, 2005; Lo et al, 2007; Chen et al, 2009; Yuan et al, 2015; Jiang et al, 2018; Barki & Xue, 2022) and the latter with attention deficit hyperactivity disorder (ADHD), SZ and generalized anxiety disorder (GAD) (Clark et al, 2005; Elia et al, 2009; Zhang et al, 2018), thus making them good candidates for the DKO phenotypes. We hypothesized that the downregulation of *Gabrb2* in the cerebral cortex is a better candidate to explain the abnormalities in DKO mice, as *GABRB2* has been reported to be decreased in the prefrontal cortex of elderly depressed patients (Zhao et al, 2012). We, then, performed Western blot analysis of GABRB2 and confirmed a 5-fold reduction in GABRB2 protein expression in the cerebral cortex of 1-year-old DKO mice (Figs. 4G and H).

## Discussion

### Roles of eutherian-specific RTL8A, 8B and 8C in the brain

This study demonstrates that the retrovirus-derived *Rtl8a* and *8b* genes, which encode proteins with only 113 aa, play important roles in the brain through their involvement in the regulation of certain basic behaviors, such as, social activity, emotions, maternal care, and food intake. This explains why *RTL8A-C* is well conserved in eutherians (Table S1). Given that there is a gene dosage effect for *RTL8A-C*, this may also explain why *RTL8A-C* exists as a cluster of multiple genes, for example, a kind of triple safety valve in the eutherian brain. In this work, mouse *Rtl8a* and *Rtl8b* were knocked out, however, *Rtl8c* still remained active, so it is of interest to determine whether complete null mice, i.e. *Rtl8a*, *Rtl8b* and *Rtl8c* triple KO (TKO) mice, will exhibit more severe phenotypes and more severe gene expression changes than the DKO mice, as discussed below. A comparison between *Rtl8c* KO, *Rtl8a* and *Rtl8b* DKO and *Rtl8a*, *Rtl8b* and *Rtl8c* TKO mice would also provide important insights into the role of *Rtl8a*, *Rtl8b* and *Rtl8c* in both the cerebral cortex and hypothalamus, the molecular mechanisms of these genes in these abnormalities and the reason why these three genes are evolutionarily conserved in eutherians.

### Down- and up-regulated genes in *Rtl8a and Rtl8b* DKO cerebral cortex

As shown in Fig. 4, *Gabrb2* and *Adra1a* are nominated as good candidate genes responsible for DKO psychiatric abnormalities in the 20 w cerebral cortex as down-regulated genes. Genome-wide association studies (GWAS) reveal that common variants within *GABRB2* are associated with an increased risk of SZ (Lo et al, 2004; Ma et al, 2005; Lo et al, 2007; Chen et al. 2009; Yuan et al, 2015), BD (Yuan et al, 2015), FTD (Jiang et al, 2018) and ASD (Ma et al,2005). In addition, it has been reported that *GABRB2* is decreased in the prefrontal cortex of elderly depressed patients (Zhao et al, 2012). Therefore, it is likely that the downregulation of *GABRB2* in the cerebral cortex (Figs. 4G, H) at least partially causes the reduced social activity and increased apathy in the DKO mice, although the mechanism by which the absence of *Rtl8a* and *8b* downregulates the *Gabrb2* expression requires further investigation. The DKO phenotype exhibits some differences from *Gabrb2* KO mice that exhibit a schizophrenia-like phenotype (Yeung et al, 2018), particularly in terms of increased body weight and food intake. It is likely that the differences are due to the fact that *Gabrb2* KO mice lack GABRB2 expression throughout the brain and that the phenotypes of our DKO mice are more complex because many other candidate genes besides *Gabrb2* are also affected with respect to cerebral cortex function.

As *ADRA1A* has also been reported to be associated with ADHD, SZ and generalized anxiety disorder by GWAS (Clark et al, 2005; Elia et al, 2009; Zhang et al, 2017), a detailed study of *Adra1a* with respect to its contribution to the DKO phenotypes is warranted in the future. It should be noted that Regulator of Solute Carriers 1 (*Rsc1a1*) exhibited an exceptional overexpression (approximately 12-fold) in DKO (Figs. 4E and F). It is possible that the *Rsc1a1* overexpression somehow compensates for the absence of RTL8A/8B. RSC1A1 is located on the intracellular side of the plasma membrane, at the trans-Golgi network, and in the nucleus (Kroiss et al. 2006). Therefore, it is interesting to know whether RTL8A/8B and RSC1A1 proteins colocalize and functionally cooperate in the brain. RSC1A1 possesses the consensus sequences for the ubiquitin-associated (UBA) domain in addition to those for phosphorylation by PKC and casein kinase 2 (Veyhl et al. 2006). UBQLN2 also possesses the UBA domain, which is required for incorporation of UBQLN2 into RTL8 subnuclear puncta, although it is not required for direct binding to RTL8 (Mohan et al. 2022). Therefore, it could also be possible that *Rsc1a1* overexpression somehow disrupts UBQLN2 function, such as trafficking of ubiquitinated proteins to proteasomes, via competition between their UBA domains. As shown in Figs. 4A and B, genes categorized as “respirasome”, “respiratory chain complex” and “mitochondrial respirasome” were also slightly up-regulated (Fig. S21-23). These upregulated genes will be further investigated in the TKO mice in addition to *Rsc1a1*.

### Possible link of *RTL8A*/*8B* to human diseases

The involvement of RTL8 has been implicated in two human diseases, ALS (Whiteley et al, 2021) and AS (Pandya et al, 2021) where RTL8 decreases in the former and accumulates in the latter. The situation in DKO is similar to the former, however, the DKO mice had a normal score in the rotarod test (Fig. 2K), suggesting that downregulation of these genes is not correlated with the ALS phenotype (Whiteley et al, 2021), although a TKO study will be required for a definitive conclusion. It is also of interest whether overexpression of human RTL8A-C together with overexpression of PEG10, another RTL gene (*RTL2*), is involved in the etiology of AS (Pandya et al, 2021). However, this will require another mouse model overexpressing *Rtl8a-c*.

It should be noted that the DKO phenotypes, the overgrowth due to increased food intake after 8w (Figs. 2A-F), as well as reduced social interaction (Fig. 2L) and increased apathy (Fig. 2M), are quite similar to those observed in the late PWS patients, who present with late-onset obesity and ASD-like symptoms from the juvenile age (Nicholls & Knepper, 2001; Tauber & Hoybye, 2021). In contrast to AS, which is caused by a paternal duplication of the chromosome 15q11-q13 and/or a UBE3A mutation (Nicholls & Knepper, 2001; Chamberlain & Lalande, 2010; Germain et al, 2020), PWS, another neurodevelopmental genomic imprinting disorder, is caused by its maternal duplication, resulting in overexpression of UBE3A. Thus, it is probable that RTL8A-C is reduced in the PWS patients, as it is in the DKO mice. It is then of interest to investigate whether the RTL8A-C protein is reduced in the PWS brain. To understand the etiology of late-onset obesity, further investigation of gene expression changes in the DKO hypothalamus is needed. It should be noted that in addition to the functional impairment of the hypothalamic regions, the medial orbital cortex (MO), a part of the orbitofrontal cortex (OFC), may explain the late-onset obesity, since the MO has been reported to play an important role in the regulation of feeding behavior (Seabrook & Borgland, 2020).

RTL8A-C is located on Xq26, far from the causative PWS chromosomal region (15q11-q13). Interestingly, however, a *de novo* paracentric inversion (X)(q26q28) with features mimicking PWS has been reported (Florez et al, 2003), suggesting the possibility that RTL8A-C may be responsible for the late PWS phenotypes. GABA_A_ receptor alterations have been reported in PWS patients, although it remains to be determined which α and β subunits are affected, i.e. α5 and β3 on chromosome 15q11-q13 (the PWS region) or α1 and β2 on chromosome 5q34-q35 or both (Lucignani et al, 2004). It will then be interesting to know whether *GABRB2* expression is actually downregulated in the PWS patients.

In addition to *RTL8A-C, RTL1* and *RTL4* have been implicated in neurodevelopmental disorders, such as KOS14/TS14 (Sekita et al, 2008; Kitazawa et al, 2021; Chou et al, 2022; Kagami et al, 2008; Kagami et al, 2015; Ioannides et al, 2014) and ASD (Lim et al, 2013; Irie et al, 2015). Eutherian-specific retroviral *Gag*-derived genes should therefore be the focus of more attention as important targets in human neurodevelopmental disorders.

## Materials and Methods

### Animals

All of the animal experiments were reviewed and approved by the Institutional Animal Care and Use Committee of RIKEN Kobe Branch, Tokai University and Tokyo Medical and Dental University (TMDU) and were performed in accordance with the RIKEN Guiding Principles for the Care and Use of Laboratory Animals, and the Guideline for the Care and Use of Laboratory Animals of Tokai University and TMDU.

### Estimation of the pairwise dN/dS ratio

Conversion of the protein sequence alignment created with the MAFFT program (https://mafft.cbrc.jp/alignment/server/index.html) into the corresponding codon alignment was performed with the PAL2NAL program (www.bork.embl.de/pal2nal/) (Suyama et al, 2006). The PAL2NAL program simultaneously calculated the nonsynonymous/synonymous substitution rate ratio (dN/dS) by the CodeML program (runmode: −2) in PAML (Yang, 2013). An aa sequence phylogenic tree was constructed with MEGA5 using the Maximum Likelihood method based on the JTT matrix based model (Tamura et al, 2011). The bootstrap consensus tree inferred from 1000 replicates is taken to represent the evolutionary history of the taxa analyzed. Branches corresponding to partitions reproduced in less than 50% bootstrap replicates are collapsed. The percentage of replicate trees in which the associated taxa were clustered together in the bootstrap test (1000 replicates) are shown next to the branches. Initial tree(s) for the heuristic search were obtained automatically as follows. When the number of common sites was < 100 or less than one fourth of the total number of sites, the maximum parsimony method was used; otherwise the BIONJ method with MCL distance matrix was used. The tree is drawn to scale, with branch lengths measured in terms of the number of substitutions per site. The analysis involved 10 aa sequences. All positions containing gaps and missing data were eliminated. There was a total of 104 positions in the final dataset. The *RTL8A, 8B* and *8C* genome sequences used for the dN/dS analysis (Table S1) were as follows: Human: NC_000023.11[c135052108-135051767, c135022459-135022118, 135032384-135032725]; Chimpanzee: NC_036902.1[c130245552-130245211, c130216735-130216394, 130226101-130226442]; Marmoset: NC_013918.1[122938486-122938827, c122959788-122959447, c123007085-123006744]; Mouse: NC_000086.8[c52645588-52645247, c52672265-52671924, 52610013-52610351]; Cow: NC_037357.1[18783020-18783361, 18794515-18794856, 18824715-18825056]; Rabbit: NC_013690.1[109509895-109510236, 109553752-109554093, 109580009 -10958035]; Dog: NC_049780.1 [106817683-106818024, 106834142-106834483, 106843724-106844065, 106859125-106859466]; Elephant: NW_003573520.1[4927338-4927679, c 4938179-4937838] and NW_003575145.1[c434-87].

### Subcellular localization of the mRTL8A and 8C in NIH3T3 cells

For overexpression of mRTL8A and 8C proteins, the ORF (open reading frame) of *mRtl8a* and *mRtl8c* together with the Kozak sequence, were amplified by PCR and cloned into pcDNA-3.1-myc His (Invitrogen) using *Nhe* I and *Apa* I restriction enzymes. The sequences were confirmed by sequencing. PCR was performed under the following conditions using the specified primers: reaction mixture (total 25 µl) contained 12.5 µl of 2 × KOD FX buffer, 5 µl of dNTP mix (2.0 mM each), 0.04 µl of forward and reverse primers (200 pmol/µl each), 1.0 µl of genomic DNA (5ng/µl), 0.5 µl of KOD FX polymerase (TOYOBO) and 6.42 µl of DDW, and the PCR condition was 1 cycle at 96°C for 45 seconds, followed by 35 cycles of 96°C for 15 seconds, 61°C for 30 seconds, and 72°C for 30 seconds. The following primers were used for *Rtl8a,* forward: 5’-gctagcgccatggaaggccaaggcaaggtaaaga-3’ and reverse: 5’-gggccccgggtgggaggaggatgaggacttc-3’ and for *Rtl8c*, forward: 5’-gctagcgccatggaccgccggattaagttgatta-3’ and reverse: 5’-gggcccgaagtcctcatcctcctcccacccg-3’. Transfection of the plasmid into NIH 3T3 cells was performed using polyethyleneimine (PEI) MAX (Polysciences, Inc.). Cell culture and transfection conditions were as follows: NIH 3T3 cells were cultured in DMEM (Dulbecco’s modified Eagle medium) supplemented with 10% FBS (fetal bovine serum) and 0.1% penicillin/streptomycin until reaching 70% confluence on the day before transfection. Prior to transfection, the cells were washed with PBS and then incubated with 350 µl of 10% FBS DMEM. The transfection mixture containing 1.16 µl of 1 mg/ml PEI MAX, 386 ng of plasmid DNA, and 35 µl of 1.5M NaCl was added to the cells. After 24 hours of incubation, the cells were used for Western blot and immunostaining. The cells were washed with PBS, fixed with 4% paraformaldehyde, permeabilized with 0.1% Triton X-100, and blocked with 10% goat serum albumin, then incubated with primary antibody followed by secondary antibody conjugated to Alexa Fluor 488. Nuclei were stained with DAPI and images were captured using a confocal microscope.

### Production of RTL8A and RTL8C chimeric proteins

The ORF sequences used for RTL8A and 8C overexpression were ligated into pcDNA-3.1-myc His after digestion with *Pci* I restriction enzyme. Chimera constructs were then selected by sequencing. Overexpression and immunostaining were performed using the same methods as for RTL8A and RTL8C overexpression.

### RT-PCR

RT-PCR was performed using cDNA. cDNA was synthesized from 1 μg of total RNA using SuperScript Ⅲ Reverse Transcriptase (Invitrogen). RNA was extracted from adult tissues at 19 weeks, including the cerebrum, cerebellum, heart, liver, lung, spleen, kidney, stomach, intestine, large intestine, skeletal muscle, thymus, bladder, adipose, uterus, ovary, testis, epididymis and vesicular gland as well as day 9.5 placenta by treatment with TRIzol Reagent (Life Technologies). Ten ng of cDNA were mixed with 1 x ExTaq buffer (TAKARA), 2.5 mM of each dNTP 2 μl, primer 0.2 μl (200 pmol/μl) and ExTaq HS 0.1 μl (TAKARA), and PCR analysis was performed under the following conditions. *Rtl8a, b* and *c*: 95℃ 1min, 32 (*Rtl8a* and *b*) and 31 (*Rtl8c*) cycles at 96℃ for 10 s, 72℃ for 30 s and 72℃ for 90 s. *β-actin*: 95℃ 1min, 26 cycles at 96℃ for 10 s, 72℃ for 30 s, and 72℃ for 90 s using a C1000 Touch thermal cycler (Biorad). The following primer sets were used. *Rtl8a, b* F1: tcccactgttgacagctcag and *Rtl8a, b* R1: ggggctactgttggaaaa, and *Rtl8c* F1: tatgcgctatctaaagacagac and *Rtl8c* R1: ggcttctgtacaggtagaggcagac. The RFLP (restriction fragment length polymorphism) experiment was carried out using *Msp*I, and the results were confirmed by electrophoresis.

### Generation of *Rtl8c* flox mice

A targeting construct spanning the two exons of *Rtl8c* was used to generate *Rtl8c* flox mice (Accession No. CDB0788K: https://large.riken.jp/distribution/mutant-list.html). Briefly, the construct included two loxP sites upstream of exon 1 and downstream of exon 2, and a frt-neo-frt cassette downstream of the latter loxP site. After linearizing, the construct was electroporated into TT2 ES cells (C57BL/6 × CBA genetic background) (Yagi et al, 1993). Two ES clones were then screened by PCR and Southern blotting. One 5′ and one 3′ probe, both of which were outside of targeting construct, were used in Southern blotting. The correct targeting clones were then used to generate chimera founder lines. Germline transmission was observed in two chimeric mice.

The *Rtl8c* flox alleles were checked by electrophoresis of the PCR amplification using the following primers. WT forward: GGCGACAACCAAGGTTTTTA and reverse:

GGGTCTGCTTCTCTTTGCTG, flox allele forward: AAATAGGCGTATCACGAGGC and reverse: GGGTCTGCTTCTCTTTGCTG. The PCR cycle was 96℃ 1min, 35 cycles at 96℃ for 15 s, 60℃ for 30 s and 72℃ for 30 s. and 72℃ 2min.

### Generation of *Rtl8a/Rtl8b* DKO mice

The DKO mice were generated by referring to our previous report (Ono et al, 2015) as follows. The plasmids expressing *hCas9* and sgRNA were prepared by ligating oligos into the *Bbs*I site of pX330 (Addgene plasmid #42230). The 20 bp sgRNA recognition sequences was: *Rtl8a, b*-sgRNA (5′-ACGGGATGGGGTTCCGCCGA-3′). The sgRNA targets both *Rtl8a* and *Rtl8b*.

To produce the *hCas9* mRNA, the T7 promoter was added to the *hCas9* coding region of the pX330 plasmid by PCR amplification, as previously reported (Wang et al, 2013). The T7-*Cas9* PCR product was gel purified and employed as the template for *in vitro* transcription (IVT) using a mMESSAGE mMACHINE T7 ULTRA kit (Life Technologies). The T7 promoter was added to the *Rtl8a, b*-sgRNA region of the pX330 plasmid by PCR amplification using the following primers. *Rtl8a, b*-sgRNA-F (5′-TGTAATACGACTCACTATAGGGACGGGATGGGGTTCCGCCGA-3′), *Rtl8a, b*-sgRNA-R (5′-AAAAGCACCGACTCGGTGCC-3′). The T7-sgRNA PCR product was gel-purified and employed as the template for IVT using a MEGAshortscript T7 kit (Life Technologies). Both the *hCas9* mRNA and *Rtl8a, b*-sgRNA were DNase-treated to eliminate template DNA, purified using a MEGAclear kit (Life Technologies), and eluted into RNase-free water.

C57BL/6N female mice were superovulated and *in vitro fertilization* was carried out using *Rtl8c* flox mouse sperm. The synthesized *hCas9* mRNA (50 ng/μl) and *Rtl8a, b*-sgRNA (25 ng/μl) with oligo DNA (50 ng/μl) were injected into the cytoplasm of fertilized eggs at the indicated concentration. The eggs were cultivated in KSOM overnight, then transferred into the oviducts of pseudopregnant ICR females. The oligo DNA designed as the HDR (homology-directed repair) donor induced a nonsense mutation in both *Rtl8a* and *Rtl8b* and harbored an accompanying *Afl*II recognition site used for PCR-RFLP (Restriction Fragment Length Polymorphism) genotyping. The sequence of the oligo DNA was: 5′-AAGGCCAAGGCAAGGTAAAGAGGCCGAAGGCCTACATGCTCAGGCACAACAGGCGGCGCCCTTAAGGGAACCCCATCCCGTTTCCAGAGCTGTTTGATGGCGAGATGGACAAGCTCCCGGAGTTCA-3′.

### Genotyping of *Rtl8a* and *8b* DKO mice

The genotype was determined by PCR-RFLP analysis. The following primers were used for PCR amplifications, *Rtl8a*: 5′-GGACTGGCGCCTGAAATAGC-3′ and 5′-GCACAATCACCACCTCTTGAACA-3′, *Rtl8b*: 5′-CCACCCCTTAAACATTCTCCTGG-3′ and 5′-AGATCGAACATCAGGCCATGAAC-3′. The PCR products were digested with *Afl*II and subjected to agarose gel electrophoresis.

### Behavioral Analysis

Behavioral analysis was performed as previously described (Irie et al, 2015; Kitazawa et al, 2021). Briefly, both DKO and littermate WT male mice (8-20 w) were analyzed in the following order, Light and Dark transition, Open field, Social Interaction, Elevated plus maze, Rotarod test, Tail suspension and Fear condition tests.

### Light and Dark transition test

The apparatus used for the light/dark transition test comprised a cage (21 x 42 x 25 cm) divided into two sections of equal size by a partition with a door (O’hara & Co., LTD.). One chamber was brightly illuminated (400 lux), whereas the other chamber was dark. Each mouse was placed into the dark side and allowed to move freely between the two chambers with the door open for 10 min.

The total number of transitions between the chambers, the time spent in each, the latency to first enter the light chamber, and the distance travelled in each chamber were recorded using a video camera attached to a computer and calculated by the Image LD program.

### Open field test

Locomotor activity was measured using an open field test. Each mouse started the test in the corner of the open field apparatus (40 x 40 x 30 cm; Accuscan Instruments, O’hara & Co., LTD.). The chamber of the test was illuminated at 100 lux. Total distance (cm), total move time, and time spent in the center area were recorded.

### Social interaction test

Sociability was measured using the open field apparatus. Mice of the same genotype were placed at diagonal corners in an open field apparatus in small baskets. After waiting for 1 minute, the behavior was recorded for 10 minutes after the mice exited from the baskets at the same time. The time to first contact, the total number of contacts, and the total time of sociable behavior were analyzed. All data was analyzed by a blind observer. The total sociable behavior time was measured excluding contact of less than one second. The test was performed with different pairs for a period of 2 days, and the average value for the 2 days was calculated. Mice were paired such that the weight difference was 2 g or less.

### Elevated plus maze test

The elevated plus maze consisted of two open arms (25 x 5 cm) and two closed arms of the same size with 15 cm high transparent walls (O’hara & Co., LTD.). The behavior testing room (170 x 210 x 200 cm) was soundproof, and the illumination level was maintained at 100 lux. Each mouse was placed in the central square of the maze with its head pointed toward a closed arm. Data were recorded for 10 min using a video camera attached to a computer and were calculated with the Image EP program. The number of entries into each arm, total arm entries and the time spent in the open arms were recorded.

### Rotarod test

Balance and motor ability were confirmed by the rotarod (BrainScience.idea.co., ltd). After placing the mouse in the rotating lane, the speed was changed from 3 to 30 rpm and the time until the mouse fell was measured. When one half of the body of the mouse was detached from the rods, it was measured as if it had fallen. The test was conducted for 3 consecutive days and 3 times a day.

### Tail suspension test

Depression was investigated using a tail suspension test. The device used was the same as the fear test described next. The mouse was hung a hook with tape, and observed for 10 minutes. Recordings were made with a camera attached to the device and the freezing percentage was measured with software (O’hara & Co., LTD.). Freezing percentages were compared during the time elapsed excluding the first 3 minutes.

### Fear condition test

On the 1st day, each mouse was placed into a test chamber (26 x 34 x 29 cm) with a stainless-steel grid floor inside a sound-attenuated chamber and allowed to explore freely for 120sec. A 60 db white noise acting as a conditioned stimulus (CS) was presented for 30 seconds, followed by a mild foot shock (2 seconds, 0.5 mA) acting as an unconditioned stimulus (US). Two more CS-US pairings were given at a stimulus interval of 2 minutes. On the 2nd day, a context test was performed in the same chamber as the conditioning test and data were recorded for 2 min. On the 3rd day, a cued test was performed in another testing chamber. The same 60 db white noise as on the 1st day was given for 30 s and data were recorded for 2 min. The entire test was recorded by a video camera attached to a computer. In each test, the movement freezing percentage and total move distance were calculated automatically by the Image FZ program.

### Immunohistochemistry

Mouse adult brains were fixed in 4% paraformaldehyde (PFA: Nacalai tesque), incubated in 10% and 25% sucrose at 4℃ overnight each and finally embedded in OCT compound (Sakura Finetek). The OCT blocks were sectioned at a 13-μm thickness with a cryostat (MICROTOME) and mounted on Superfrost Micro Slides (Matsunami Glass). The cryosections were fixed in 4% PFA for 10 min at room temperature and washed three times with PBS for 5 min. For antigen retrieval, the sections were boiled in 0.01 M Citrate Buffer pH 4.0 at 98℃ for 30 min (for RTL8A/8B), and then immersed (dehydrated) in cold methanol at −30℃ 30min. After being air dried, the sections were blocked with 10% goat serum, 1% bovine serum albumin (BSA: Sigma Aldrich), and 0.1% Triton-X 100 (WAKO) in PBS at room temperature for 1 hr.

For the staining, an anti-RTL8A/8B antibody (SCRUM 1:1000), was used as the primary antibodies and were prepared in 1% BSA and 0.1% Triton-X 100 in PBS at 4℃ overnight. The second antibody was Biotin-αRabbit-IgG (VECTOR STAIN 1:200).

Nuclei were stained with Nuclear fast red. The slides were mounted with malinol (MUTO Pure Chemicals, Tokyo, Japan). The images were captured using a BIOREVO microscope (KEYENCE).

### Immunofluorescence staining using floating frozen section

Mouse brains were fixed and embedded in OCT blocks as described in the immunochemistry section. The OCT blocks were thinly sliced at 50 µm, floated in a glass bottom dish with 2 ml of PBS and washed for 10 min. PBS was removed and 2 ml of 0.1 M PBS/0.3% Triton was added and washed for 15 min three times. The sections were blocked with 10% donkey serum, 5% bovine serum albumin (BSA: Sigma Aldrich), and 0.1% Triton-X 100 (WAKO) in PBS at room temperature for 1 hr. For the staining, an anti-RTL8A/8B (SCRUM 1:2000) and anti-MAP2 (ab5392 1:20000) antibodies were used as the primary antibodies and were prepared in 5% BSA and 0.1% Triton-X 100 in PBS at 4℃ two overnight. Second antibodies used were the Alexa Fluor 488 donkey anti-Mouse IgG (H+L) (Life technologies, A21203,1:2000) and Alexa Fluor 488 donkey anti-Rabbit IgG (H+L) (Jackson ImmunoResearch, 711-545-152, 1:2000) and were prepared in 5% BSA and 0.1% Triton-X 100 in PBS at 4℃ overnight. Nuclei were stained with DAPI (VECTOR STAIN, 1:1000). The slides were mounted with VECTERSHIED Hardset antifade mounting medium (VECTOR STAIN). The images were captured using a confocal laser microscope TCS SP8 (Leica).

### Immunofluorescence staining using paraffin embedded samples

Mouse adult brains were fixed overnight using 4% paraformaldehyde (PFA: Nacalai tesque), and soaked in 70% ethanol at 4℃ overnight, then dehydrated in 70%, 80%, 90% ethanol for 1 hr each, 100% ethanol for 2 hr twice and 3hr once. They were embedded in paraffin wax after treatment with Gnox (Genostaff, GN04) for 2 hr twice, Gnox and paraffin for 2 hr, and paraffin for 3 hr and 4.5 hr. The paraffin blocks were sectioned at a 5μm thickness with a microtome and mounted on Superfrost Micro Slides (Matsunami Glass). For antigen retrieval, the sections were boiled in 0.01 M Citrate Buffer (pH 6.0) at 90℃ for 15 min then immersed (dehydrated) in cold methanol at - 30℃ for 30min. After being air dried, the sections were treated with 10% goat serum, 1% bovine serum albumin (BSA: Sigma Aldrich), and 0.1% Triton-X 100 (WAKO) in PBS at room temperature for 1 hr.

For the immunofluorescence staining, anti-MAP2 (Proteintech 17490-1-AP 1:200) antibody was reacted as the primary antibodies in 1% BSA and 0.1% Triton-X 100 in PBS at 4℃ overnight (approximately 16 hr). Second antibodies used were the Alexa Fluor 488 donkey anti-Rabbit IgG (H+L) (Jackson ImmunoResearch, 711-545-152, 1:1000). Nuclei were stained with DAPI (VECTOR STAIN, 1:1000). The slides were mounted with VECTERSHIED Hardset antifade mounting medium (VECTOR STAIN). Neuronal areas were measured by a blind observer using ImageJ. The neuronal cells and nuclei were outlined manually with “Polygon selections” tool. The area of MAP2-positive neurons and the area of the nucleus were measured. Cytoplasmic area was calculated by subtracting nuclear area from cell area.

### RNA-Seq

For RNA isolation and quality control, the integrity and quantity of the total RNA was measured using an Agilent Technologies 2200 TapeStation (Agilent Technologies, Inc., Santa Clara, CA). For library preparation and sequencing, total RNA obtained from each sample was subjected to a sequencing library construction using the TruSeq Stranded mRNA Library Prep Kit (Illumina, Inc., San Diego, CA) according to the manufacturer’s protocols. Sample libraries were sequenced using NovaSeq 6000 (Illumina, Inc., San Diego, CA) in 100-base pair (bp) paired-end reads. For alignment to the whole transcriptome, quantification of gene expression levels and detection of differentially expressed genes, sequencing adaptors, low-quality reads, and bases were trimmed using the Trimmomatic-0.39 tool (Bolger et al, 2014). Sequence reads were aligned to the mouse reference genome (mm10) using STAR 2.7.9a (Dobin et al, 2013). The aligned reads were subjected to downstream analyses using StrandNGS 4.0 software (Agilent Technologies, Inc.). Read counts assigned to each gene and transcript (Ensembl Database 2018.02.25) were quantified using an iDEP 1.12 (Ge et al, 2018) with the DESeq2 package. Finally, the gene set enrichment analysis was performed using this platform with GO cellular component gene sets (FDR < 0.05). The tree plot and network diagram were classified according to the GO cellular component process collection.

### Western blot

Membrane proteins were extracted from mouse brain using the Thermo Scientific Mem-PER Plus Membrane Protein Extraction Kit (Thermo: 89842). The extracted proteins were boiled in 4X sample buffer at 95°C for 3 minutes. Electrophoresis was then performed on a 4–20% gradient gel (BIO-RAD: #4561095) at 100V for 40 minutes.

After electrophoresis, the gel was transferred to membrane (BIO-RAD: #1704156) and blocked overnight with 5% skim milk in TBS. Primary antibodies, GABRB2 (Sigma: ZMS1130 1/1000) and GAPDH (1/20000 Proteintech:10494-1-AP), were incubated with the membrane overnight at 4°C. Secondary antibodies, anti-mouse (Proteintech:SA00001-2, 1/2000) and anti-rabbit (Proteintech: SA00001-2, 1/2000), were then incubated for 2 hours at room temperature. The reaction was visualized by adding detection reagent (GE Healthcare: 28980926), and images were captured using a gel documentation system (ImageQuant™ LAS 4010(4000). Protein expression levels were measured using the imageJ gel tool and corrected for GAPDH levels.

## Acknowledgments

The authors would like to thank Ms. Hitomi Takahashi (TMDU) for technical assistance in the embryo transfer experiment and Drs. Moe Kitazawa and Ayumi Matsuzawa (TMDU) for providing advice on the histological and bioinformatics experiments. Pacific Edit reviewed the manuscript prior to submission.

## Author contributions

Y.F., M.I., H.S. T.E. Y.H. M.K. and F.I. performed the experiments and analyzed the data. M.I., R.O. H.K. and MK generated *Rtl8c*flox mice and R.O. and H. S. generated *Rtl8a* and *Rtl8b* DKO mice. F.I. and T.K.-I. designed the study and Y.F., T.K.-I. and F.I. wrote the manuscript. All authors agree to be accountable for the content of the work.

---

**Fig. S1.**
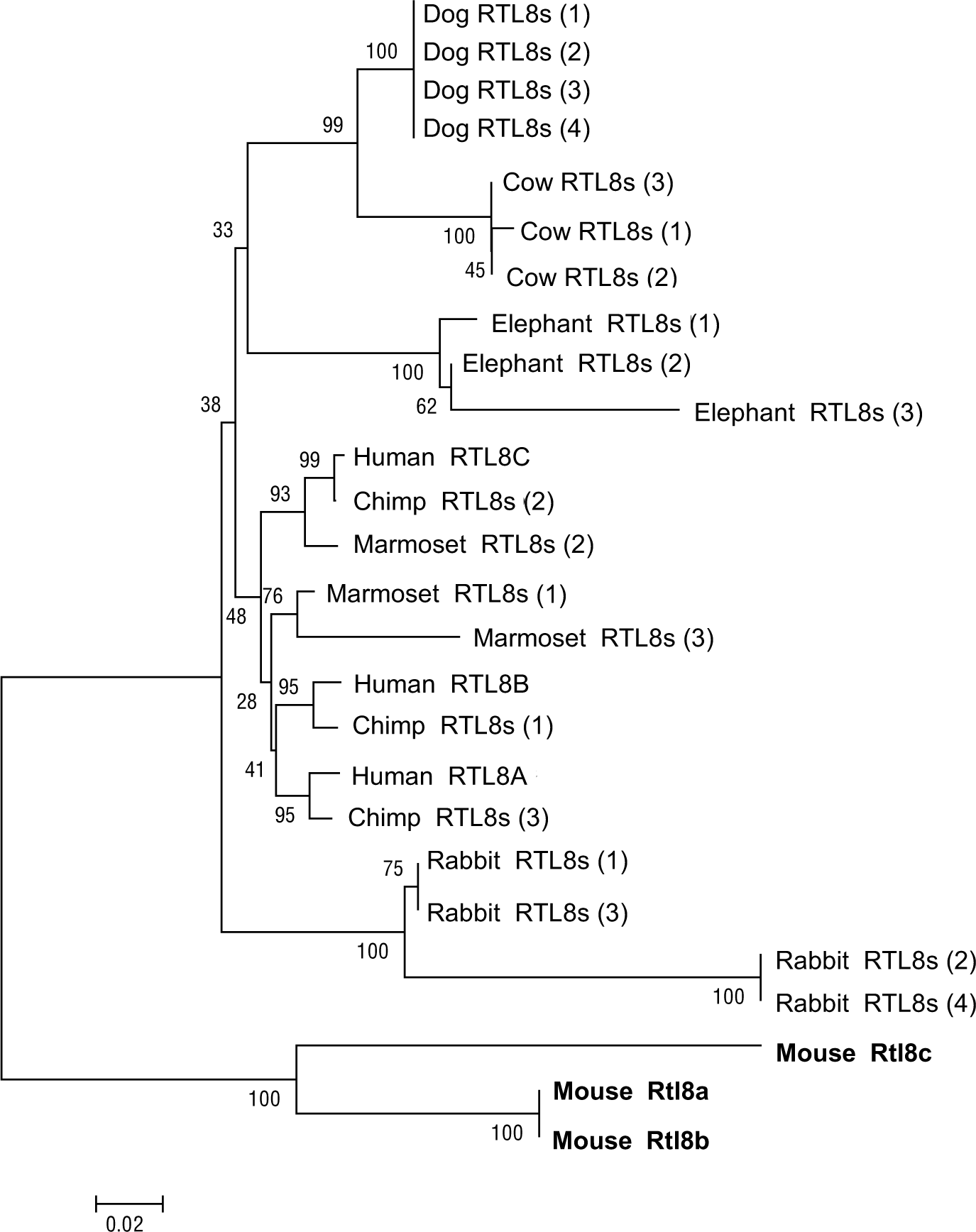
Evolutionary status of the *RTL8s* in eutherians. In primates, human *RTL8A*, *8B* and *8C* are orthologous to Chimp *RTL8s* (1)-(3), but only marmoset *RTL8s* (2) is orthologous to human *RLT8C* and chimp *RTL8s* (2). However, in most cases, *RTL8*s within each species form a cluster, such as dog, cow, elephant, rabbit and mouse, indicating a higher homology to each other than to those in other species. Therefore, mouse *Rtl8a* is not an ortholog of human *RTL8A*, and the same it true for *Rtl8b* and *Rtl8c*. The evolutionary history was inferred by using the Maximum Likelihood method based on the JTT matrix-based model, MEGA5.

**Fig. S2.**
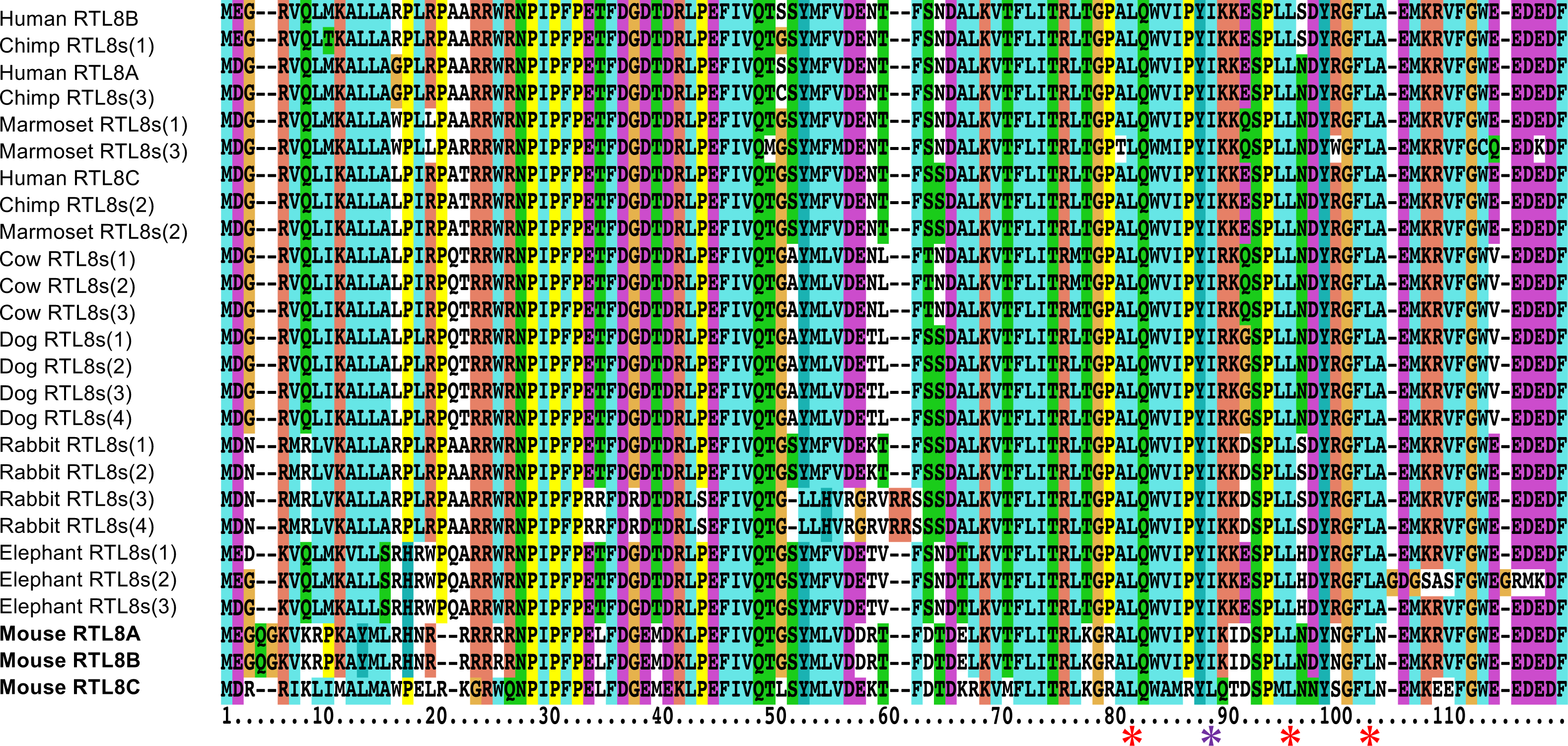
Comparison of amino acid sequences of RTL8A-C proteins in eutherians. A leucine zipper motif (3 red asterisks and 1 purple asterisk under the aa sequence is only present in mouse RTL8C (the bottom line) because the second leucine (purple) is replaced with isoleucine in the other RTL8A-C. Mouse RTL8A and 8B are unique because their N-terminal sequences contain the nuclear localization peptide-like sequences (see also Fig. 1B).

**Fig. S3.**
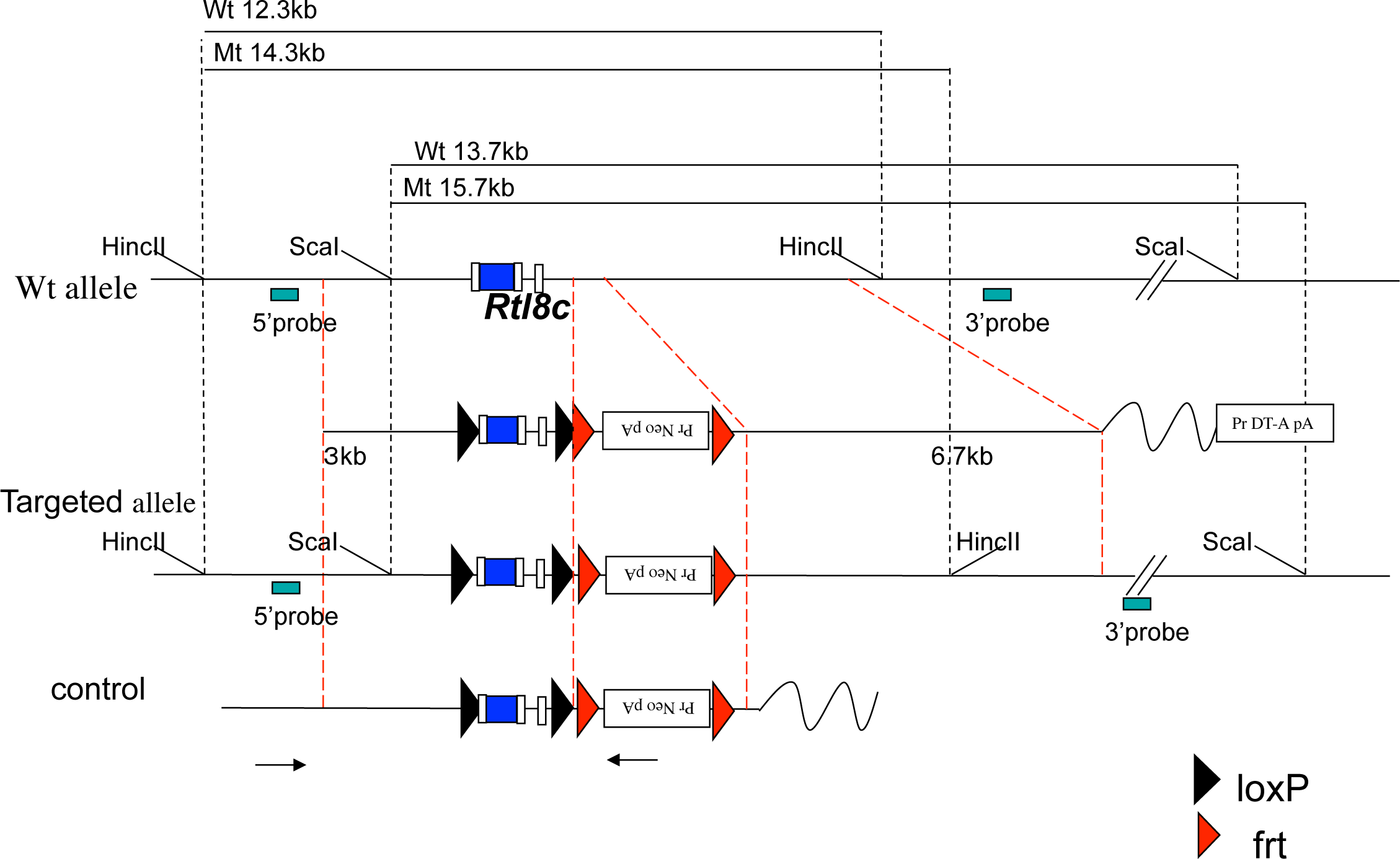
Construction of *Rtl8c* flox mouse. The *Rtl8c* flox targeting construct was generated using three genomic fragments, the 5′-arm (3.0 kb), middle arm containing *Rtl8c* exons and 3′-arm (6.5 kb). Two loxP sites were inserted upstream of exon 1 and downstream of exon 2, a frt-neo-frt cassette was inserted downstream of the latter loxP site. After linearizing, the construct was electroporated into TT2 ES cells. ES cells in which homologous recombination had occurred were injected into 8-cell stage embryos. Germ line transmission of the *Rtl8c* flox with the neomycin cassette (*Rtl8c* flox MT) was confirmed by Southern blot and PCR using the genome prepared from pups in which male *Rtl8c* flox chimeric mice had been crossed with female C57BL/6J. To remove the neo cassette, we mated the *Rtl8c* flox MT mice with CAG-FLP TG mice. The two white boxes represent *Rtl8c* exons and blue box in the exon 1 represent RTL8C ORF. The black and red arrowheads represent loxP and frt sites, respectively. The black arrows represent the PCR primers for screening of the targeted ES clones.

**Fig. S4.**
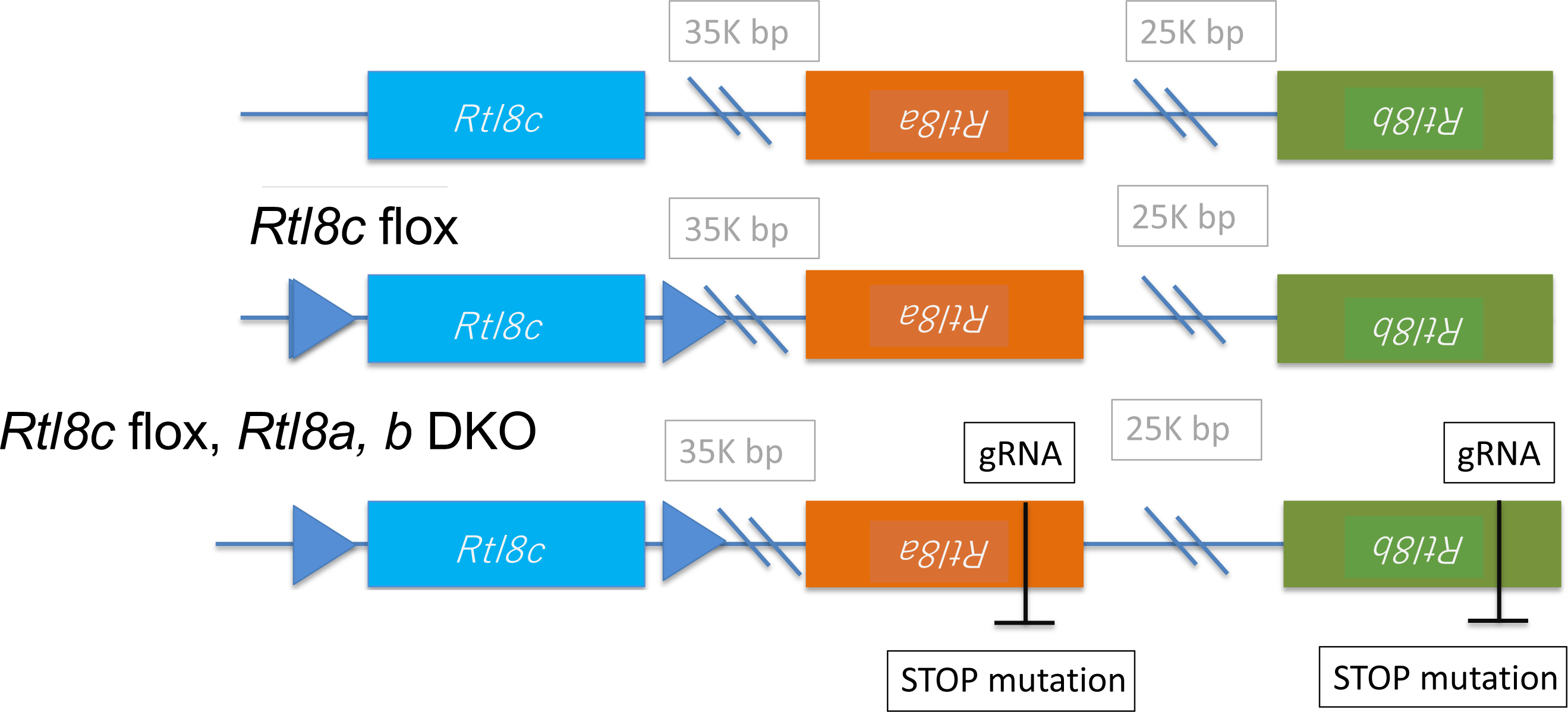
Strategy for making *Rtl8a* and *Rtl8b* DKO mouse. *Rtl8a* and *Rtl8b* are located tandemly while *Rtl8c* is reversed on the mouse X chromosome. First, *Rtl8c* flox mice were generated using TT2 ES cells (see Materials and Methods), then, *Rtl8a* and *Rtl8b* DKO mice were generated by integration of a stop codon in both *Rtl8a* and *Rtl8b*, as shown by the black bars, using CRIPR/Cas9.

**Fig. S5.**
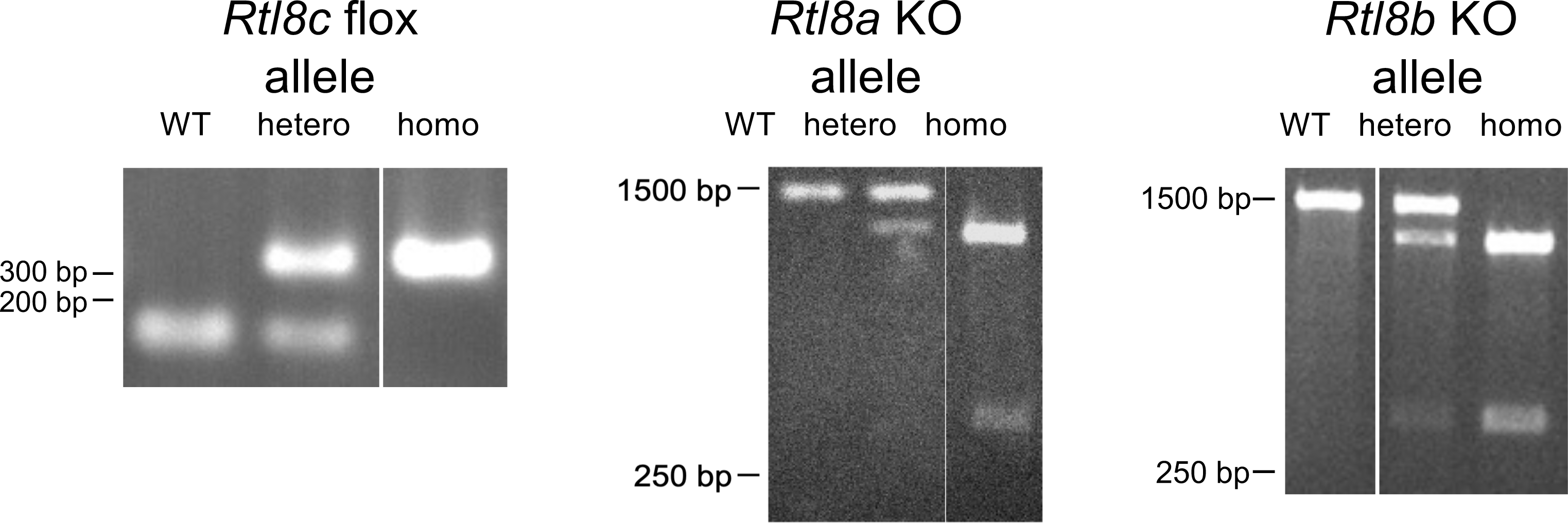
Genotyping the *Rtl8c* flox mice and the *Rtl8a* and *Rtl8b* DKO mice. Left: WT and *Rtl8c* flox alleles were detected as 169 and 349 bp bands in the PCR experiment, respectively. Middle: A *Rtl8a* KO allele with a stop mutation gave the same 1502 bp bands as WT, but were further digested by *Afi*II to give 1170 and 332 bp bands. Right: A *Rtl8b* KO allele with a stop mutation gave the same 1542 bp bands as WT, but were further digested by *Afi*II to give 1186 and 356 bp bands.

**Fig. S6.**
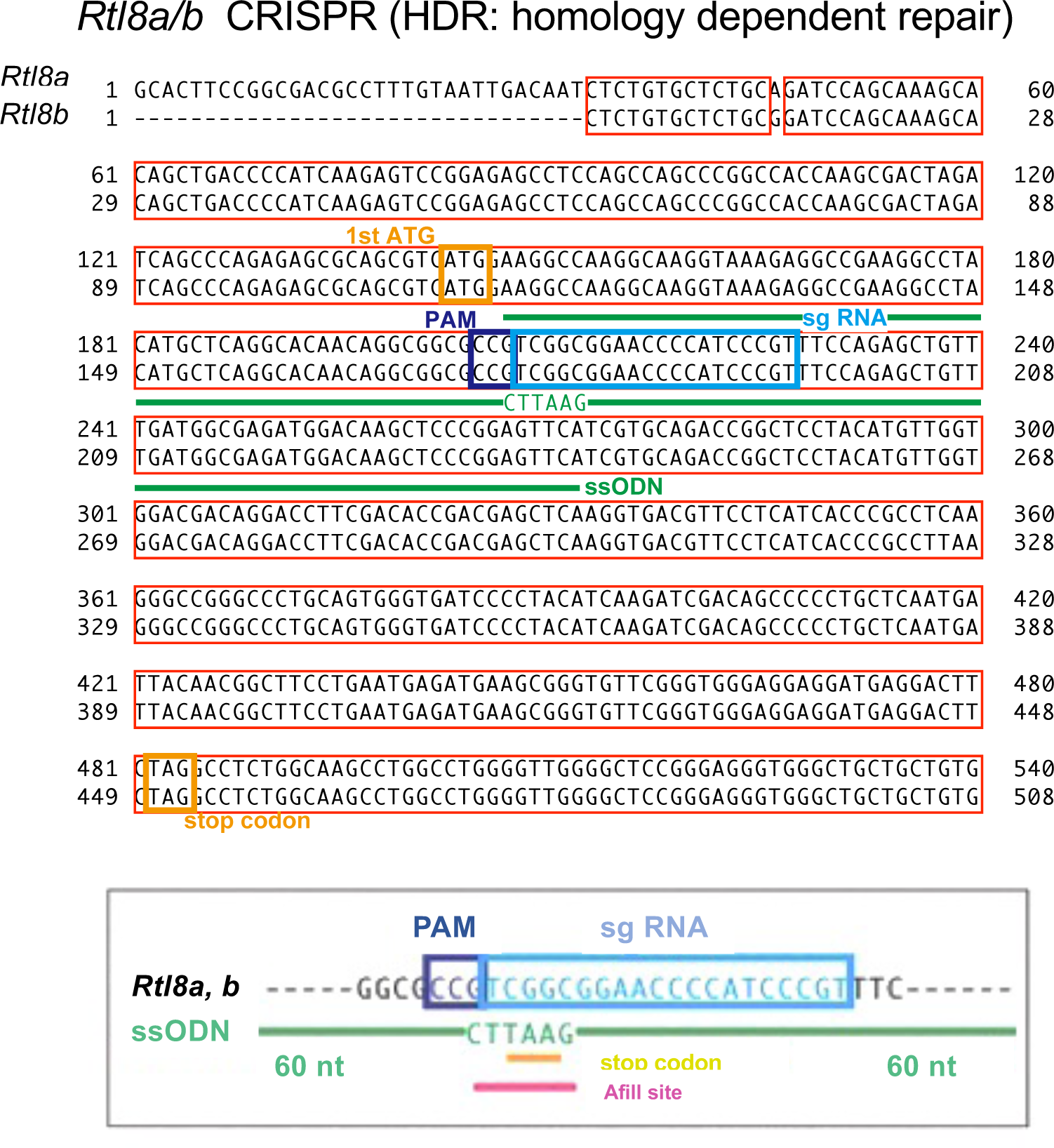
Construction of *Rtl8a* and *8b* DKO mouse. A stop codon was integrated in both *Rtl8a* and *Rtl8b* genes using the CRISPR/Cas9 method. The red and yellow boxes indicate the gene sequences of *Rtl8a* and *Rtl8b* as well as the start and stop codons, respectively. The light blue and purple box indicate the sgRNA and sPAM sequences, respectively. Pink line indicates the *Afl*II recognition site. ssODNs: single-stranded oligodeoxynucleotides.

**Fig. S7.**
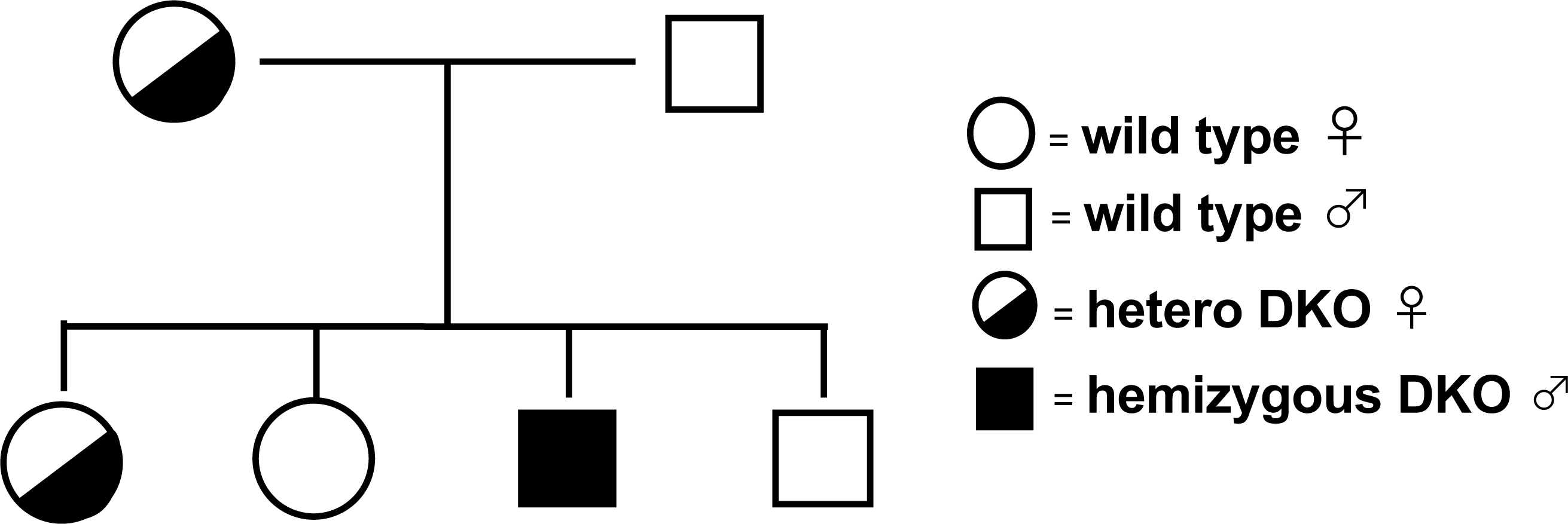
Cross diagram of *Rtl8a* and *8b* DKO and wild type mice. As *Rtl8a* and *Rtl8b* are X-linked genes, hemizygous DKO males exhibited a null phenotype but heterozygous females exhibited different phenotypes because one of the two X chromosomes is inactivated.

**Fig. S8.**
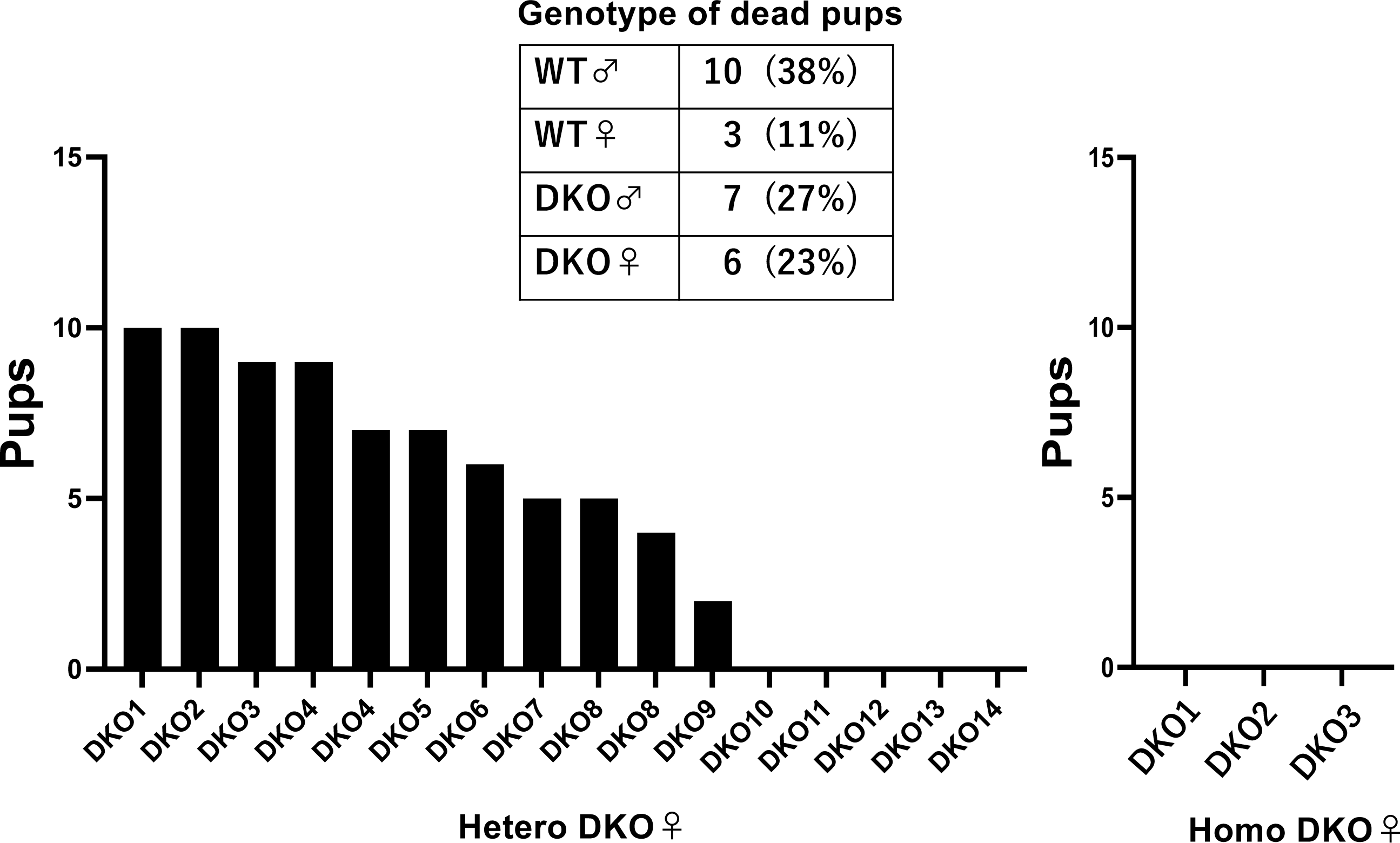
One half of the hetero DKO mothers failed to take proper care of their pups. Left: In most cases, hetero DKO mothers delivered 6-10 pups, but most pups were lost in half of the cases while most were viable in the other half. DKO10-DKO14 dead pups included pups with WT and DKO genotypes in approximately equal proportions (top right table). Right: In the case of homo DKO mothers, no pups survived.

**Fig. S9.**
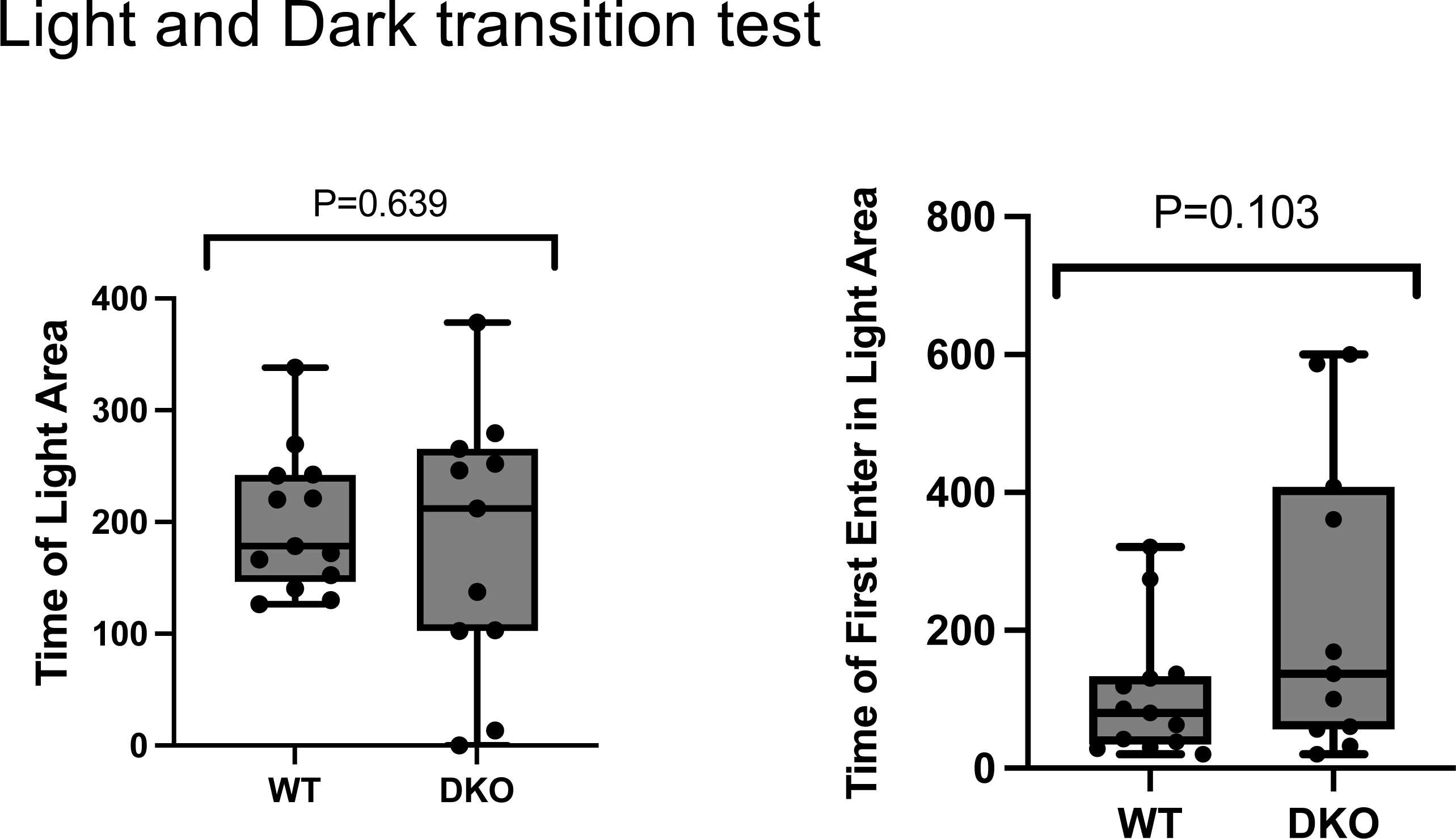
Light/Dark transition test. Results of Light/Dark transmission test except for those already shown in Fig. 2B. WT: n = 14, DKO: n = 11.

**Fig. S10.**
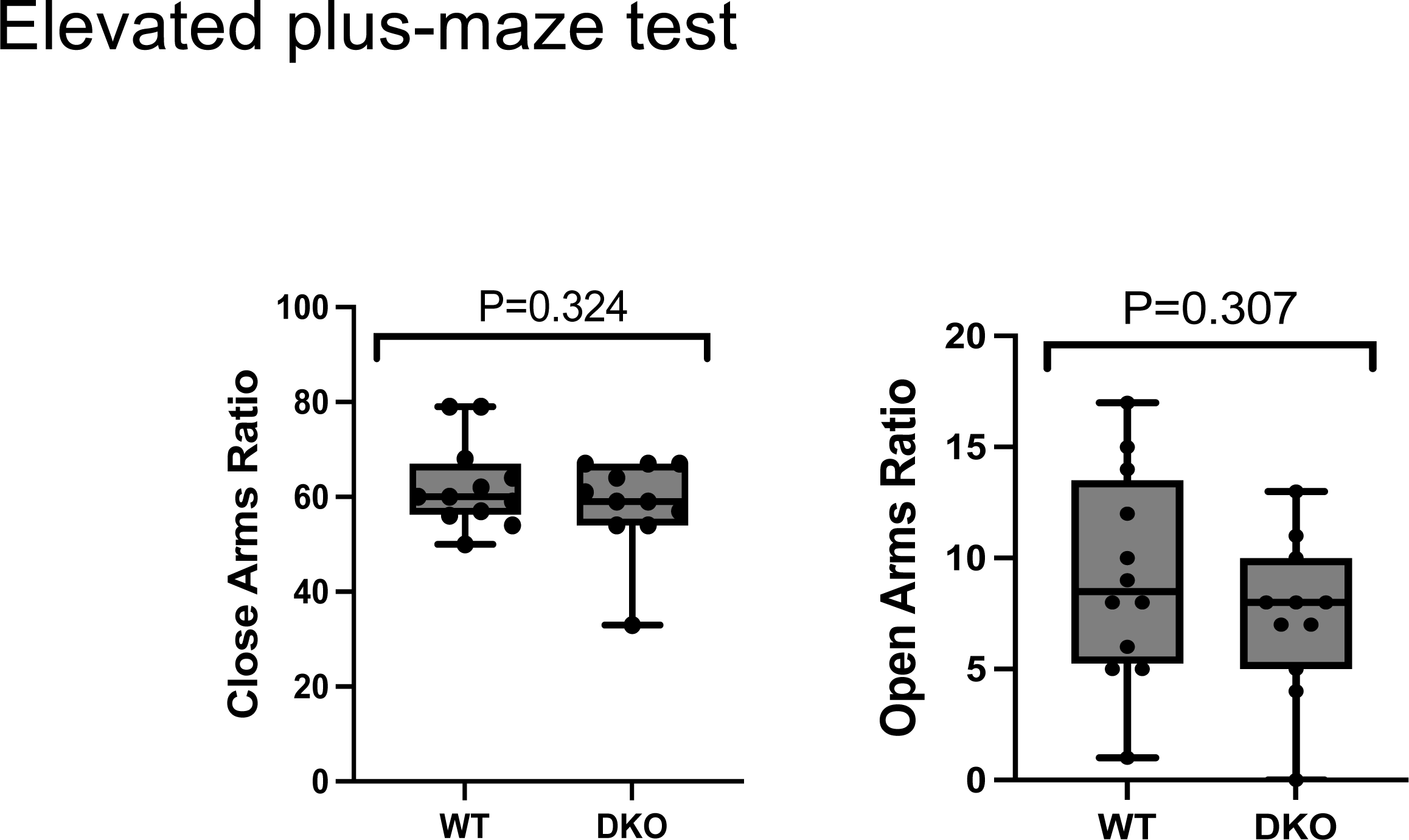
Elevated plus-maze test. Results of Elevated plus-maze test except for those already shown in Fig. 2B. WT: n = 12, DKO: n = 11.

**Fig. S11.**
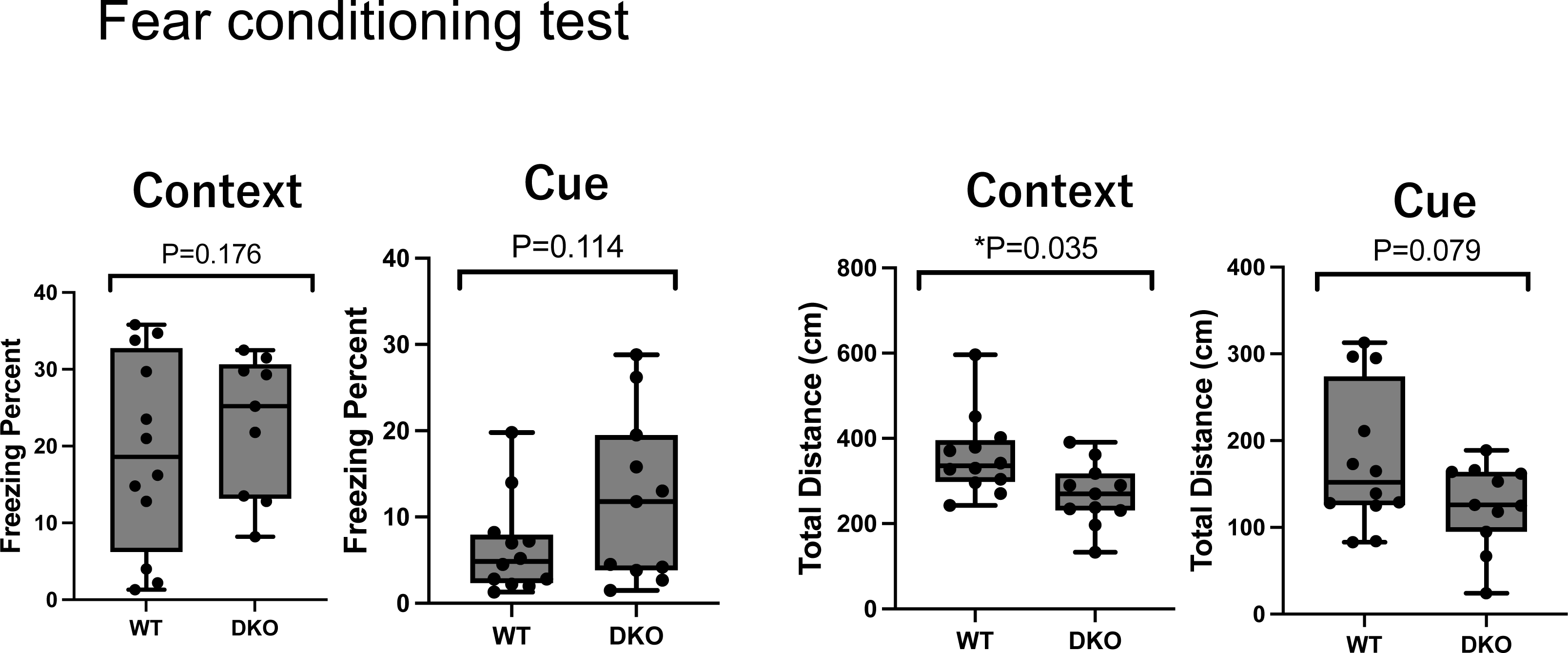
Fear conditioning test. Results of the Fear conditioning test are shown, WT: n = 12, DKO: n = 11.

**Fig. S12.**
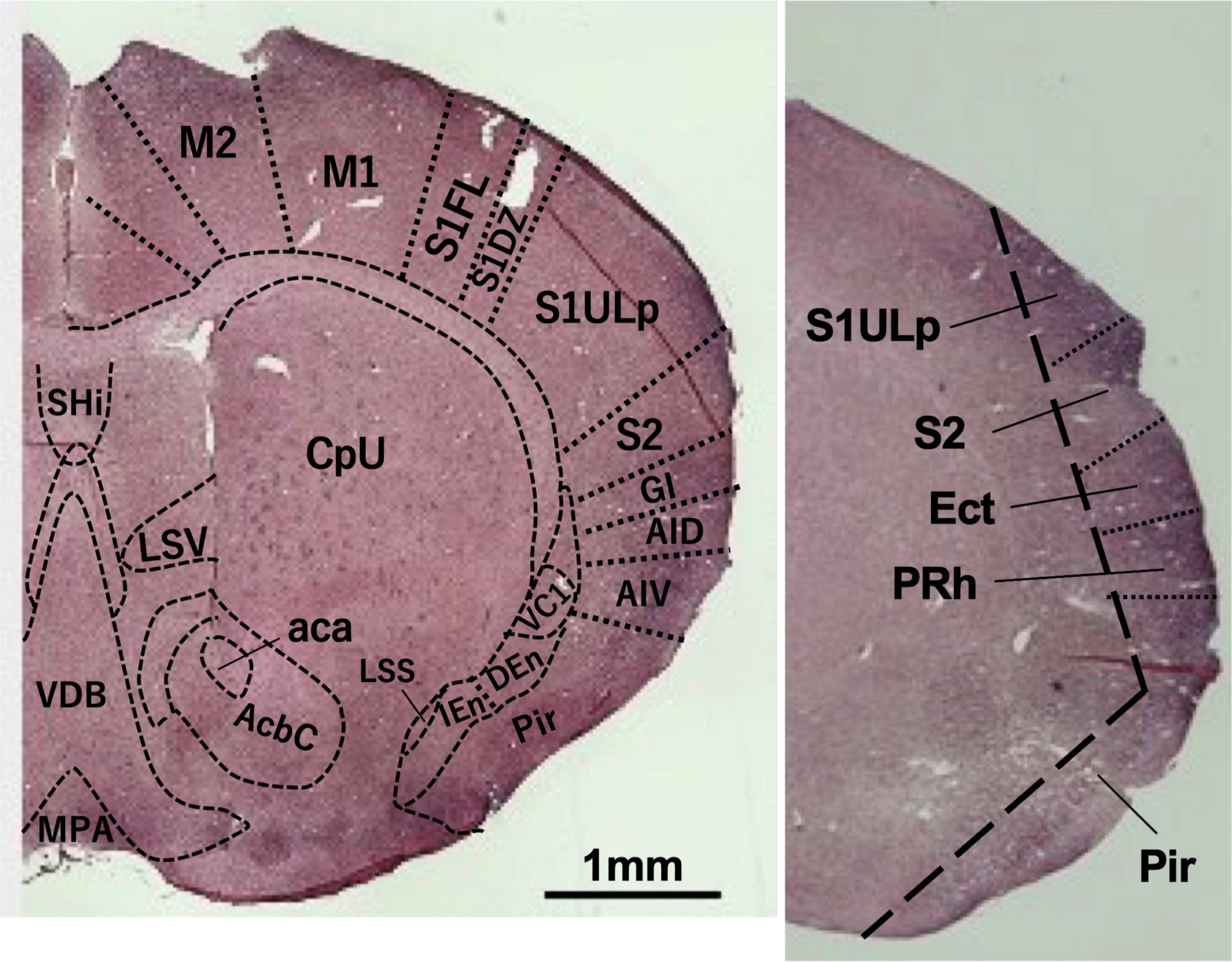
Immunostaining of RTL8A and RTL8B proteins in the striatum and temporal lobe. Coronal section images of WT mice showing staining in the striatum region (left: 10 w) and temporal lobe (right: 20 w). aca: anterior commissure, anterior part, AcbC: accumbens nucleus, core (ventral striatum), AID: agranular insular cortex, dorsal part, AIV: agranular insular cortex, ventral part, CpU: caudate putamen (dorsal striatum), DEn: dorsal endopiriform claustrum, Ect: ectorhinal cortex, GI: granular insular cortex, IEn: intermediate endopiriform claustrum, MPA: medial preootic area, M1: primary motor cortex, M2: secondary motor cortex, LSS: lateral stripe of the striatum (ventral stiatum), LSV: lateral septal nucleus, ventral part, Pir: piriform cortex, PRh: perirhinal cortex, S1DZ:primary somatosensory cortex, dysgranular zone, S1FL: primary somatosensory cortex, forelimb region, S1ULp: primary somatosensory cortex, upper lip region, S2: secondary somatosensory cortex, SHi: septohippocampal nucleus, VC1: ventral part of claustrum, VDB: nucleus of the vertical limb of the diagonal band. c

**Fig. S13.**
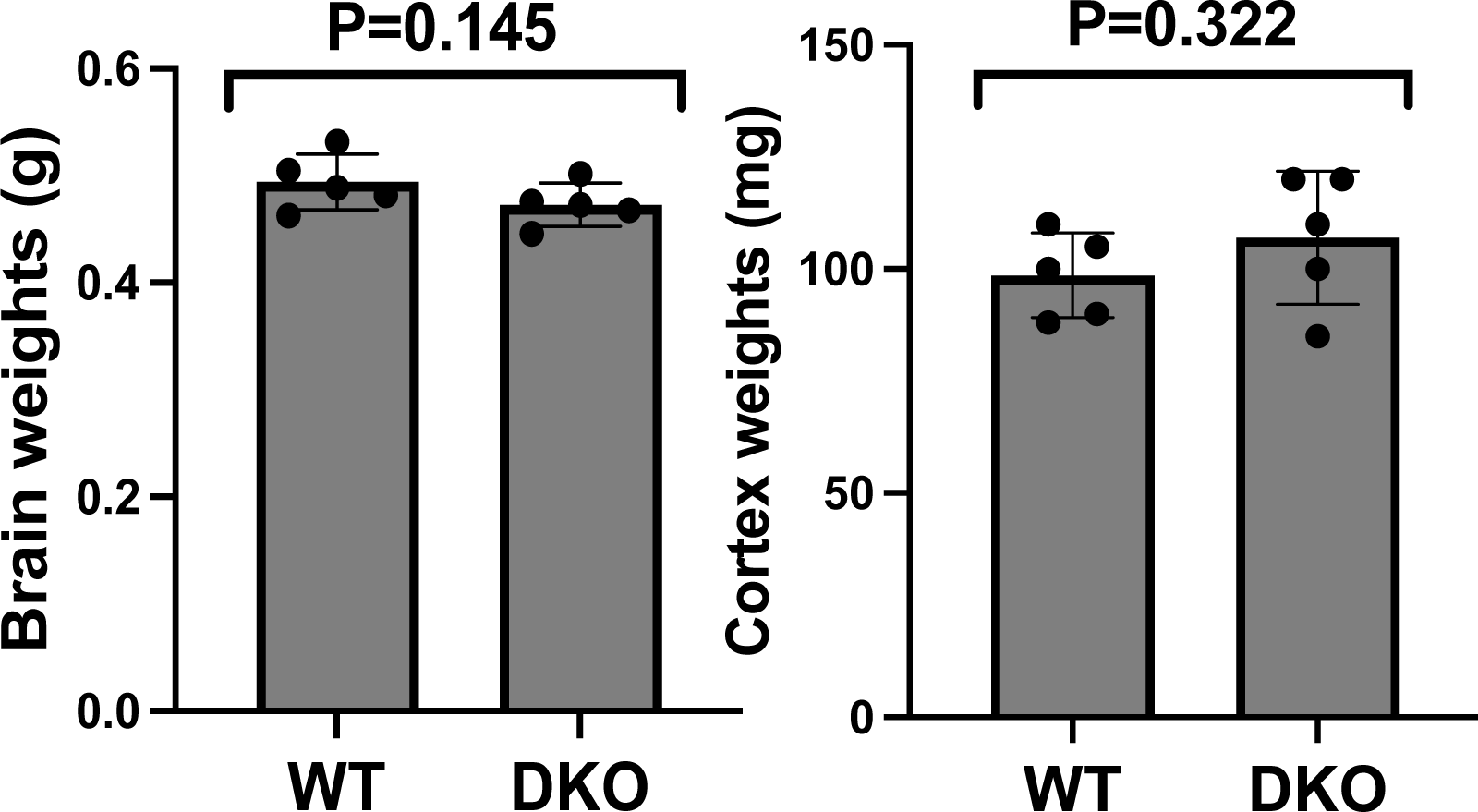
Compared to brain and cortex weights for WT and DKO mice. The figure on the left shows the weight of the mouse brain. Light figure shows the weight of the mouse cerebral cortex (20w).

**Fig. S14.**
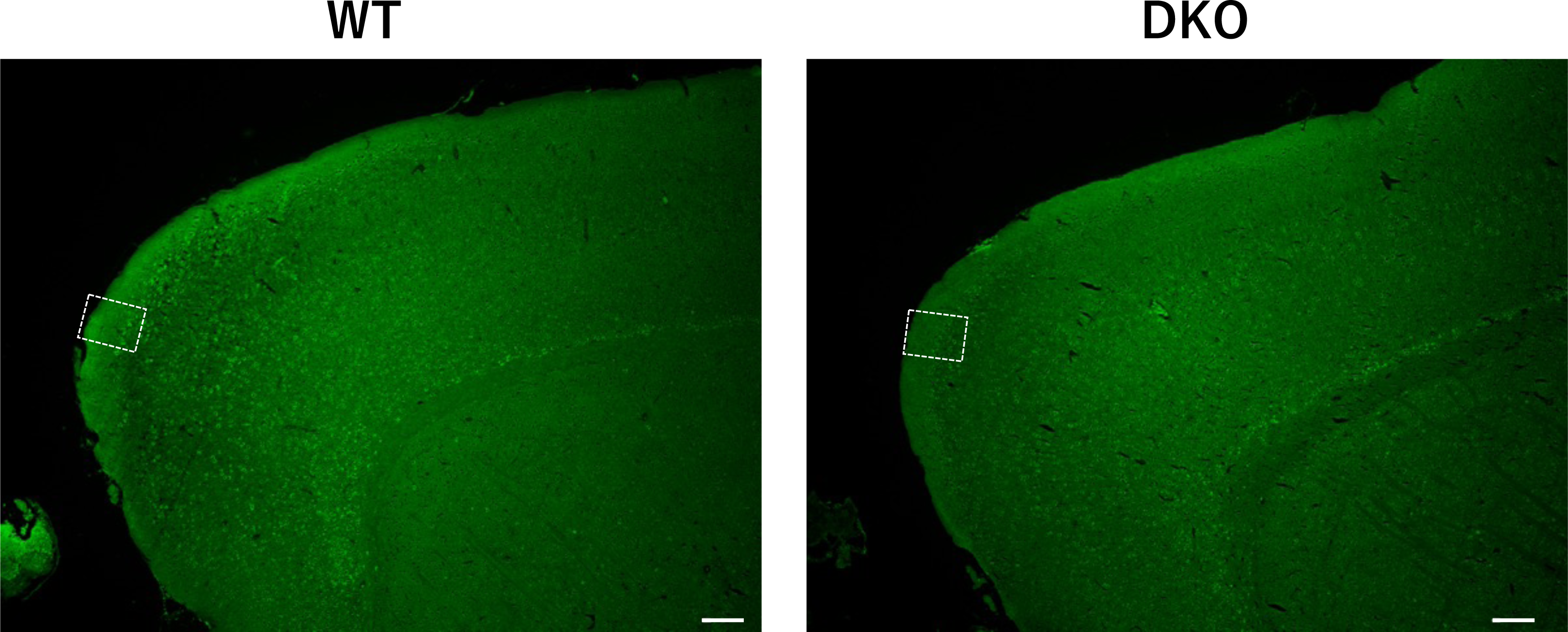
Fluorescence immunostaining images of the FrA region in WT and DKO mice. The areas surrounded by white squares are enlarged in Fig. 5A. Green: MAP2. Scale bar: 200 μm.

**Fig. S15.**
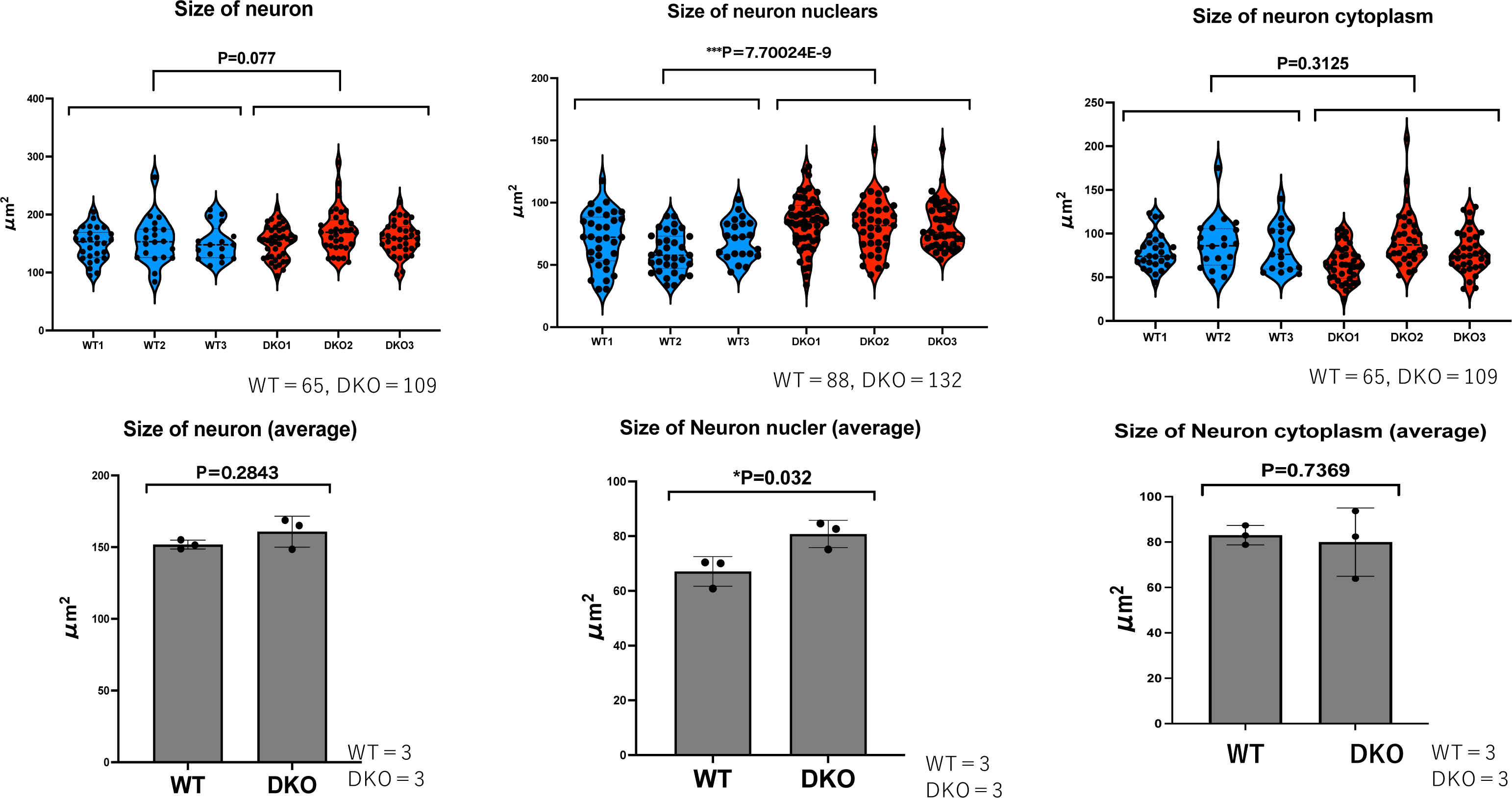
The sizes and average sizes of neurons, neuronal nuclei and neuronal cytoplasm in the FrA region. Top: Each of three individuals of the WT (blue) and DKO (red) data in Fig. 3S were presented. Bottom: Their averages were presented statistically analyzed with Student’s t-test. *P < 0.05, **P < 0.01, ***P< 0.001.

**Fig. S16.**
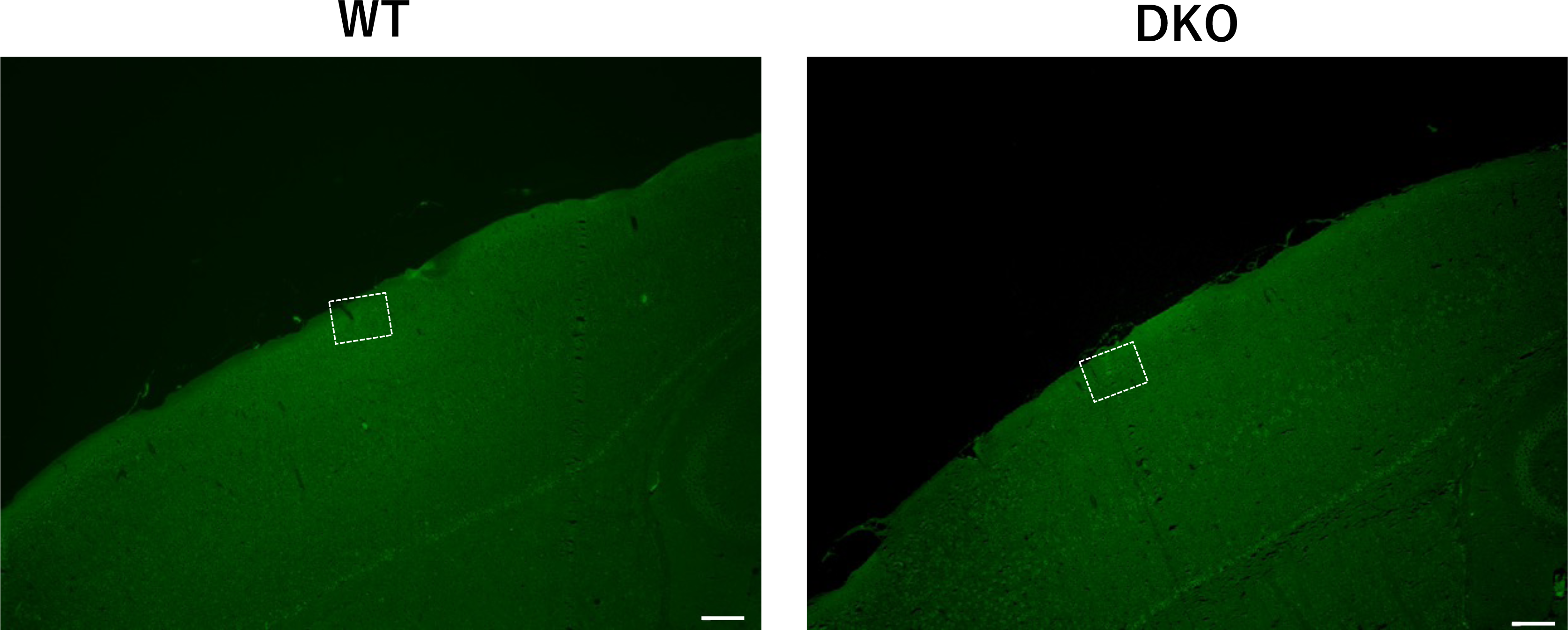
Fluorescence immunostaining images of the S1Sh region in WT and DKO mice (1) Sagittal sections, each at 20 w are shown. The areas surrounded by white dashed squares are enlarged in Fig. S17. Green: MAP2. Scale bar: 200 μm.

**Fig. S17.**
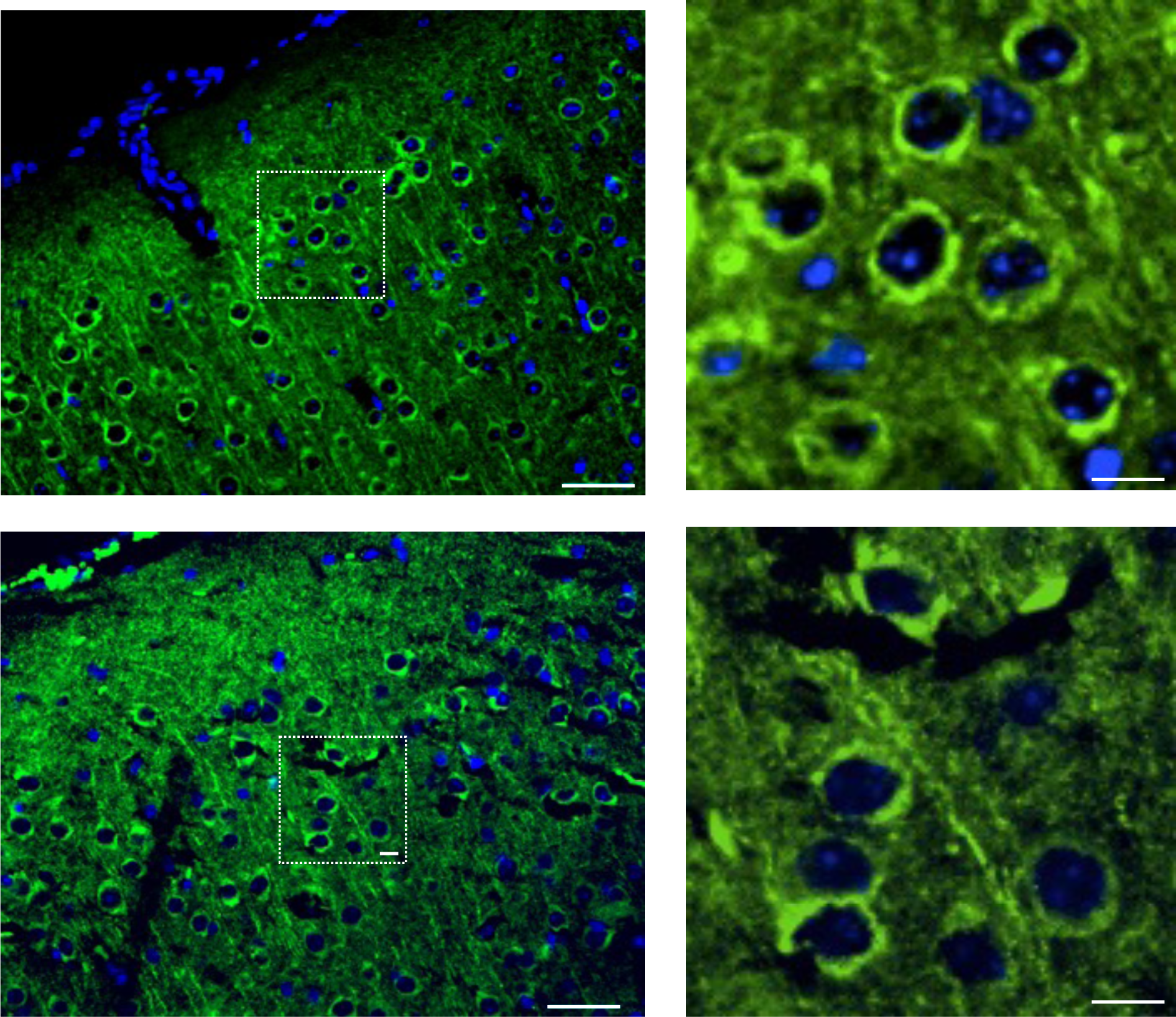
Fluorescence immunostaining images of the S1Sh region in WT and DKO mice (2) Left: Enlarged images of the dashed box areas in the Fig. S16. Scale bars: 40 μm. Right: Magnified views of the dashed box areas in the left pictures. Scale bars: 10 μm. Green: MAP2, red: DAPI.

**Fig. S18.**
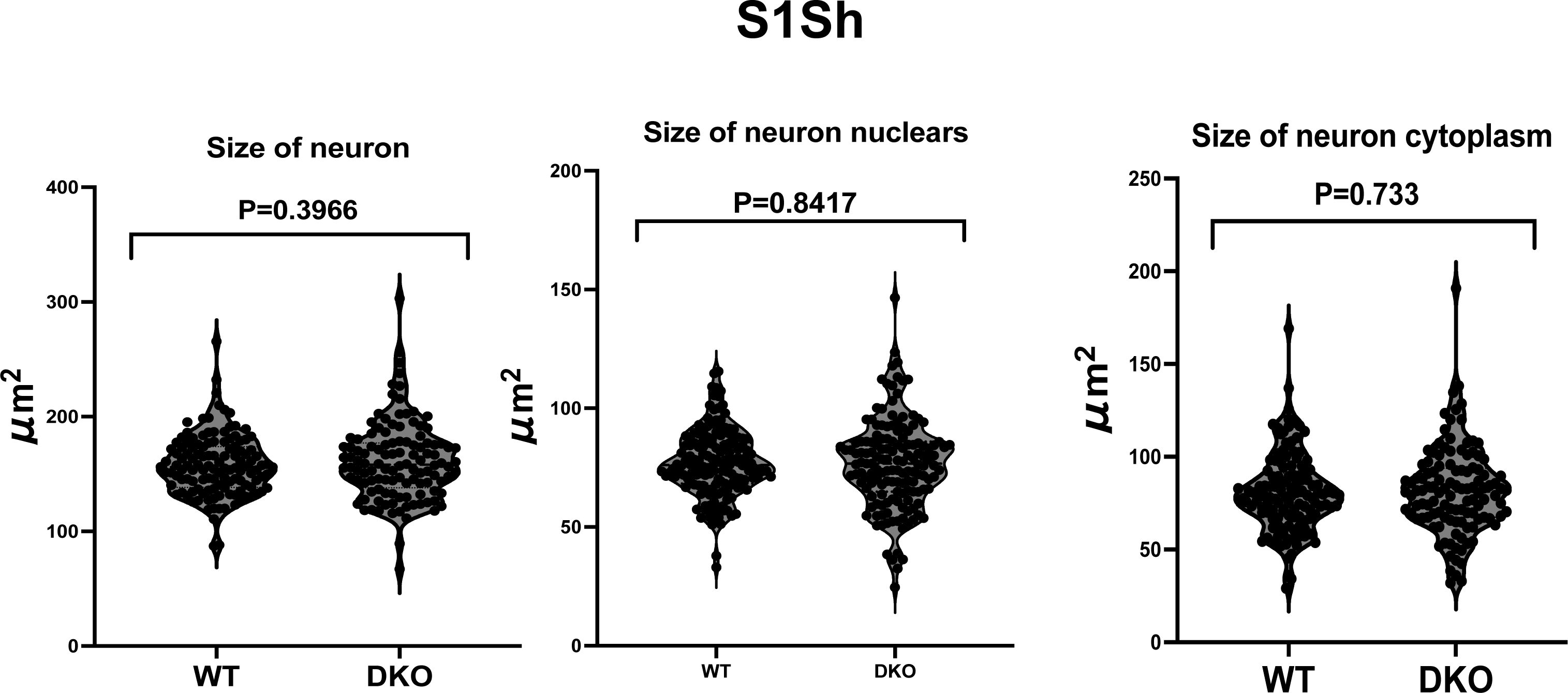
The sizes and average sizes of the neurons, neuronal nuclei and neuronal cytoplasm in the S1Sh region (1) The sizes of the neuronal cells and their nuclei in Fig.S17 were measured by a blind observer using ImageJ. Three individuals from each of the WT (blue) and KO (red) mice were examined for the measurement of a total 125 and 116 neurons, 163 and 149 neuronal nuclei and 125 and 116 neuronal cytoplasm samples, respectively (see also Fig. S19 top). The averages are presented. All measurements were statistically analyzed with Student’s t-test and the results are displayed as Violin Plots.

**Fig. S19.**
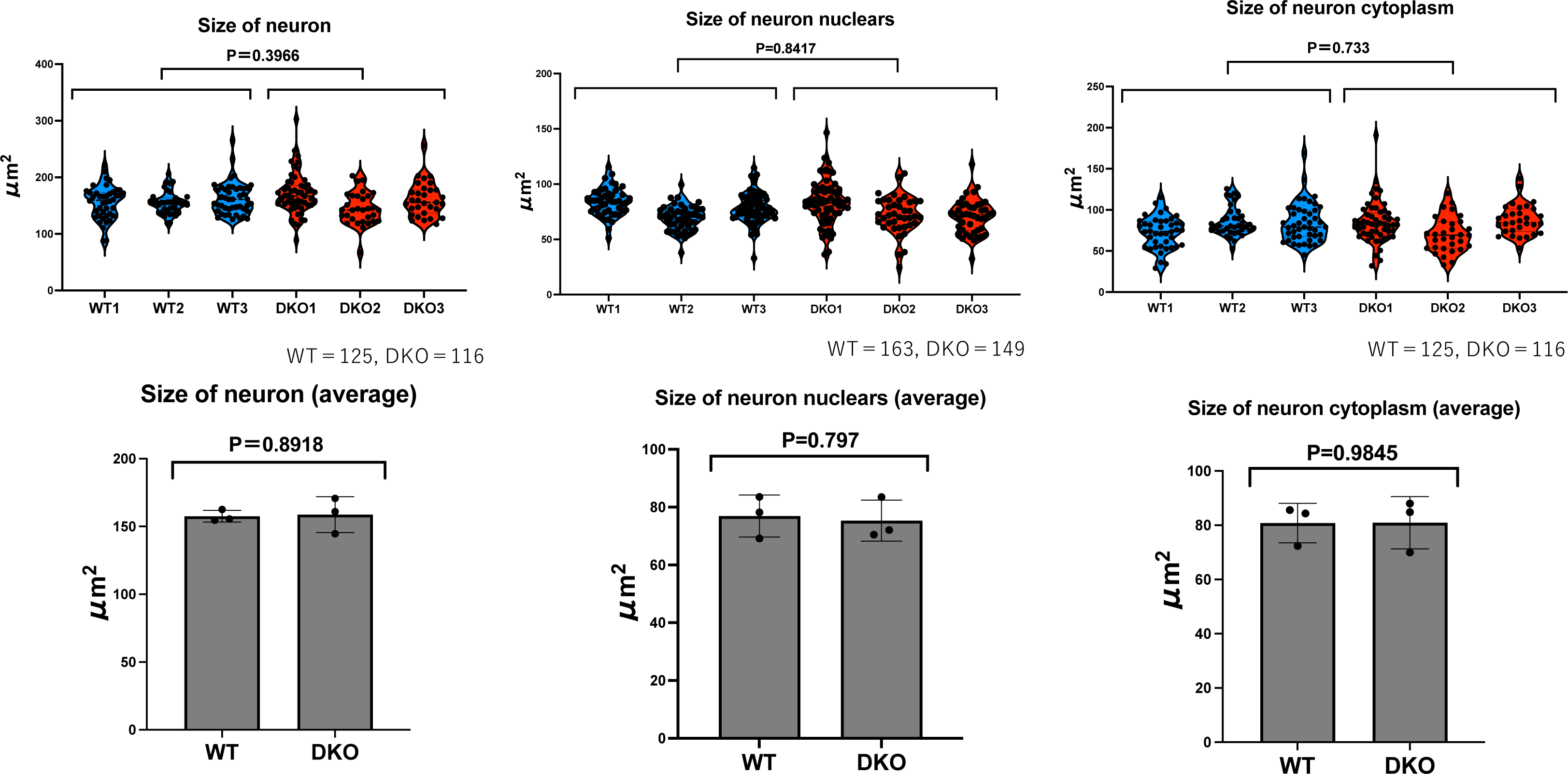
The sizes and average sizes of the neurons, neuronal nuclei and neuronal cytoplasm in the S1Sh region (2) Top: Each of three individuals of the WT (blue) and DKO (red) data in Fig. S18 were presented. Bottom: Their averages were presented statistically analyzed with Student’s t-test.

**Fig. S20.**
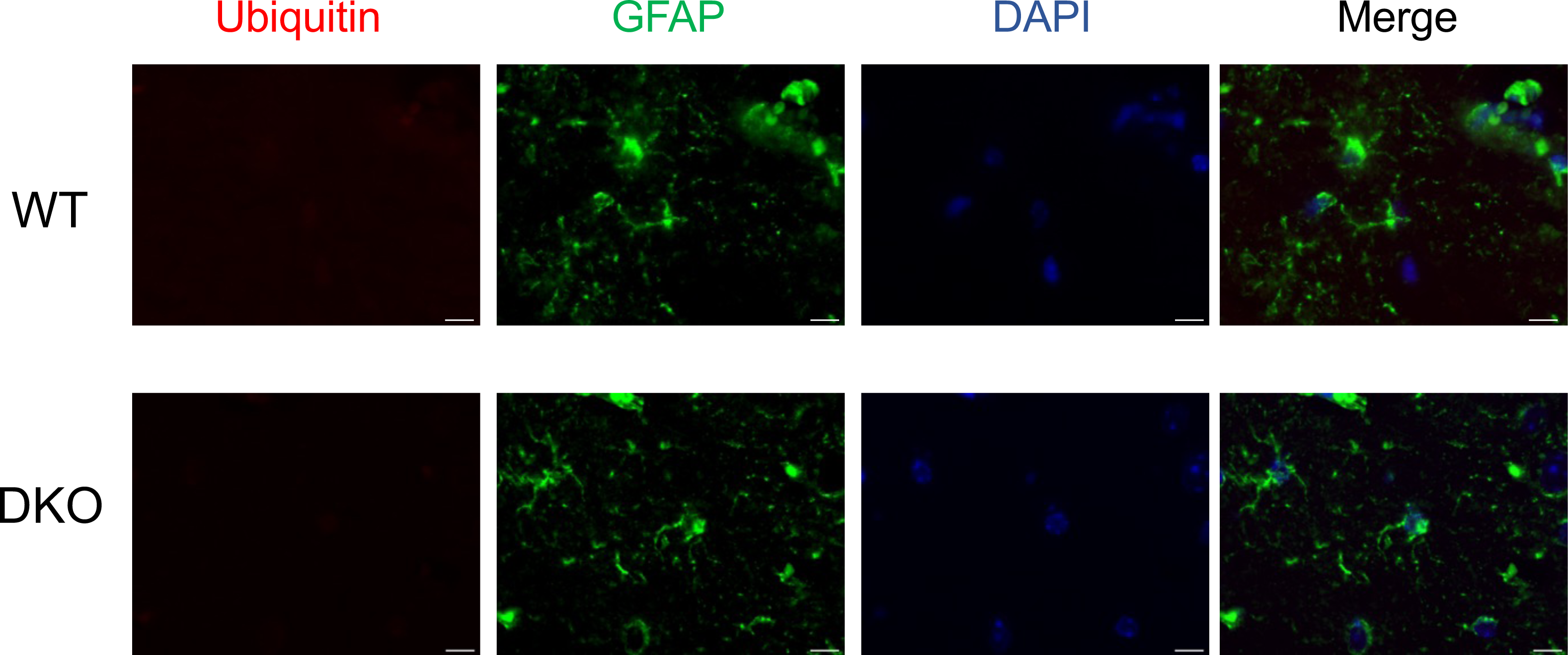
Co-immunofluorescence staining images of Ubiquitin and GFAP in the FrA region in WT and DKO. Sagittal sections, each at 20 w. Scale bars: 10 μm. Blue: DAPI, Green: MAP2, Red: Ubiquitin

**Fig. S21.**
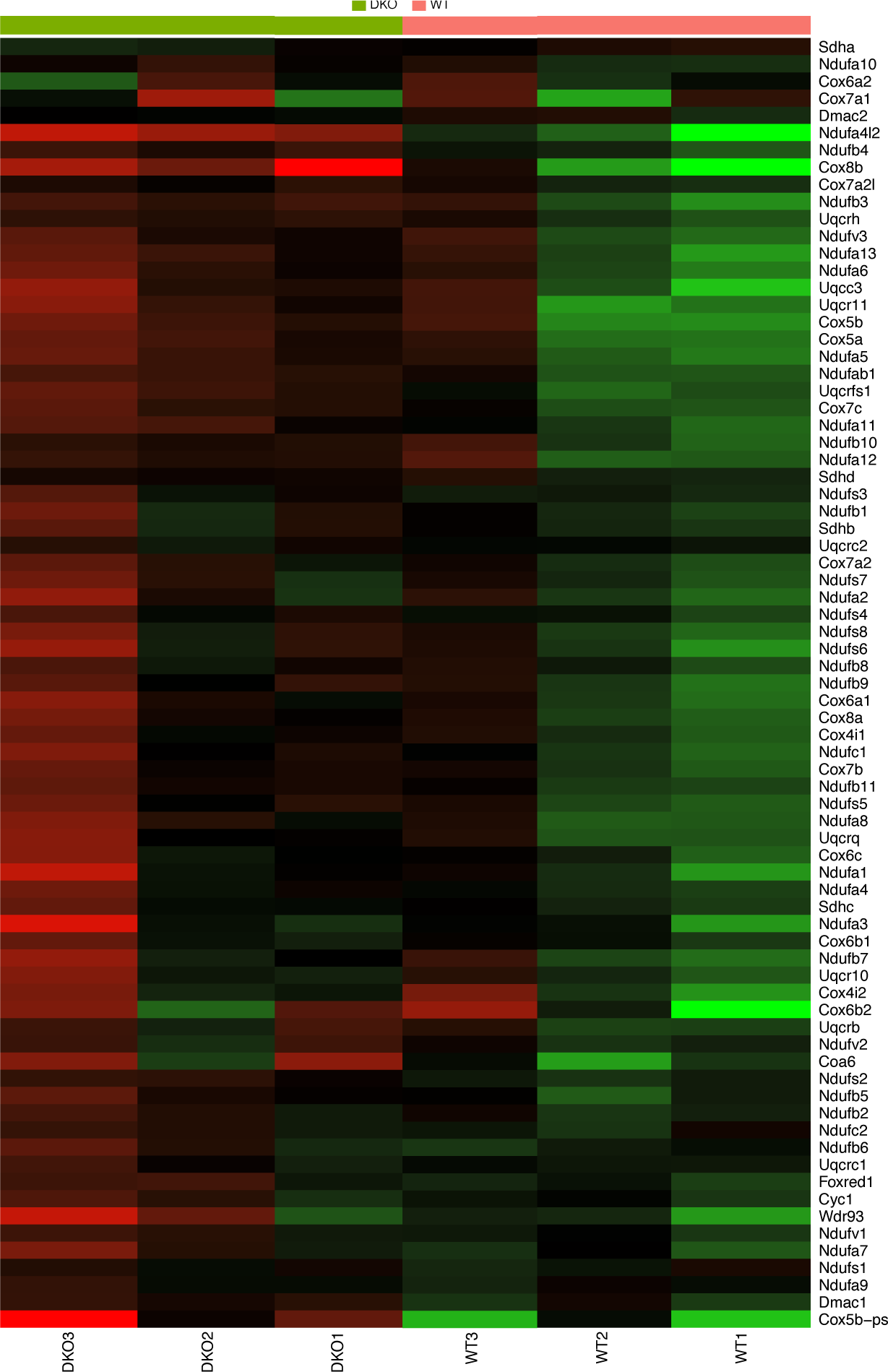
Heat map of of genes categorized Respiratory chain complex. The heat map shows the expression level (normalized Fold Change (FC) of each individual for the gene expression level by categorized “Respiratory chain complex”.

**Fig. S22.**
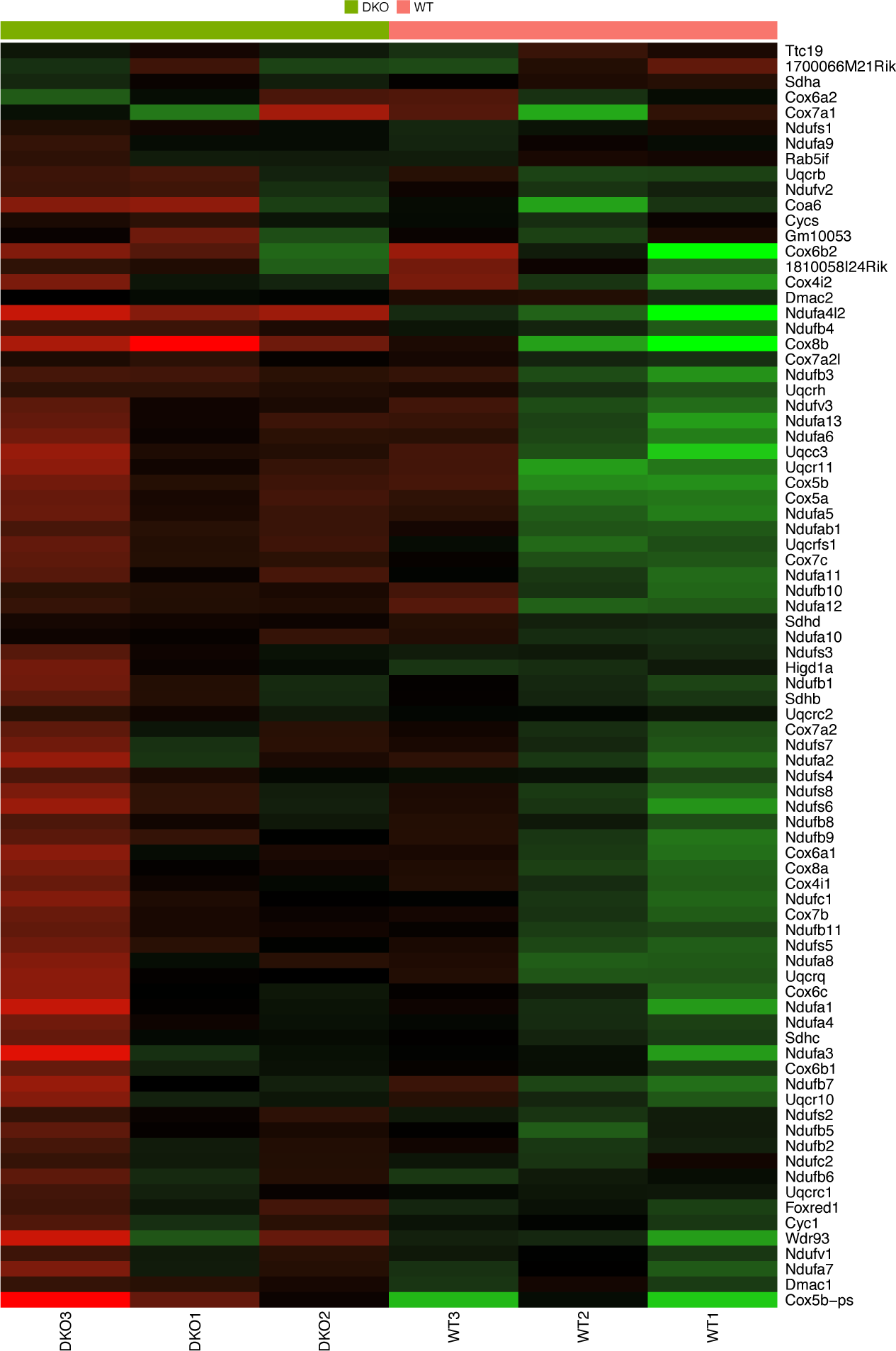
Heat Map of genes categriozed Respirasome. The heat map shows the expression level (normalized Fold Change (FC) of each individual for the gene expression level by categorized “Respirasome”.

**Fig. S23.**
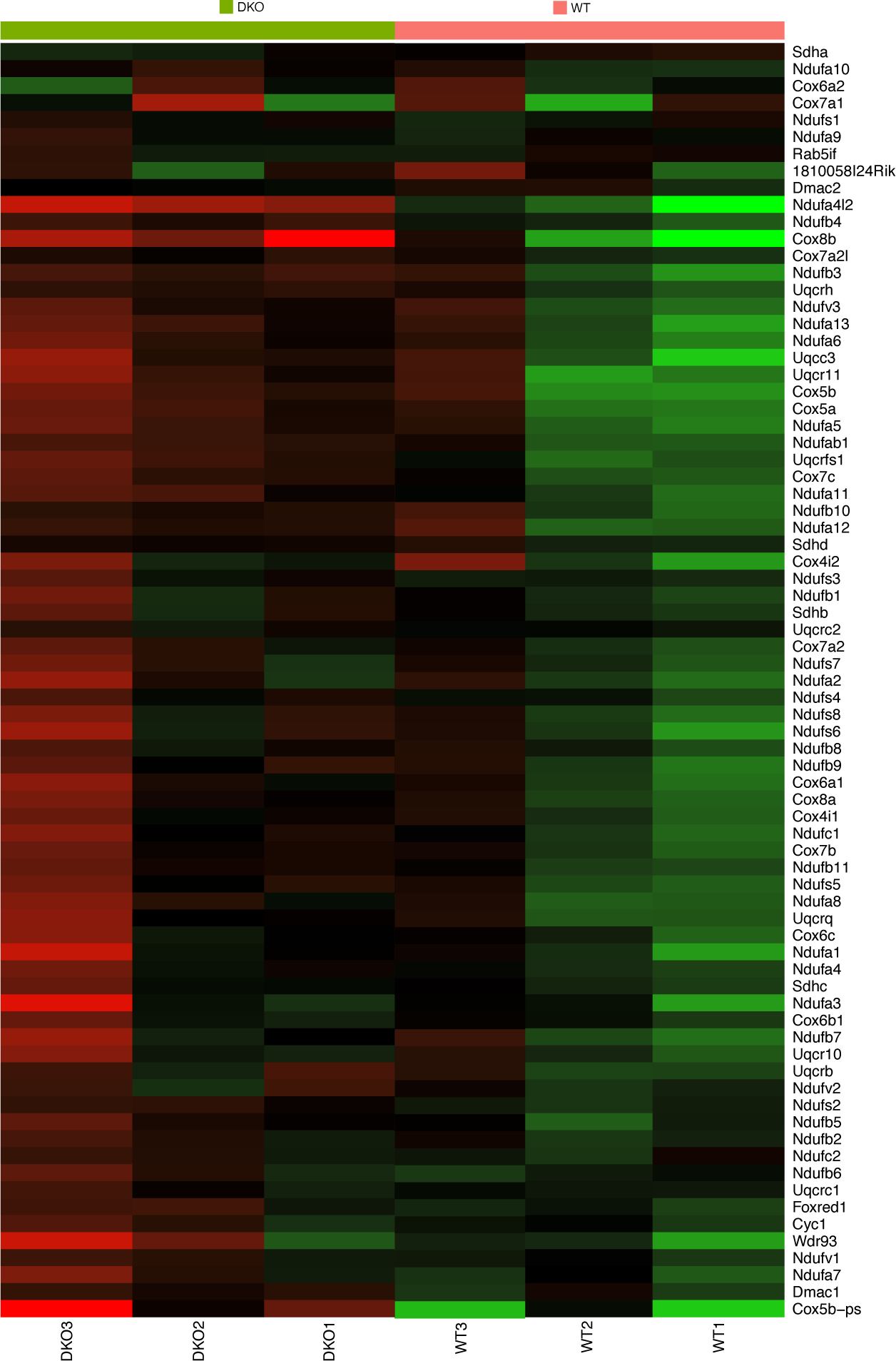
Heat map of of genes categorized Mitochondria. The heat map shows the expression level (normalized Fold Change (FC) of each individual for the gene expression level by categorized “mitochondria”.

**Table S1.** High conservation of *RTL8* genes in eutherian species. *RTL8* genes are mostly comprised of 2-4 homologs as shown in the figures in parentheses in each species, except humans and mice. The dN/dS values of *RTL8* genes in several eutherian species are shown. These values are especially low in the cases shown in red (dN/dS < 0.1) and pink (0.1 ≤ dN/dS < 0.2), indicating that *RTL8* genes are highly conserved in eutherians. The other sequences are also below 0.5, as shown in orange (0.2 ≤ dN/dS < 0.3) and yellow (0.3 ≤ dN/dS < 0.5). The asterisks indicate that the DNA sequences and aa sequences are completely the same between the two genes. It should be noted that in most cases there is no orthogous relationship among *RTL8A*, *8B* and *8C* in eutherian species. For example, human *RTL8A*, *8B* and *8C* and mouse *Rtl8a*, *8b* and *8c* are not orthologs, respectively (see also Fig. S1).

|  | 1 | 2 | 3 | 4 | 5 | 6 | 7 | 8 | 9 | 10 | 11 | 12 | 13 | 14 | 15 | 16 | 17 | 18 | 19 | 20 | 21 | 22 | 23 | 24 | 25 |
| --- | --- | --- | --- | --- | --- | --- | --- | --- | --- | --- | --- | --- | --- | --- | --- | --- | --- | --- | --- | --- | --- | --- | --- | --- | --- |
| 1. Human_RTL8B |  |  |  |  |  |  |  |  |  |  |  |  |  |  |  |  |  |  |  |  |  |  |  |  |  |
| 2. Human_RTL8C | 0.1088 |  |  |  |  |  |  |  |  |  |  |  |  |  |  |  |  |  |  |  |  |  |  |  |  |
| 3. Human_RTL8A | 0.0715 | 0.1499 |  |  |  |  |  |  |  |  |  |  |  |  |  |  |  |  |  |  |  |  |  |  |  |
| 4. Chimp_RTL8s(1) | 0.1075 | 0.1099 | 0.122 |  |  |  |  |  |  |  |  |  |  |  |  |  |  |  |  |  |  |  |  |  |  |
| 5. Chimp_RTL8s(2) | 0.1188 | 0.001 | 0.1699 | 0.1224 |  |  |  |  |  |  |  |  |  |  |  |  |  |  |  |  |  |  |  |  |  |
| 6. Chimp_RTL8s(3) | 0.0822 | 0.1512 | 0.0995 | 0.1197 | 0.1746 |  |  |  |  |  |  |  |  |  |  |  |  |  |  |  |  |  |  |  |  |
| 7. Marmoset_RTL8s(1) | 0.1754 | 0.2162 | 0.1442 | 0.2186 | 0.267 | 0.1442 |  |  |  |  |  |  |  |  |  |  |  |  |  |  |  |  |  |  |  |
| 8. Marmoset_RTL8s(2) | 0.12 | 0.001 | 0.1288 | 0.1261 | 0.001 | 0.176 | 0.1753 |  |  |  |  |  |  |  |  |  |  |  |  |  |  |  |  |  |  |
| 9. Marmoset_RTL8s(3) | 0.2648 | 0.1859 | 0.2112 | 0.2623 | 0.2006 | 0.2217 | 0.2008 | 0.2021 |  |  |  |  |  |  |  |  |  |  |  |  |  |  |  |  |  |
| 10. Mouse_Rtl8c | 0.1008 | 0.0515 | 0.0799 | 0.0864 | 0.0565 | 0.0836 | 0.076 | 0.07 | 0.0829 |  |  |  |  |  |  |  |  |  |  |  |  |  |  |  |  |
| 11. Mouse_Rtl8a | 0.1259 | 0.0545 | 0.1243 | 0.1144 | 0.0602 | 0.1151 | 0.1148 | 0.0664 | 0.1247 | 0.3271 |  |  |  |  |  |  |  |  |  |  |  |  |  |  |  |
| 12. Mouse_Rtl8b | 0.1259 | 0.0545 | 0.1243 | 0.1144 | 0.0602 | 0.1151 | 0.1148 | 0.0664 | 0.1247 | 0.3271 | * |  |  |  |  |  |  |  |  |  |  |  |  |  |  |
| 13. Cow_RTL8s(1) | 0.1137 | 0.0632 | 0.0979 | 0.0919 | 0.0621 | 0.094 | 0.0867 | 0.0832 | 0.1077 | 0.0698 | 0.040 | 0.040 |  |  |  |  |  |  |  |  |  |  |  |  |  |
| 14. Cow_RTL8s(2) | 0.1313 | 0.0614 | 0.1134 | 0.1051 | 0.0602 | 0.1082 | 0.1008 | 0.0710 | 0.1224 | 0.0665 | 0.0551 | 0.0551 | 0.001 |  |  |  |  |  |  |  |  |  |  |  |  |
| 15. Cow_RTL8s(3) | 0.1313 | 0.0614 | 0.1134 | 0.1051 | 0.0602 | 0.1082 | 0.1008 | 0.0710 | 0.1224 | 0.0665 | 0.0551 | 0.0551 | 0.001 | * |  |  |  |  |  |  |  |  |  |  |  |
| 16. Rabbit_RTL8s(1) | 0.0635 | 0.0817 | 0.0672 | 0.0624 | 0.0877 | 0.0674 | 0.1059 | 0.0684 | 0.1240 | 0.046 | 0.0620 | 0.0620 | 0.0752 | 0.0858 | 0.0858 |  |  |  |  |  |  |  |  |  |  |
| 17. Rabbit_RTL8s(2) | 0.1307 | 0.1224 | 0.1329 | 0.0624 | 0.1278 | 0.1365 | 0.1716 | 0.1033 | 0.1288 | 0.0685 | 0.0804 | 0.0804 | 0.1069 | 0.0936 | 0.0936 | 0.2344 |  |  |  |  |  |  |  |  |  |
| 18. Rabbit_RTL8s(3) | 0.0635 | 0.0817 | 0.0672 | 0.0624 | 0.0877 | 0.0674 | 0.1059 | 0.0684 | 0.1240 | 0.046 | 0.0620 | 0.0620 | 0.0752 | 0.0858 | 0.0858 | * | 0.2344 |  |  |  |  |  |  |  |  |
| 19. Rabbit_RTL8s(4) | 0.0982 | 0.1172 |  | 0.0953 | 0.1233 | 0.0909 | 0.1288 | 0.0956 | 0.1512 | 0.0045 | 0.0029 | 0.0029 | 0.0841 | 0.0940 | 0.0940 | 0.1607 | 0.1755 | 0.1607 |  |  |  |  |  |  |  |
| 20. Dog_RTL8s(1) | 0.1801 | 0.0794 | 0.1396 | 0.1408 | 0.0781 | 0.1470 | 0.1628 | 0.0931 | 0.1775 | 0.0732 | 0.0717 | 0.0717 | 0.0682 | 0.0901 | 0.0901 | 0.0845 | 0.0782 | 0.0845 | 0.0806 |  |  |  |  |  |  |
| 21. Dog_RTL8s(2) | 0.1801 | 0.0794 | 0.1396 | 0.1408 | 0.0781 | 0.1470 | 0.1628 | 0.0931 | 0.1775 | 0.0732 | 0.0717 | 0.0717 | 0.0682 | 0.0901 | 0.0901 | 0.0845 | 0.0782 | 0.0845 | 0.0806 | * |  |  |  |  |  |
| 22. Dog_RTL8s(3) | 0.1801 | 0.0794 | 0.1396 | 0.1408 | 0.0781 | 0.1470 | 0.1628 | 0.0931 | 0.1775 | 0.0732 | 0.0717 | 0.0717 | 0.0682 | 0.0901 | 0.0901 | 0.0845 | 0.0782 | 0.0845 | 0.0806 | * | * |  |  |  |  |
| 23. Dog_RTL8s(4) | 0.1801 | 0.0794 | 0.1396 | 0.1408 | 0.0781 | 0.1470 | 0.1628 | 0.0931 | 0.1775 | 0.0732 | 0.0717 | 0.0717 | 0.0682 | 0.0901 | 0.0901 | 0.0845 | 0.0782 | 0.0845 | 0.0806 | * | * | * |  |  |  |
| 24. Elephant_RTL8s(1) | 0.0959 | 0.1147 | 0.1209 | 0.1021 | 0.1213 | 0.1007 | 0.1096 | 0.1144 | 0.1629 | 0.0122 | 0.0706 | 0.0706 | 0.0801 | 0.0907 | 0.0907 | 0.1015 | 0.1421 | 0.1015 | 0.0774 | 0.1184 | 0.1184 | 0.1184 | 0.1184 |  |  |
| 25. Elephant_RTL8s(2) | 0.0993 | 0.1317 | 0.1419 | 0.1059 | 0.1394 | 0.1195 | 0.1307 | 0.1310 | 0.1792 | 0.037 | 0.0657 | 0.0657 | 0.089 | 0.1014 | 0.1014 | 0.1105 | 0.1377 | 0.1105 | 0.0844 | 0.1362 | 0.1362 | 0.1362 | 0.1362 | 0.2159 |  |
| 26. Elephant_RTL8s(3) | 0.1654 | 0.2096 | 0.2112 | 0.1750 | 0.2200 | 0.1828 | 0.1977 | 0.1934 | 0.2266 | 0.0323 | 0.0889 | 0.0889 | 0.1433 | 0.1616 | 0.1616 | 0.1441 | 0.2004 | 0.1441 | 0.0909 | 0.2142 | 0.2142 | 0.2142 | 0.2142 | 0.4607 | 0.2942 |

## References

Ashburner M, Ball CA, Blake JA, Botstein D, Butler H, Cherry JM, Davis AP, Dolinski K, Dwight SS, Eppig JT et al (2000) Gene ontology: tool for the unification of biology. Nat Genet 25: 25–29

Atasoy D, Betley JN, Su HH, Sternson SM (2012) Deconstruction of a neural circuit for hunger. Nature 488: 172–177

Barki M, Xue H (2022) GABRB2, a key player in neuropsychiatric disorders and beyond. Gene 809: 146021

Baumann SW, Baur R, Sigel E (2002) Forced subunit assembly in alpha1beta2gamma2 GABAA receptors. Insight into the absolute arrangement. J Biol Chem 277: 46020– 46025

Bolger AM, Lohse M, Usadel B (2014) Trimmomatic: a flexible trimmer for Illumina sequence data. Bioinformatics 30: 2114–2120

Brandt J, Veith AM, Volff JN (2005) Transposable elements as a source of genetic innovation: expression and evolution of a family of retrotransposon-derived neogenes in mammals. Gene 345: 101–111

Broschard MB, Kim J, Love BC, Wasserman EA, Freeman JH (2021) Prelimbic cortex maintains attention to category-relevant information and flexibly updates category representations. Neurobiol Learn Mem 185: 107524

Chamberlain SJ, Lalande M (2010) Angelman syndrome, a genomic imprinting disorder of the brain. J Neurosci. 30: 9958–9963

Charlier C, Segers K, Wagenaar D, Karim L, Berghmans S, Jaillon O, Shay T, Weissenbach J, Cockett N, Gyapay G et al (2001) Human–ovine comparative sequencing of a 250-kb imprinted domain encompassing the callipyge (*clpg*) locus and identification of six imprinted transcripts: *DLK1*, *DAT, GTL2*, PEG11, antiPEG11, and MEG8. Genome Res 11: 850–862

Chen J, Tsang SY, Zhao CY, Pun FW, Yu Z, Mei L, Lo WS, Fang S, Liu H, Stöber G, Xue H (2009) GABRB2 in schizophrenia and bipolar disorder: disease association, gene expression and clinical correlations. Biochem Soc Trans 37: 1415–1418

Chou M-Y, Hu M-C, Chen P-Y, Hsu C-L, Lin T-Y, Tan M-J, Chih Y-L, Meng F-K, Pei H-H, Vin C-W et al (2022) RTL1/PEG11 imprinted in human and mouse brain mediates anxiety-like and social behaviors and regulates neuronal excitability in the locus coeruleus. Hum Mol Genet 31: 3161–3180

Clark DA, Arranz MJ, Mata I, Lopéz-Ilundain J, Pérez-Nievas F, Kerwin RW (2005) Polymorphisms in the promoter region of the alpha1A-adrenoceptor gene are associated with schizophrenia/schizoaffective disorder in a Spanish isolate population. Biol Psychiatry 58:435–439

Deng H-X, Chen W, Hong S-T, Boycott KM, Gorrie GH, Siddique N, Yang Y, Fecto F, Shi Y, Zhai H et al (2011) Mutations in UBQLN2 cause dominant X-linked juvenile and adult-onset ALS and ALS/dementia. Nature 477: 211–215

Dobin A, Davis CA, Schlesinger F, Drenkow J, Zaleski C, Jha S, Batut P, Chaisson M, Gingeras TR (2013) STAR: ultrafast universal RNA-seq aligner. Bioinformatics 29: 15–21

Elia J, Capasso M, Zaheer Z, Lantieri F, Ambrosini P, Berrettini W, Devoto M, Hakonarson H (2009) Candidate gene analysis in an on-going genome-wide association study of attention-deficit hyperactivity disorder: suggestive association signals in ADRA1A. Psychiatr Genet 19:134–141

Florez L, Anderson M, Lacassie Y (2003) De novo paracentric inversion (X)(q26q28) with features mimicking Prader-Willi syndrome. Am J Med Genet A 121A: 60–64

Ge SX, Son EW, Yao R (2018) iDEP: an integrated web application for differential expression and pathway analysis of RNA-Seq data. BMC Bioinformatics 19: 534

Germain ND, Levine ES, Chamberlain SJ (2020) IPSC Models of Chromosome 15Q Imprinting Disorders: From Disease Modeling to Therapeutic Strategies. Adv Neurobiol 25: 55–77.

Hare BD, Duman RS (2020) Prefrontal cortex circuits in depression and anxiety: contribution of discrete neuronal populations and target regions. Mol Psychiatry 25: 2742–2758

Hare TA, Camerer CF, Rangel A (2009) Self-control in decision-making involves modulation of the vmPFC valuation system. Science 324: 646–648

Henriques WS, Young JM, Nemudryi A, Nemudraia A, Wiedenheft B, Malik HS (2024) The Diverse Evolutionary Histories of Domesticated Metaviral Capsid Genes in Mammals. Mol Biol Evol 41: msae061

Imakawa K, Kusama K, Kaneko-Ishino T, Nakagawa S, Kitao K, Miyazawa T, Ishino F (2022) Endogenous Retroviruses and Placental Evolution, Development, and Diversity. Cells 11 2458

Ioannides Y, Lokulo-Sodipe K, Mackay DJG, Davies JH, Temple IK (2014) Temple syndrome: improving the recognition of an underdiagnosed chromosome 14 imprinting disorder: an analysis of 51 published cases. J Med Genet 51: 495–501

Irie M, Itoh J, Matsuzawa A, Ikawa M, Kiyonari H, Kihara M, Suzuki T, Hiraoka Y, Ishino F, Kaneko-Ishino T (2022) Retrovirus-derived *RTL5* and *RTL6* genes are novel constituents of the innate immune system in the eutherian brain. Development 149: dev200976

Irie M, Yoshikawa M, Ono R, Iwafune H, Furuse T, Yamada I, Wakana S, Yamashita Y, Abe T, Ishino F et al (2015). Cognitive Function Related to the *Sirh11/Zcchc16* Gene Acquired from an LTR Retrotransposon in Eutherians. PLoS Genetics 11: e1005521

Ishino F, Itoh J, Irie M, Matsuzawa A, Naruse M, Suzuki T, Hiraoka Y, Kaneko-Ishino T (2023) Retrovirus-derived RTL9 plays an important role in innate antifungal immunity in the eutherian brain. Int J Mol Sci 24: 14884

Jiang S, Wen N, Li Z, Dube U, Del Aguila J, Budde J, Martinez R, Hsu S, Fernandez MV, Cairns NJ, et al (2018) Integrative system biology analyses of CRISPR-edited iPSC-derived neurons and human brains reveal deficiencies of presynaptic signaling in FTLD and PSP. Transl Psychiatry, 8: 265

Kagami M, Kurosawa K, Miyazaki O, Ishino F, Matsuoka K, Ogata T (2015) Comprehensive clinical studies in 34 patients with molecularly defined UPD (14)pat and related conditions (Kagami Ogata syndrome). Eur J Hum Genet 23: 1488–1498

Kagami M, Sekita Y, Nishimura G, Irie M, Kato F, Okada M, Yamamori S, Kishimoto H, Nakayama M, Tanaka Y et al (2008) Deletions and epimutations affecting the human 14q32.2 imprinted region in individuals with paternal and maternal upd(14)-like phenotypes. Nat Genet 40: 237–242

Kalderon D, Roberts BL, Richardson WD, Smith AE (1984) A short amino acid sequence able to specify nuclear location. Cell 39: 499–509

Kaneko-Ishino T, Ishino F (2012) The role of genes domesticated from LTR retrotransposons and retroviruses in mammals. Front Microbiol 3: 262

Kaneko-Ishino T, Ishino F (2015) Mammalian-specific genomic functions: Newly acquired traits generated by genomic imprinting and LTR retrotransposon-derived genes in mammals. Proc Jpn Acad Ser B Phys Biol Sci 91: 511–538

Kaneko-Ishino T, Ishino F (2023) Retrovirus-derived RTL/SIRH: their diverse role in the current eutherian developmental system and contribution to eutherian evolution. Biomolecules 13: 1436

Kim A, Terzian C, Santamaria P, Pelisson A, Purd’homme N, Bucheton A (1994) Retroviruses in invertebrates: The gypsy retrotransposon is apparently an infectious retrovirus of Drosophila melanogaster. Proc Natl Acad Sci USA 91: 1285–1289

Kitazawa M, Hayashi S, Imamura M, Takeda S, Oishi Y, Kaneko-Ishino T, & Ishino F. (2020). Deficiency and overexpression of *Rtl1* in the mouse cause distinct muscle abnormalities related to Temple and Kagami-Ogata syndromes. Development 147: dev185918

Kitazawa M, Sutani A, Kaneko-Ishino T, Ishino F (2021) The role of eutherian-specific RTL1 in the nervous system and its implications for the Kagami-Ogata and Temple syndromes. Genes Cells 26: 165–179

Kroiss M, Leyerer M, Gorboulev V, Kühlkamp T, Kipp H, Koepsell H (2006) Transporter regulator RS1 (RSC1A1) coats the trans-Golgi network and migrates into the nucleus. Am J Physiol Renal Physiol 291: F1201–1212

Lanfray D, Richard D (2017) Emerging Signaling Pathway in Arcuate Feeding-Related Neurons: Role of the Acbd7. Front Neurosci 11: 328

Lim ET, Raychaudhuri S, Sanders SJ, Stevens C, Sabo A, MacArthur DG, Neale BM, Kirby A, Ruderfer DM, Fromer M et al (2013) Rare complete knockouts in humans: population distribution and significant role in autism spectrum disorders. Neuron 77: 235–242

Lo W-S, Harano M, Gawlik M, Yu Z, Chen J, Pun F W, Tong K-L, Zhao C, Ng S-K, Tsang S-Y et al (2007) GABRB2 association with schizophrenia: commonalities and differences between ethnic groups and clinical subtypes. Biol Psychiatry 61: 653– 660

Lo W-S, Lau C-F, Xuan Z, Chan C-F, Feng G-Y, He L, Cao Z-C, Liu H, Luan Q-M, Xue H (2004) Association of SNPs and haplotypes in GABAA receptor beta2 gene with schizophrenia. Mol Psychiatry 9: 603–608

Lucignani G, Panzacchi A, Bosio L, Moresco RM, Ravasi L, Coppa I, Chiumello G, Frey K, Koeppe R, Fazio F (2004) GABA A receptor abnormalities in Prader-Willi syndrome assessed with positron emission tomography and [11C]flumazenil. Neuroimage 22: 22–28

Lyon MF (1961) Gene action in the X-chromosome of the mouse (Mus musculus L.). Nature 190: 372–373

Ma DQ, Whitehead PL, Menold MM, Martin ER, Ashley-Koch AE, Mei H, Ritchie MD, Delong GR, Abramson RK, Wright HH et al (2005) Identification of significant association and gene-gene interaction of GABA receptor subunit genes in autism. Am J Hum Genet 77: 377–388

McKernan RM, Whiting PJ (1996) Which GABAA-receptor subtypes really occur in the brain? Trends Neurosci 19: 139–143

Merkestein M, van Gestel MA, van der Zwaal EM, Brans MA, Luijendijk MC, van Rozen AJ, Hendriks J, Garner KM, Boender AJ, Pandit R et al (2014) GHS-R1a signaling in the DMH and VMH contributes to food anticipatory activity. Int J Obes (Lond) 38: 610–618

Mohan HM, Trzeciakiewicz H, Pithadia A, Crowley EV, Pacitto R, Safren N, Trotter B, Zhang C, Zhou X, Zhang Y et al (2022) RTL8 promotes nuclear localization of UBQLN2 to subnuclear compartments associated with protein quality control. Cell Mol Life Sci 79: 176

Nakayama D, Baraki Z, Onoue K, Ikegaya Y, Matsuki N, Nomura H (2015) Frontal association cortex is engaged in stimulus integration during associative learning. Curr Biol 25: 117–123

Naruse M, Ono R, Irie M, Nakamura K, Furuse T, Hino T, Oda K, Kashimura M, Yamada I, Wakana S et al (2014) *Sirh7/Ldoc1* knockout mice exhibit placental P4 overproduction and delayed parturition. Development 141: 4763–4771

Nicholls RD, Knepper JL (2001) Genome organization, function, and imprinting in Prader-Willi and Angelman syndromes. Annu Rev Genomics Hum Genet 2: 153–175

Ono R, Ishii M, Fujihara Y, Kitazawa M, Usami T, Kaneko-Ishino T, Kanno J, Ikawa M., Ishino F (2015) Double strand break repair by capture of retrotransposon sequences and reverse-transcribed spliced mRNA sequences in mouse zygotes. Sci Rep 5: 12281

Ono R, Kobayashi S, Wagatsuma H, Aisaka K, Kohda T, Kaneko-Ishino T, Ishino F (2001) A retrotransposon-derived gene, *PEG10*, is a novel imprinted gene located on human chromosome 7q21. Genomics 73: 232–237

Ono R, Nakamura K, Inoue K, Naruse M, Usami T, Wakisaka-Saito N, Hino T, Suzuki-Migishima R, Ogonuki N, Miki H et al (2006) Deletion of *Peg10*, an imprinted gene acquired from a retrotransposon, causes early embryonic lethality. Nat Genet 38: 101–106

Pandya NJ, Wang C, Costa V, Lopatta P, Meier S, Zampeta FI, Punt AM, Mientjes E, Grossen P, Distler T et al (2021) Secreted retrovirus-like GAG-domain-containing protein PEG10 is regulated by UBE3A and is involved in Angelman syndrome pathophysiology. Cell Rep Med 2: 100360

Petryshen TL, Middleton FA, Tahl AR, Rockwell GN, Purcell S, Aldinger KA, Kirby A, Morley CP, McGann L, Gentile KL et al (2005) Genetic investigation of chromosome 5q GABAA receptor subunit genes in schizophrenia. Mol Psychiatry 10: 1074–1088

Pizzagalli DA, Roberts AC (2022) Prefrontal cortex and depression. Neuropsychopharmacology, 47: 225–246

Satterstrom FK, Kosmicki JA, Wang J, Breen MS, De Rubeis S, An JY, Peng M, Collins R, Grove J, Klei L et al (2020) Large-scale exome sequencing study implicates both developmental and functional changes in the neurobiology of autism. Cell, 180: 568–584.e23

Seabrook LT, Borgland SL (2020) The orbitofrontal cortex, food intake and obesity. J Psychiatry Neurosci 45: 304–312

Sekita Y, Wagatsuma H, Nakamura K, Ono R, Kagami M, Wakisaka N, Hino T, Suzuki-Migishima R, Kohda T, Ogura A et al (2008) Role of retrotransposon-derived imprinted gene, *Rtl1*, in the feto-maternal interface of mouse placenta. Nat Genet 40: 243–248

Sente A, Desai R, Naydenova K, Malinauskas T, Jounaidi Y, Miehling J, Zhou X, Masiulis S, Hardwick SW, Chirgadze DY et al (2022) Differential assembly diversifies GABAA receptor structures and signalling. Nature 604: 190–194

Shiura H, Kitazawa M, Ishino F, Kaneko-Ishino T (2023) Functions of two retrovirus-derived acquired genes, *PEG10* and PEG11/RTL1, in mammalian development and their relation to human genomic imprinting diseases. Front Cell Dev Biol 11: 1273638

Shrestha P, Mousa A, Heintz N (2015) Layer 2/3 pyramidal cells in the medial prefrontal cortex moderate stress-induced depressive behaviors. eLife 4: e08752

Sigel E, Steinmann ME (2012) Structure, function, and modulation of GABA(A) receptors. J Biol Chem 287: 40224–40231

Song SU, Gerasimova T, Kurkulos M, Boeke JD, Corces VG (1994). An env-like protein encoded by a Drosophila retroelement: Evidence that gypsy is an infectious retrovirus. Genes Dev 8: 2046–2057

Suyama M, Torrents D, Bork P (2006) PAL2NAL: robust conversion of protein sequence alignments into the corresponding codon alignments. Nucleic Acids Res 34: W609–W612

Szczepanski SM, Knight RT (2014) Insights into human behavior from lesions to the prefrontal cortex. Neuron, 83: 1002–1018

Takagi N, Sasaki M (1975) Preferential inactivation of the paternally derived X chromosome in the extraembryonic membranes of the mouse. Nature 256: 640–642

Tamura K, Peterson D, Peterson N, Stecher G, Nei M, Kumar S (2011) MEGA5: molecular evolutionary genetics analysis using maximum likelihood, evolutionary distance, and maximum parsimony methods. Mol Biol Evol 28: 2731–2739

Tauber M, Hoybye C (2021) Endocrine disorders in Prader-Willi syndrome: a model to understand and treat hypothalamic dysfunction. Lancet Diabetes Endocrino. 9: 235– 246

Veyhl M, Keller T, Gorboulev V, Vernaleken A, Koepsell H (2006) RS1 (RSC1A1) regulates the exocytotic pathway of Na+-D-glucose cotransporter SGLT1. Am J Physiol Renal Physiol 291: F1213–1223

Wang H, Yang H, Shivalila C S, Dawlaty M M, Cheng A W, Zhang F, Jaenisch R (2013) One-step generation of mice carrying mutations in multiple genes by CRISPR/Cas-mediated genome engineering. Cell 153: 910–918

Whiteley AM, Prado MA, de Poot SAH, Paulo JA, Ashton M, Dominguez S, Weber M, Ngu H, Szpyt J, Jedrychowski MP et al (2021) Global proteomics of Ubqln2-based murine models of ALS. J Biol Chem 296:100153.

Xu B, Yang Z (2013) PAMLX: a graphical user interface for PAML. Mol Biol Evol 30: 2723–2724

Yagi T, Tokunaga T, Furuta Y, Nada S, Yoshida M, Tsukada T, Saga Y, Takeda N, Ikawa Y, Aizawa S (1993) A novel ES cell line, TT2, with high germline-differentiating potency. Anal Biochem 214: 70–76

Yeung RK, Xiang Z -H, Tsang S -Y, Li R, Ho TYC, Li Q, Hui C -K, Sham P -C, Qiao M -Q, Xue H (2018) Gabrb2-knockout mice displayed schizophrenia-like and comorbid phenotypes with interneuron-astrocyte-microglia dysregulation. Trans Psychiatry 8:128

Youngson NA, Kocialkowski S, Peel N, Ferguson-Smith AC (2005) A small family of sushi-class retrotransposon-derived genes in mammals and their relation to genomic imprinting. J Mol Evol 61: 481–490

Yousefvand S, Hamidi F (2020) Role of Paraventricular Nucleus in Regulation of Feeding Behaviour and the Design of Intranuclear Neuronal Pathway Communications. Int J Pept Res Ther 26: 1231–1242

Yuan H, Low C-M, Moody OA, Jenkins A, Traynelis SF (2015) Ionotropic GABA and Glutamate Receptor Mutations and Human Neurologic Diseases. Mol Pharmacol 88: 203–217

Zhang T, Li J, Yu H, Shi Y, Li Z, Wang L, Wang Z, Lu T, Wang L, Yue W et al (2018) Meta-analysis of GABRB2 polymorphisms and the risk of schizophrenia combined with GWAS data of the Han Chinese population and psychiatric genomics consortium. PLoS One 13: e0198690

Zhang X, Norton J, Carrière I, Ritchie K, Chaudieu I, Ryan J, Ancelin ML (2017) Preliminary evidence for a role of the adrenergic nervous system in generalized anxiety disorder. Sci Rep 7: 42676

Zhao J, Bao AM, Qi XR, Kamphuis W, Luchetti S, Lou JS, Swaab DF (2012) Gene expression of GABA and glutamate pathway markers in the prefrontal cortex of non-suicidal elderly depressed patients. J Affect Disord 138: 494–502

